# Automated, self-resistance gene-guided, and high-throughput genome mining of bioactive natural products from *Streptomyces*

**DOI:** 10.1101/2023.10.26.564101

**Authors:** Yujie Yuan, Chunshuai Huang, Nilmani Singh, Guanhua Xun, Huimin Zhao

**Affiliations:** Carl R. Woese Institute for Genomic Biology, University of Illinois at Urbana-Champaign, Urbana, IL 61801, USA; Departments of Chemistry, Biochemistry, and Bioengineering, University of Illinois at Urbana-Champaign, Urbana, IL 61801, USA; Department of Chemical and Biomolecular Engineering, University of Illinois at Urbana-Champaign, Urbana, IL 61801, USA

**Author notes:** These authors contributed equally: Yujie Yuan, Chunshuai Huang.

## Abstract

Natural products (NPs) produced by bacteria, fungi and plants are a major source of drug leads. *Streptomyces* species are particularly important in this regard as they produce numerous natural products with prominent bioactivities. Here we report a fully <u>a</u>utomated, <u>s</u>calable and high-throughput platform for discovery of bioactive <u>n</u>atural <u>p</u>roducts in *<u>S</u>treptomyces* (FAST-NPS). This platform comprises computational prediction and prioritization of target biosynthetic gene clusters (BGCs) guided by self-resistance genes, highly efficient and automated direct cloning and heterologous expression of BGCs, followed by high-throughput fermentation and product extraction from *Streptomyces* strains. As a proof of concept, we applied this platform to clone 105 BGCs ranging from 10 to 100 kb that contain potential self-resistance genes from 11 *Streptomyces* strains with a success rate of 95%. Heterologous expression of all successfully cloned BGCs in *Streptomyces lividans* TK24 led to the discovery of 23 natural products from 12 BGCs. We selected 5 of these 12 BGCs for further characterization and found each of them could produce at least one natural product with antibacterial and/or anti-tumor activity, which resulted in a total of 8 bioactive natural products. Overall, this work would greatly accelerate the discovery of bioactive natural products for biomedical and biotechnological applications.

**Graphic Abstracts:** 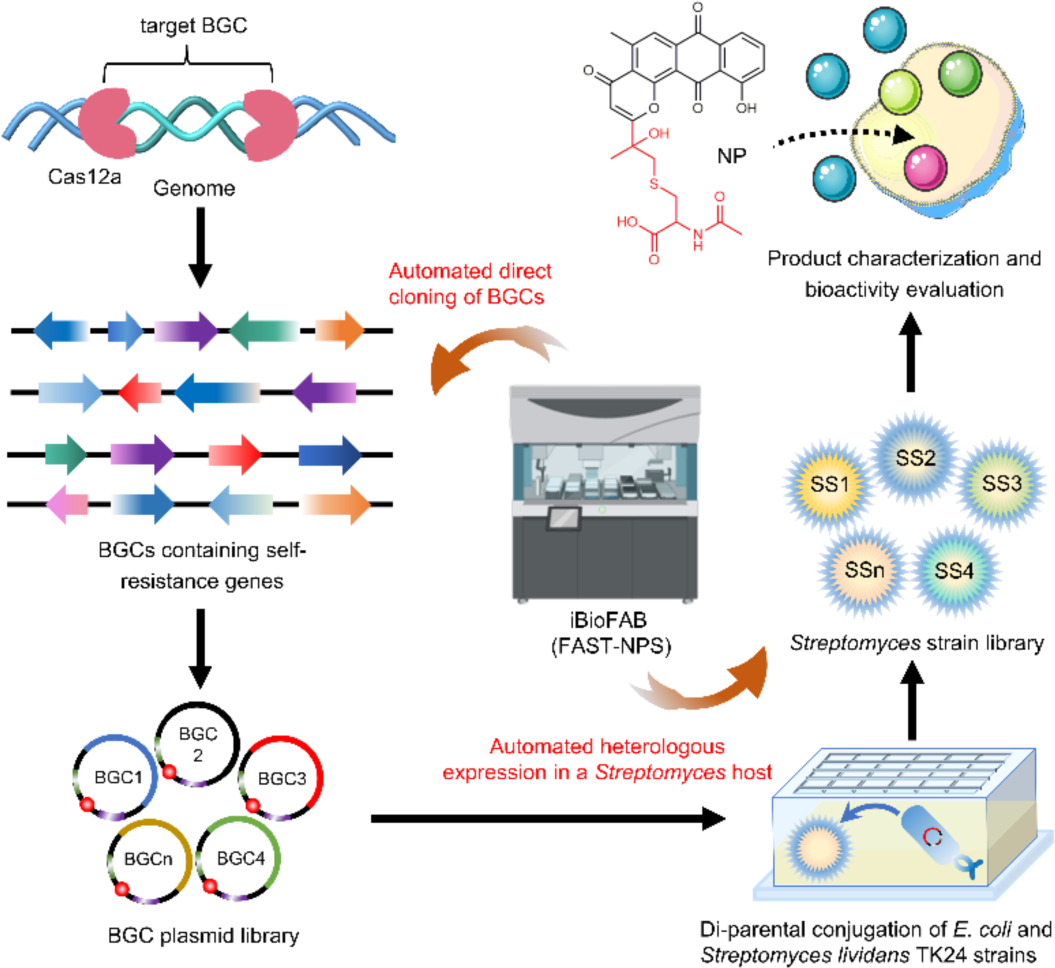

## Introduction

Natural products (NPs) produced by microorganisms and plants have been attractive in drug discovery because of their diversity in structures and biological functions^1^. *Streptomyces* species are of particular importance as they produce numerous natural products with bioactivities, such as erythromycin, daptomycin, and vancomycin^2, 3^. Recent advances in genome sequencing, bioinformatics, and machine learning tools have enabled the discovery of natural product biosynthetic gene clusters (BGCs) at an unprecedented rate^4–6^. However, only a limited number of NPs from these BGCs have been characterized, indicating a huge reservoir of unexploited natural products^7, 8^.

In the past decades, a wide variety of strategies have been developed to investigate uncharacterized BGCs and their corresponding natural products^9–11^. Prominent examples include native host-based BGC activation strategies such as genome editing by CRISPR/Cas9^12, 13^, use of elicitors^14^, and manipulation of global or pathway-specific regulators^15, 16^, and heterologous host-based strategies such as *Saccharomyces cerevisiae* based HEx (Heterologous expression) platform, and use of *Streptomyces* and *Aspergillus oryzae* chassis^8, 17, 18^. However, these strategies are low-throughput, time-consuming, and labor-intensive. To address these limitations, automated biofoundries have been developed to discover natural products in a high-throughput manner, including ribosomally synthesized and post-translationally modified peptides (RiPPs) and terpenoids in *Escherichia coli* and fungi, respectively^19, 20^. However, both studies were based on BGC refactoring strategies which are not suitable for large BGCs. Therefore, it is highly desirable to apply biofoundries to explore large BGCs for discovery of new natural products.

More importantly, as shown in numerous examples in literature, most of the natural products discovered by the above-mentioned strategies do not exhibit typical bioactivities such as antibacterial, antifungal, or antitumor activities, which indicates that they are either not biologically active or require further characterization by more specialized and sophisticated bioassays^12, 14, 17^. One effective strategy to address this challenge is to use self-resistance genes within the target BGCs to guide the discovery of bioactive natural products^21^. Self-resistance is an essential evolutionary response to the synthesis of bioactive natural products. To shield themselves from the detrimental impacts of their own natural products, one strategy the organisms utilize is to encode a functionally analogous self-resistance enzyme (SRE), which is a modified version of the housekeeping enzyme. The SRE exhibits a few mutations compared to the housekeeping enzyme, thereby hindering its interaction with toxic natural products while retaining its functional integrity^21–23^. The inclusion of the SRE within a BGC can act as a predictive indicator for the biological efficacy of the natural products synthesized through the pathway. The utilization of the self-resistance gene as a guiding principle has resulted in the identification of bioactive natural products, including aspterric acid, clipibicyclene, pekiskomycin, and thiotetroamide^22–25^. Hence, utilizing self-resistance mechanisms as a guide can increase the probability of discovering new bioactive natural products and their modes of action.

In this work, we report the development of a fully <u>a</u>utomated, <u>s</u>calable and high-throughput platform for discovery of bioactive <u>n</u>atural <u>p</u>roducts in *<u>S</u>treptomyces* (FAST-NPS). This platform seamlessly integrates the Antibiotic Resistant Target Seeker (ARTS) tool used to identify BGCs containing potential self-resistance genes^26^ with Illinois Biological Foundry for Advanced Biomanufacturing (iBioFAB)^27, 28^ used for automated direct cloning and heterologous expression of BGCs in a *Streptomyces* host (**Fig. 1**). We successfully adapted our previously developed direct cloning method named Cas12a-assisted precise targeted cloning using *in vivo* Cre-*lox* recombination (CAPTURE) on iBioFAB platform^29^. Unlike BGC refactoring strategies, direct cloning methods can capture microbial BGCs from genomic DNA, especially the large BGCs from high GC-content organisms including *Streptomyces*^29–32^. We demonstrate the capability of FAST-NPS by investigating 105 BGCs from 11 different *Streptomyces* strains, leading to the discovery of 23 natural products from 12 BGCs. Five of these 12 BGCs were selected for further characterization and each of them produced at least one compound with antibacterial and/or anti-tumor bioactivity, resulting in the discovery of 8 bioactive natural products.

**Fig. 1.**
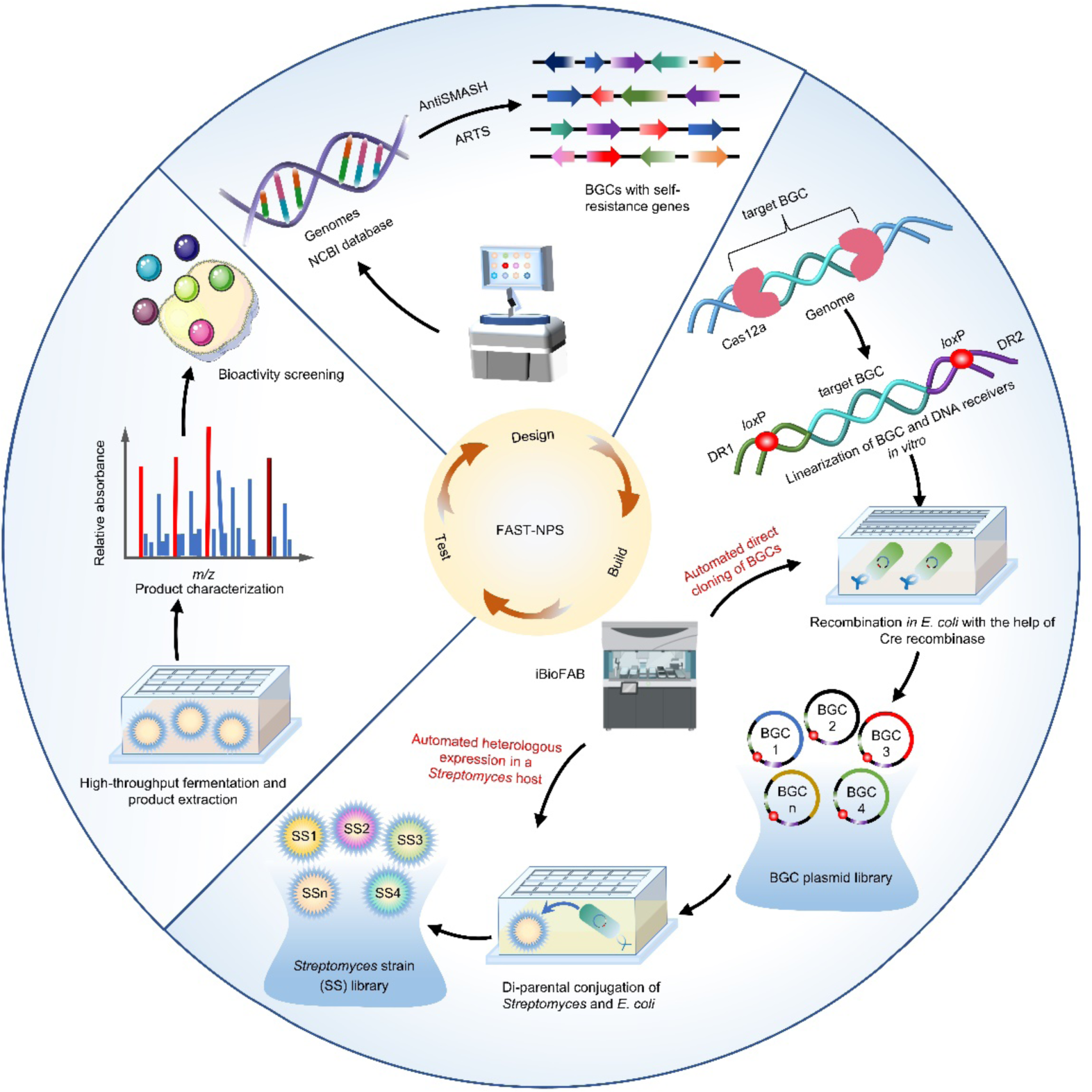
Overview of the FAST-NPS workflow for rapid discovery of bioactive natural products in a *Streptomyces* host. This workflow consists of three main modules, ranging from bioinformatic analysis of BGCs containing potential self-resistance genes (Design) to BGC plasmid library construction by automated direct cloning and heterologous expression of BGCs in a *Streptomyces* host (Build) to high-throughput fermentation of *Streptomyces* strains, high-throughput product extraction and structural and bioactivity characterization (Test). NCBI, National Center for Biotechnology Information; antiSMASH, Antibiotics and Secondary Metabolite Analysis Shell; ARTS, Antibiotic Resistant Target Seeker; BGC, Biosynthetic Gene Cluster; DR, DNA receiver fragment; iBioFAB, Illinois Biological Foundry for Advanced Biomanufacturing.

## Results

### Development of the FAST-NPS platform

The FAST-NPS platform consists of three main modules (**Fig. 1**): 1) In the Design module, we used the genome mining tools antiSMASH^33^ and ARTS^26^ to identify and prioritize putative BGCs that contain potential self-resistance genes. 2) In the Build module, we used iBioFAB to automate the workflow of CAPTURE-based direct cloning and heterologous expression of target BGCs in a *Streptomyces* host. iBioFAB is an integrated robotic system that enables rapid prototyping of biological systems^27^ (**Supplementary Fig. 1**). The automated CAPTURE workflow consists of PCR amplification of the DNA receiver (DR), PCR product recovery, guide RNA (gRNA) cassette transcription and recovery, genomic DNA (gDNA) digestion by FnCas12a enzyme, target BGC fragment recovery, linearization of DNA receivers and BGC fragments, and plasmids extraction. The automated heterologous expression workflow consists of transformation of *E. coli* and di-parental (*E. coli*-*Streptomyces*) conjugation. In the automated workflow, the Momentum^TM^ Workflow Scheduling software was utilized to control the robotic arm, communicate with devices and execute user commands. The automated pipeline was designed by first identifying well-defined unit operations, such as liquid handling, centrifugation, incubation, and temperature control, which were then orchestrated by corresponding instruments on the iBioFAB. The location of the samples and the routes for their transportation between the different unit operations were precisely determined and synchronized with the corresponding instruments using the Momentum software (**Supplementary Fig. 1**). 3) In the Test module, we developed an automated workflow for high-throughput fermentation and product extraction using *Streptomyces* in 24 deep-well plates. After the extracts detection by high-resolution-electrospray ionization-mass spectrometry (HR– ESI–MS), the target natural products were isolated, characterized, and their bioactivities were evaluated.

### Identification of target BGCs containing potential self-resistance genes

We first searched the sequenced and annotated *Streptomyces* genomes deposited in the National Center for Biotechnology Information (NCBI) and selected 11 newly sequenced *Streptomyces* strains, namely *Streptomyces aureoverticillatus* ISP 5080, *Streptomyces bikiniensis* ISP 5582, *Streptomyces rubradiris* NRRL 3061, *Streptomyces echinatus* ISP 5013, *Streptomyces thermoviolaceus* NRRL B-12374, *Streptomyces violascens* NRRL B-2700, *Streptomyces nojiriensis* NRRL B-16930, *Streptomyces spororaveus* NRRL B-16378, *Streptomyces nitrosporeus* ATCC12769, *Streptomyces* sp. NRRL F-5635, and *Streptomyces monomycini* NRRL B-24309. Subsequently, we used antiSMASH (v6.0)^33^, one of the most popular genome mining tools, to predict the BGCs in these strains (**Fig. 2a**).

**Fig 2.**
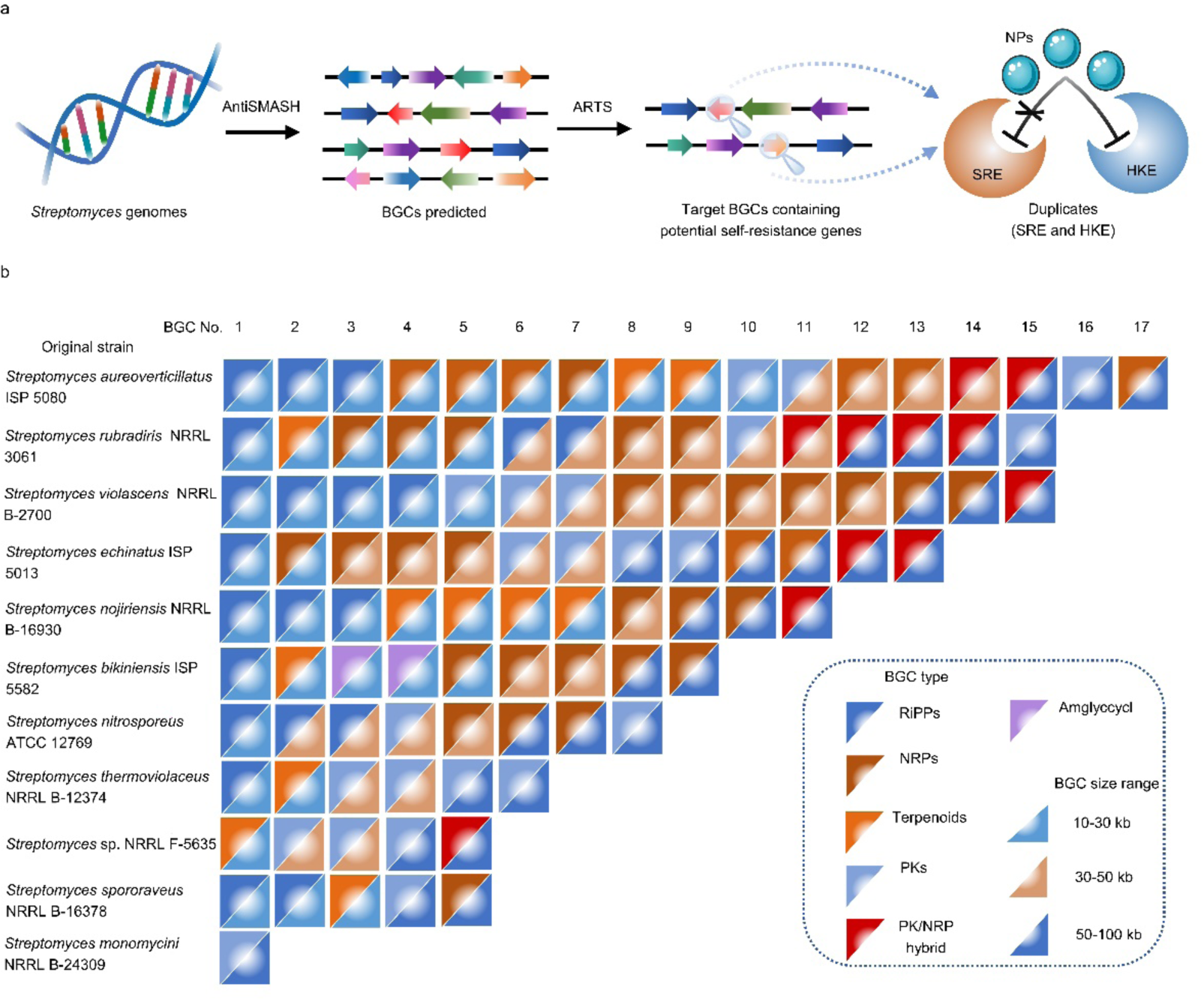
Bioinformatic analysis of BGCs containing potential self-resistance genes. **a.** Discovery of target BGCs guided by self-resistance genes. A self-resistance enzyme (SRE) is a modified version of the housekeeping enzyme (HKE). The SRE exhibits a few mutations compared to the HKE, thereby hindering its interaction with toxic natural products while retaining its functional integrity. **b**. Target BGCs selected from various *Streptomyces* strains. The 105 BGCs ranging from 10 kb to 100 kb covered 6 different classes of natural products. SRE, self-resistance enzyme; HKE, housekeeping enzyme; RiPPs, ribosomally synthesized and post-translationally modified peptides; NRPs, non-ribosomally synthesized peptides; PKs, polyketides.

Next, we used ARTS2.0 to further analyze the BGCs containing potential self-resistance genes. ARTS utilizes the antiSMASH platform to identify BGCs, which are then analyzed to identify core genes (those present in a majority of strains from a given microbial family) or known resistance genes. These genes are assigned a “proximity score” to facilitate efficient genome mining for antibiotics with interesting and novel targets by linking housekeeping and known resistance genes to BGC proximity, duplication, and horizontal gene transfer (HGT) events. ARTS detects possible resistant housekeeping genes based on three criteria: duplication (including small modifications), localization within a BGC, and evidence of HGT^26^ (**Fig. 2a**). On the basis of ARTS analysis, 105 BGCs containing known resistance genes or duplicated housekeeping genes from 11 *Streptomyces* strains were selected as the candidates for characterization (**Fig. 2b**, **Supplementary Table 1**). In the meantime, to demonstrate the generality of the FAST-NPS platform, the 105 BGCs covered 6 different classes of natural product families including RiPPs, non-ribosomally synthesized peptides (NRPs), terpenoids, polyketides (PKs), hybrid of PKs/NRPs and amglyccycl, and their sizes ranged from 10 kb to 100 kb (**Fig. 2b**, **Supplementary Table 1**).

### Automated direct cloning of target BGCs

We previously developed CAPTURE for direct cloning of large genomic fragments (>100 kb) with high efficiency^29^. In this study, we adapted this method to an automated, high-throughput workflow for direct cloning of target BGCs on the iBioFAB. This workflow consists of five major steps: (1) Preparation of DNA receiver (DR) fragments using PCR and DNA purification; (2) Preparation of guide RNAs using *in vitro* transcription; (3) Digestion of purified genomic DNA by FnCas12a enzyme *in vitro*; (4)

Linearization of DR fragments and digested target BGC DNAs in a T4 DNA polymerase exo + fill in DNA assembly reaction; and (5) Site-specific recombination of assembled linear DNA in *E. coli* by Cre-*lox*P system followed by BGC plasmid purification and confirmation (**Fig. 3a**). A script was created for Momentum to execute this workflow. The comprehensive experimental procedures, along with the inputs and outputs between each step, and the equipment utilized, were thoroughly documented and presented in **Supplementary Fig. 2**.

**Fig 3.**
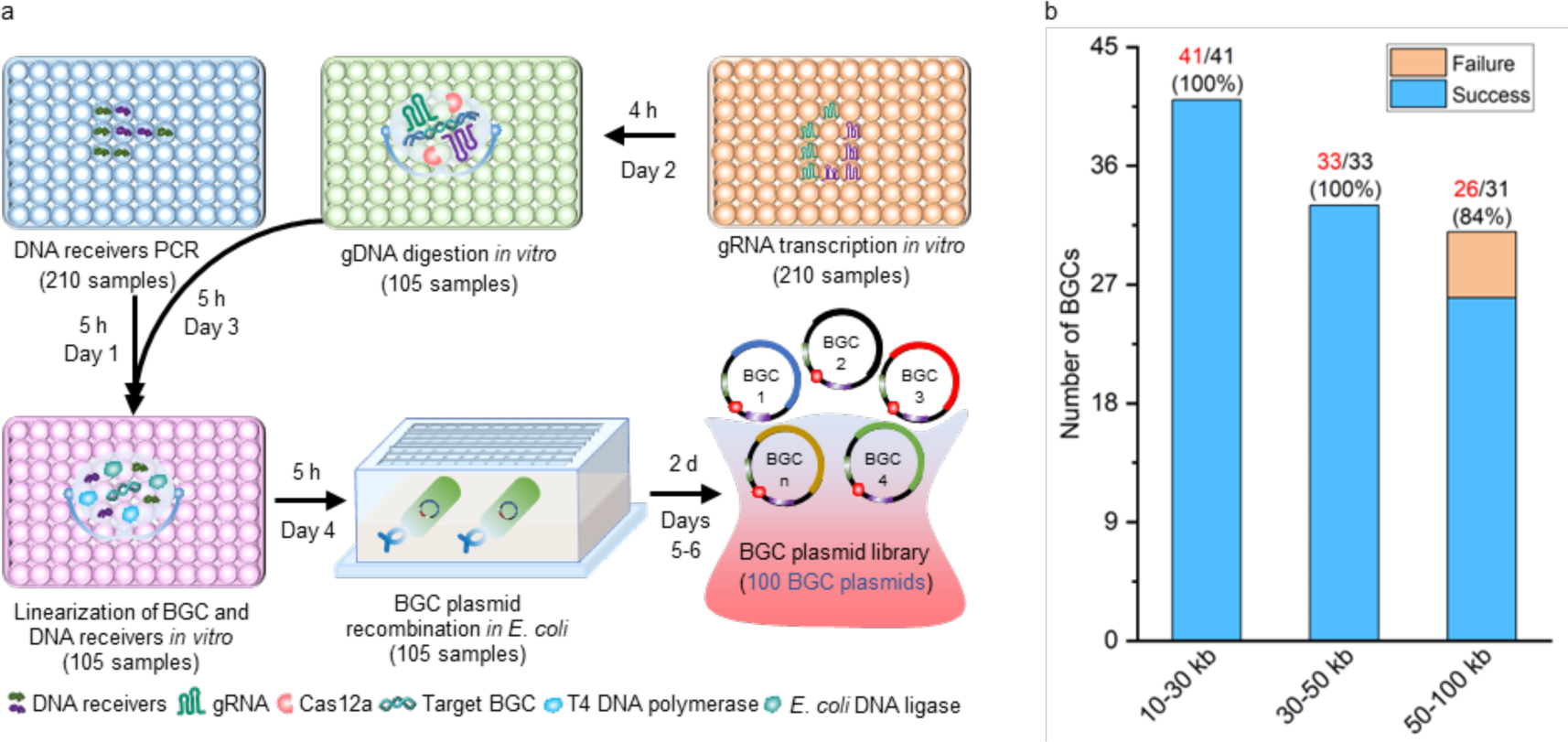
Automated direct cloning of target BGCs using iBioFAB. **a**, Schematic overview of a high- throughput BGC direct cloning workflow. There are five major steps in the workflow: step 1, preparation of DNA receivers using PCR; step 2, preparation of gRNAs using *in vitro* transcription; step 3, digestion of gDNAs by FnCas12a *in vitro*; step 4, linear assembly of DNA receivers and BGC fragments *in vitro*; step 5, linear DNA fragments circularization in *E. coli* followed by BGC plasmid purification and confirmation. The number in brackets represents the total samples in each step. **b**, Success rate of BGC direct cloning. The results show the cloning success rates among target BGCs with sizes ranging from 10 kb to 100 kb.

The schematic overview of these five steps was shown in **Supplementary Fig. 3**. Firstly, we designed and ordered DR primers carrying a *lox*P site on one end and a 39 bp DNA sequence sharing identity to the target BGC on the other end, and gRNA guides based on previous design principles for each target BGC^29^. Following that, the Tecan FluentControl system was employed to carry out a series of liquid handling steps, wherein the DR primers were diluted to a concentration of 5 μM. The gRNAs were prepared at a concentration of 25 μM for step 2 to set up the transcription by Fluent. During steps 1-4, the master mix, consisting of DNA polymerase for PCR amplification, T7 RNA polymerase for gRNA transcription, FnCas12a reaction system for gDNA digestion and T4 DNA polymerase for linearization of DNA fragments, were pre-prepared and aliquoted into destination plates by flexible-channel arm (FCA) tips. The additional necessary components for each step were then added by Fluent. The destination plates containing all components were then transferred to PCR thermocyclers to perform the reactions. After obtaining the assembled linear DNA products of DR and BGC fragments generated by step 4, the circularization of linear DNA by transformation into *E. coli* (step 5) was processed through the FluentControl system. After agar plating and colony picking, the *E. coli* strains containing corresponding BGCs were cultured overnight in the Cytomat incubator to facilitate plasmids extraction. The purified plasmids were subsequently subjected to digestion using specific restriction enzymes. Overall, this automated workflow can construct many BGC plasmids (in this study, 105) in parallel within six days. It is worth mentioning that the total duration of six days encompassed time-consuming steps such as cell inoculation and overnight culture. If we only consider the time spent on each step per day, an average of approximately five hours per day is required which minimizes the labor-intensive and time-consuming aspects of the automated workflow (**Fig. 3a**).

Using this automated BGC direct cloning workflow, we obtained 210 DR fragments (∼100% of the designed sequences) and 210 high-yield gRNAs, along with 105 purified BGC fragments and 105 assembled linear DNA fragments. The linear DNA fragments containing corresponding BGC parts were then circularized in *E. coli* harboring a helper plasmid expressing phage lambda Red Gam protein and Cre recombinase^29^. After confirmation by restriction enzymes, 100 out of 105 BGC plasmids were successfully cloned (∼95% success rate) (**Fig. 3b**, **Supplementary Fig. 4**, **Supplementary Table 3**). BGCs with sizes ranging from 10 kb to 50 kb were cloned with a ∼100% success rate while BGCs with sizes larger than 50 kb exhibited a ∼84% success rate (**Fig. 3b**). All incorrect constructs were determined to be false positives without BGC fragment insertion. This is likely due to the fact that the integrity of larger BGC fragments was disrupted because the preservation of gDNA integrity is crucial for the successful cloning of large BGCs. Therefore, to mitigate any potential damage to gDNA, particularly during the mixing steps, the lowest mixing speed is employed throughout the experimental procedures.

### Automated heterologous expression of BGCs in *Streptomyces*

Conjugation transfer was used to introduce heterologous genes into *Streptomyces*^34^ and the *E. coli*- *Streptomyces* di-parental intergenic conjugation method represents a commonly employed strategy in genetic engineering of *Streptomyces*^35, 36^. However, to our knowledge, no automated workflow for this conjugation process has been reported thus far. Therefore, to characterize BGCs in *Streptomyces* hosts on a large scale, we sought to develop an automated high-throughput workflow for heterologous expression of target BGCs in *Streptomyces*. This workflow comprises two major steps: high-throughput transformation of BGC plasmids into *E. coli* WM6026 and conjugation of *E. coli*-*Streptomyces* (**Fig. 4a**, **Supplementary Fig. 5**). For the first step, the chemical competent *E. coli* WM6026 cells were prepared and aliquoted into cold 96-well plates followed by heat shock transformation using heat blocks on Fluent deck. After agar plating on 8-well plates and colony picking, the *E. coli* WM6026 strains containing corresponding BGCs were cultured overnight in the Cytomat incubator. In the following day, the seed cultures were inoculated into fresh medium in 24 deep-well plates to generate BGC donor strains. In the meantime, the *Streptomyces lividans* TK24 spores were collected as the receiver strains and mixed with *E. coli* donors by using multi-channel arm (MCA) tips. The exconjugants were screened on 8-well agar plates supplemented with antibiotics. A total of 100 *Streptomyces* strains (SSs) harboring the corresponding BGCs were constructed within one week (**Fig. 4a**, **Supplementary Table 4**).

**Fig 4.**
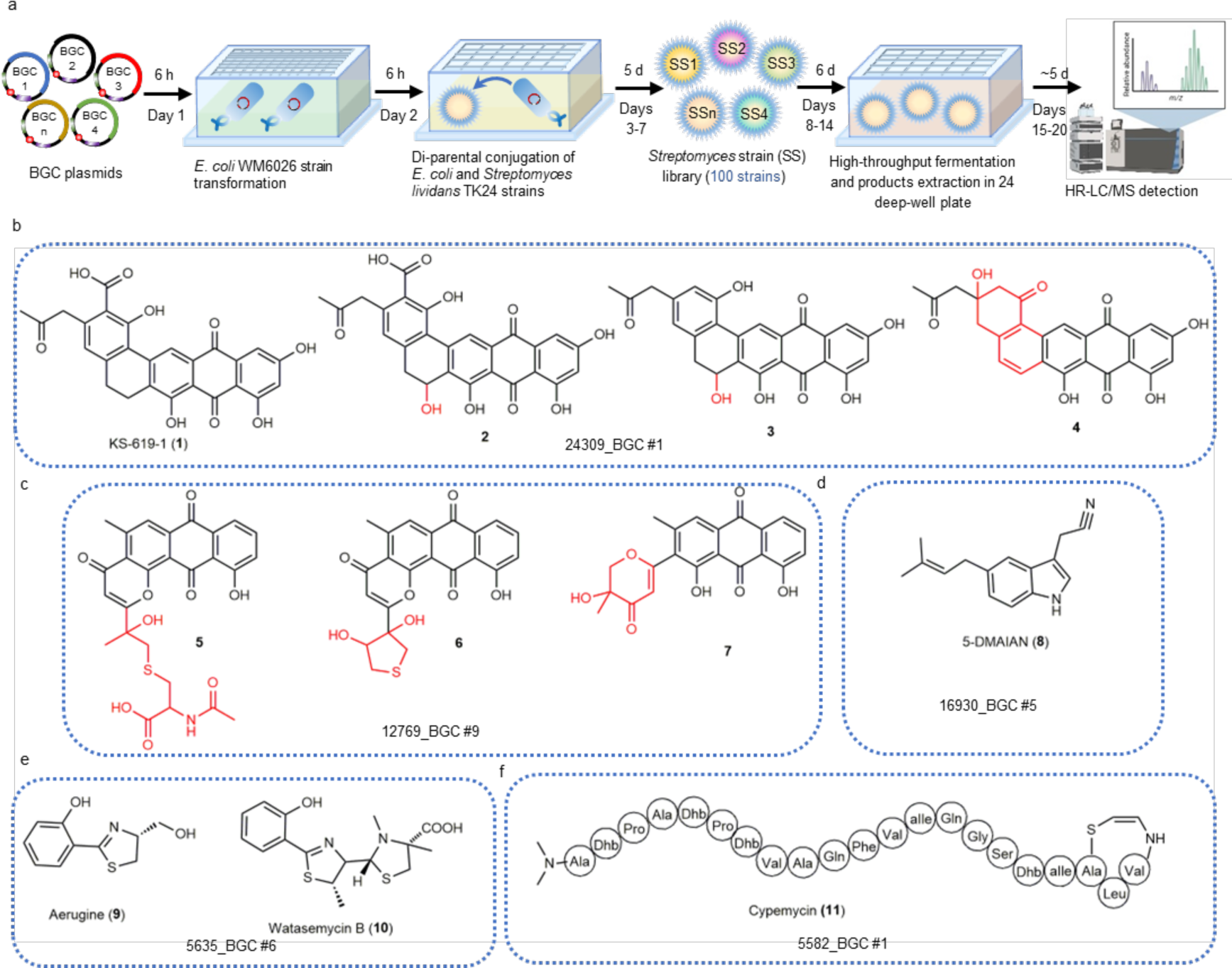
Automated heterologous expression of BGCs in *Streptomyces*. **a**, Schematic workflow for high-throughput di-parental conjugation, fermentation and product isolation in *Streptomyces*. BGC plasmids were transformed into *E. coli* WM6026 to generate donor strains. The di-parental conjugation method was adopted to introduce BGC DNA into *S. lividans* TK24 host to construct a *Streptomyces* strain (SS) library. The *Streptomyces* strains were fermented and extracted in 24 deep-well plates. The crude extracts were detected using HR-ESI-MS equipped with a HPLC system. **b**, Chemical structures of characterized compounds (**1**−**4**) identified from *Streptomyces monomycini* NRRL B-24309_BGC#1. **c**, Chemical structures of characterized compounds (**5**−**7**) identified from *Streptomyces nitrosporeus* ATCC 12769_BGC#9. **d**, Chemical structure of 5-DMAIAN (**8**) identified from *Streptomyces nojiriensis* NRRL B-16930_BGC#5. **e**, Chemical structures of aerugine and watasemycin B (**9**-**10**) identified from *Streptomyces* sp. NRRL F-5635_BGC#6. **f**, Chemical structure of cypemycin (**11**) identified from *Streptomyces bikiniensis* ISP 5582_BGC#1.

Next, we developed a high-throughput fermentation and product isolation workflow for *Streptomyces* strains (**Fig. 4a**, **Supplementary Fig. 5**). *Streptomyces* strains harboring target BGCs were inoculated into 24 deep-well plates and cultured for around six days. Afterwards, natural products were extracted from the cultures and detected using HR-ESI-MS equipped with a high-performance liquid chromatography (HPLC) system. A total of 23 (**1**-**23**) natural products from 12 BGCs were detected (**Figs. 4b-f**, **Supplementary Fig. 6**, **Supplementary Table 5**). To get a better understanding of the chemical diversity generated by our FAST-NPS platform, we characterized a representative subset of compounds from five different BGC strains by HR-ESI-MS and nuclear magnetic resonance (NMR). The 24309_BGC#1 from *S. monomycini* NRRL B-24309 encoded a putative type II polyketide synthase (T2PKS) pathway. HPLC analysis of the metabolite profile led to the discovery of one anthraquinone-type compound KS-619-1 (**1**) that was previously reported^37^ and its three new derivatives, designated as grianquinones A−C (**2**−**4**) (**Fig. 4b**, **Supplementary Figs. 7-26**, **Supplementary Tables 6-7**). The 12769_BGC#9 from *S. nitrosporeus* ATCC 12769 encoded another putative T2PKS pathway, which led to the discovery of three novel anthrapyrones A−C (**5**−**7**) through HPLC analysis of the metabolite profile (**Fig. 4c**, **Supplementary Figs. 27-44**, **Supplementary Tables 8-9**). Further metabolic profiling analyses of the other positive BGCs resulted in the isolation of one terpenoid compound 5-DMAIAN (**8**), two analogs of thiazostatin aerugine and watasemycin B (**9** and **10**) and a linaridin RiPP, cypemycin (**11**) (**Figs. 4d**-**f**, **Supplementary Figs. 45-51**)^38–41^. The remaining 12 uncharacterized natural products (**12**−**23**) were presented with HR-ESI-MS data, which were collected and analyzed by Qual Browser Thermo Xcalibur (**Supplementary Fig. 6**).

### Bioactivity evaluation of characterized natural products

The strategy of using self-resistance genes to discover bioactive natural products has been demonstrated in a few case studies^22–25^. In this strategy, certain self-resistance elements that were found to co-localize within corresponding BGCs, such as ATP-binding cassette (ABC) transporter, DNA gyrase, beta-lactamase, and acyl-CoA carboxylase, have been functionally validated^21, 42^. Under the guidance of self-resistance genes, we conducted a thorough investigation of the five BGCs responsible for producing natural products **1**−**11**. Within the 24309_BGC#1, we discovered the presence of an acyl-CoA carboxylase self-resistance model gene within the BGC. Similarly, the 12769_BGC#9 contained a metallo-hydrolase self-resistance model gene in close proximity to the BGC. Furthermore, within the 16930_BGC#5, we identified a beta-lactamase self-resistance model gene within the BGC. Additionally, both the 5635_BGC#6 and the 5582_BGC#1 contained an ABC efflux pump transporter self-resistance model gene (**Supplementary Table 1**). These findings suggest that the characterized natural products **1**−**11** potentially possess specific bioactivities, which prompted us to perform bioassays.

As summarized in **Table 1**, we analyzed the antimicrobial and anti-tumor activities of compounds **1**−**10** against various Gram-positive bacteria strains and four commonly used human tumor cell lines. Among the compounds **1**−**4** produced by 24309_BGC#1, **1**, **3**, and **4** exhibited antibacterial properties against five distinct Gram-positive bacteria, including methicillin-resistant *Staphylococcus aureus* USA300 (MRSA) with minimal inhibitory concentration (MIC) ranging from 12.5 to 100 μg/mL. Moreover, **3** and **4** demonstrated anti-tumor activity against two human cell lines: human colon carcinoma (HCT- 116) with the half maximal inhibitory concentration (IC_50_) of 84.12 ± 3.08 μg/mL and 23.91 ± 0.67 μg/mL, respectively, as well as non-small cell lung cancer cells (A549) with IC_50_ values of 74.29 ± 5.93 μg/mL and 45.49 ± 6.01 μg/mL, respectively. Regarding the compounds **5**−**7** derived from 12769_BGC#9, we observed that the compounds, **5** and **7**, displayed anti-tumor activity against A549 cell lines, with IC_50_ values of 29.62 ± 2.73 μg/mL and 55.63 ± 8.76 μg/mL, respectively. Similarity, the 5-DMAIAN (**8**) identified from 16930_BGC#5, exhibited anti-tumor activity against A549 cell lines with an IC_50_ value of 59.90 ± 4.49 μg/mL. Furthermore, for the compounds **9** and **10** produced by the 5635_BGC#6, we discovered that the compound watasemycin B (**10**) demonstrated significant bioactivity against six Gram-positive bacteria with MIC values ranging from 3.1 μg/mL to 50 μg/mL. Notably, its antibacterial efficacy against to *Bacillus subtilis* ATCC6633 was consistent with a previous study^40^. Additionally, watasemycin B also exhibited potent anti-tumor bioactivity against HCT-116 cell lines with an IC_50_ value of 13.87 ± 6.78 μg/mL. Regarding to the compounds **6** and cypemycin (**11**), we were unable to evaluate their bioactivity due to the limited quantity available. Nevertheless, previous studies have reported diverse bioactivities of cypemycin, ranging from antimicrobial effects against methicillin-resistant *Staphylococcus aureus* (MRSA) to anti-cancer activity against mouse leukemia cells^41, 43^. For the compounds **2** from 24309_BGC#1 and **9** from 5635_BGC#6, we did not observe distinct bioactivities (MIC > 100 μg/mL, IC_50_ > 100 μg/mL). One possible reason for this could be that the compounds are intermediates lacking the essential core pharmacophore. Overall, the compounds identified from the five BGCs guided by self-resistance genes exhibited antibacterial and/or anti-tumor activities, demonstrating the effectiveness of utilizing self-resistance mechanisms as a guiding principle to increase the probability of discovering bioactive natural products and establishing their bioactivities.

**Table 1.** Bioactivity evaluation of characterized natural products.

|  | MIC (μg/mL) |  |  |  |  |  |  |
| --- | --- | --- | --- | --- | --- | --- | --- |
| Gram-positive bacteria | 1 | 3 | 4 | 5 | 7 | 8 | 10 |
| *Bacillus halodurans* C-125 | 12.5 | 50.0 | 100.0 | >100 | >100 | >100 | 3.1 |
| *Bacillus anthracis* str. Sterne | 25.0 | 50.0 | 100.0 | >100 | >100 | >100 | 3.1 |
| *Bacillus cereus* TZ417 | 50.0 | 50.0 | >100 | >100 | >100 | >100 | 3.1 |
| *Bacillus subtilis* ATCC6633 | >100 | >100 | >100 | >100 | >100 | >100 | 50.0 |
| *Enterococcus faecium* U503 | >100 | >100 | >100 | >100 | >100 | >100 | >100 |
| *Lactococcus lactis* CNRZ 481 | >100 | >100 | >100 | >100 | >100 | >100 | >100 |
| *Micrococcus leteus* ATCC 4698 | >100 | >100 | >100 | >100 | >100 | >100 | >100 |
| *Staphylococcus aureus* USA300 | 50.0 | 50.0 | 100.0 | >100 | >100 | >100 | 12.5 |
| *Staphylococcus epidermidis* 15X154 | 100.0 | 100.0 | 100.0 | >100 | >100 | >100 | 25.0 |
| *Streptococcus mutans* ATCC 25175 | >100 | >100 | >100 | >100 | >100 | >100 | >100 |
|  | IC<sub>50</sub> (μg/mL) |  |  |  |  |  |  |
| Human tumor cell lines | 1 | 3 | 4 | 5 | 7 | 8 | 10 |
| HCT-116 (human colon carcinoma) | >100 | 84.12±3.08 | 23.91±0.67 | >100 | >100 | >100 | 13.87±6.78 |
| Hela (human cervical carcinoma) | >100 | >100 | >100 | >100 | >100 | >100 | >100 |
| A549 (Non-small cell lung cancer cells) | >100 | 74.29±5.93 | 45.49±6.01 | 29.62±2.73 | 55.63±8.76 | 59.90±4.49 | >100 |
| HepG2 (human liver carcinoma) | >100 | >100 | >100 | >100 | >100 | >100 | >100 |

## Discussion

The proliferation of sequenced microbial genomes has led to numerous uncharacterized natural product BGCs. However, a significant challenge persists in identifying and elucidating the structures of the compounds associated with these BGCs. The most efficient strategy for studying these uncharacterized BGCs and discovering their associated compounds is through heterologous expression^18^. Despite numerous attempts to develop universal high-throughput expression platforms, heterologous expression is currently limited to refactoring small BGCs and does not cater specifically for *Streptomyces* strains^19,20^. To address this limitation, we developed an automated, scalable, and high-throughput platform specifically designed to facilitate the discovery of natural products from *Streptomyces* genus harboring large BGCs with high GC-content. This FAST-NPS platform incorporated the guidance of self- resistance genes that are co-located within the corresponding BGCs, which significantly increased the chance of discovering bioactive natural products. Using this platform, we cloned 100 out of 105 BGCs ranging from 10 kb to 100 kb with a success rate of 95% and discovered 23 natural products derived from 12 BGCs in a much reduced timeline compared to traditional approaches. (**Fig. 3a**, **Fig. 4a**). We selected five of these 12 BGCs for further characterization and found that each of them produced at least one natural product (a total of 8) showing antibacterial and/or anti-tumor activities.

Despite our high success rate of discovering new bioactive compounds, it is important to acknowledge the limitations of our platform. First, the FAST-NPS platform does not encompass genomic DNA extraction steps. This means that for the direct cloning process, substantial amounts of purified genomic DNA (at least 15 μg of each BGC) are necessary. The preparation of high-quality genomic DNA typically involves the use of toxic reagents such as phenol/chloroform, which is challenging to implement on the biofoundry. Second, the workflow of the FAST-NPS platform does not extend to the isolation and structural characterization of the discovered compounds. In our case, the structures of over half of the natural products discovered in this study remain unknown at present. Third, the *Streptomyces* strains associated with the majority of BGCs (>80%) did not produce any detectable amounts of natural products. This observation can be attributed to two factors: 1) It is possible that evolutionarily dead core biosynthetic genes within the BGCs may have rendered them non-functional. Gaining deeper insights into the core biosynthetic enzymes might facilitate more robust selections of active BGCs in the future^20^; 2) Expression in heterologous host poses significant challenges, such as insufficient or minimal transcription of essential biosynthetic genes and the absence of necessary precursors or cofactors for certain enzymes^29^. Strategies such as replacing native promoters with strong promoters upstream of core biosynthetic genes, overexpression of pathway-specific activators, deletion of pathway-specific repressors, transcription factor decoys, and the development of more suitable chassis containing essential precursor encoding systems, may hold promise for mitigating these challenges^2, 13, 44, 45,46^.

One prominent application of self-resistance genes is the assignment of a mode of action to bioactive compounds^22, 23, 25^. In our study, we observed the presence of self-resistance model enzymes, acyl-CoA carboxylase, metallo-hydrolase, and beta-lactamase, in the 24309_BGC#1, 12769_BGC#9 and 16930_BGC#5, respectively. Gaining deeper insights into the functionality and characteristics of these enzymes could potentially aid in deciphering the mode of action of the bioactive natural products produced by these corresponding BGCs in future investigations. However, it should be noted that despite the guidance provided by self-resistance genes in BGC prioritization, our current limited knowledge on natural product biosynthesis poses certain challenges. For instance, the relatively low occurrence rate of self-resistance enzymes in the currently sequenced *Streptomyces* genomes (<5%) means that predictions of self-resistance genes in BGCs may sometimes be inaccurate, resulting in both false positives and false negatives^42^. Several scenarios illustrate the complexities involved: 1) a housekeeping enzyme homologue may coincidently be duplicated within the BGC of a natural product and may not actually serve as a SRE; 2) a variant of a housekeeping enzyme within a BGC may indeed function as a bona fide biosynthetic enzyme involved in the assembly of natural products rather than as a SRE; 3) a SRE may be co-located within the BGC but duplicated without any associated housekeeping enzyme^47–49^. To improve the accuracy of BGC prioritization, it is crucial to expand our knowledge on biosynthetic mechanisms and enzymology and gain a deeper understanding of self-resistance mechanisms. The accumulation of such knowledge in the future could pave the way for the development of computational prediction tools that prioritize BGCs with greater precision and reliability. This would ultimately streamline the process of natural product discovery and enhance our ability to unlock the full potential of BGCs.

In summary, our FAST-NPS platform enables highly efficient discovery of bioactive natural products produced by *Streptomyces* by integrating automated and high-throughput direct cloning and heterologous expression of BGCs with the use of self-resistance genes for BGC prioritization. This platform can be easily adapted for the investigation of other natural products and microorganisms, which holds immense potential in accelerating the discovery and development of new pharmaceutical agents with diverse therapeutic applications.

## Methods

### Materials and reagents

Polymerase chain reaction (PCR) amplification of DNA receivers for plasmids construction and colony PCR were performed using PrimeSTAR^®^ Max DNA Polymerase (R045B, TaKaRa, Kyoto, Japan). PCR primers were synthesized by Integrated DNA Technologies (IDT, Coralville, IA, USA). UltraPure Phenol: Chloroform: Isoamyl Alcohol (15593-049), Chloroform (C298-500), Proteinase K (fungal) (25530-015), isopropanol (A416-1) for genomic DNA (gDNA) extraction, and RNAse A (EN0531) for gDNA digestion were purchased from Thermo Fisher Scientific (Indianapolis, IN, USA). RNase A (R-050-500) for gDNA extraction, apramycin (A-600-10), isopropyl β-D-1-thiogalactopyranoside (IPTG) (I2481), and 5-bromo-4-chloro-3-indolyl-β-D-galactoside (X-GAL) (X4281C10) for blue-white screening were purchased from Gold Biotechnology (Olivette, Missouri). Lysozyme (L6876-10G), mutanolysin (M9901-10KU), and achromopeptidase (A3547- 100KU) for gDNA extraction were purchased from Sigma-Aldrich (Louis, MO, USA). HiScribe™ T7 High Yield RNA Synthesis Kit (E2040S), NAD^+^ (B9007S), dNTPs (N0447S), T4 DNA polymerase (M0203L), *E. coli* DNA ligase (M0205L), proteinase K (P8107S) for gDNA digestion, and restriction enzymes for BGC plasmid verification were purchased from New England Biolabs (NEB, Ipswich, MA, USA). ZR-96 DNA Clean-Up Kit (D4018) for DNA receiver PCR product purification and Quick-RNA 96 Kit for guide RNAs (gRNAs) purification were purchased from Zymo Research (Irvine, CA, USA). Invitrogen™ PureLink™ Pro Quick96 Plasmid Purification Kit (K211004A) was purchased from Thermo Fisher Scientific. Cloning was performed in *E. coli* strain NEB 10β (NEB). L-arabinose (A3256-25 g), 2,6-diaminopimelic acid (DAP) (D1377-5G), and nalidixic acid (N8878-25G) were purchased from Sigma-Aldrich (Louis, MO, USA). *Streptomyces lividans* TK24 was used for heterologous expression of natural product BGCs cloned from Actinomycete strains. Tryptone and yeast extract for medium preparation were purchased from Thermo Fisher Scientific. Malt Extract (M41600- 500.0) and D-maltose (M22000-500.0) were purchased from Research Products International (Mount Prospect, IL, USA). All salts and organic reagents for product extraction and detection were purchased from Thermo Fisher Scientific.

### Preparation of DNA receiver fragments

The primers were designed as previously reported^29^. The primers were synthesized into 96-well IDT plates and diluted into 25 μM using RNA-free water when the orders were received from IDT. The primers were further diluted to 5 μM and aliquoted into new 96-well PCR plates using Multi-Channel Arm (MCA) inside FluentControl 1080 liquid handler (Tecan, Männedorf, Switzerland) for sample preparation of PCR reactions. The DNA polymerase master mix containing templates (pBE44, pBE48, or pBE45; 1 ng of plasmid DNA of each sample) was prepared in 96 deep-well plates and aliquoted into 96-well PCR plates with 35 μL of each well by Flexible-Channel Arm (FCA) for sample preparation of PCR reactions. 5 μL of primers were then transferred into a destination plate using MCA and mixed 8 times. Then, the destination plate was transferred into PCR machine to start PCR reaction. After checking by automated parallel capillary gel electrophoresis using Fragment Analyzer (Agilent Technologies, Santa Clara, CA), the PCR products were digested by *Dpn*I and purified using ZR-96 DNA Clean-Up Kit according to the manufacturer’s protocol. The detailed procedures were as presented as the step 1 in Supplementary Fig. 3. All primers used in this study were listed in Supplementary Table 2.

### Guide RNA (gRNA) transcription and purification *in vitro*

The procedures included four steps as follows: 1) Hybridization of gRNA oligonucleotides: 12 μL of master mix containing 2 μL of 10 × NEBuffer 3.1 were aliquoted into 96-well PCR plates using FCA for sample preparation. Subsequently, 4 μL of oligonucleotide-Forward (25 μM) and 4 μL of oligonucleotide-Reverse (25 μM) template ssDNA were added. The mixture was incubated at 98 °С for 5 min and the temperature was then slowly lowered with the rate of 0.1 °C/s until 10 °C was reached. 2) gRNA transcription *in vitro*: 26 μL of master mix containing 2 μL of T7 RNA polymerase and 10 μL of NTP mix (10 × buffer, ATP, UTP, CTP, GTP) were aliquoted into 96-well PCR plates for sample preparation. 4 μL of hybridized mixture from step 1 plate was added as template for *in vitro* RNA transcription following the manufacturer’s protocol for transcription of short RNA molecules provided by HiScribe T7 quick high yield RNA Synthesis kit. 3) DNase I treatment: 20 μL of DNase I solution containing 5 μL of DNase I provided by Quick-RNA 96 Kit was added into step 2 plate which was then incubated at 37 °С for 15 min. 4) gRNA purified: the gRNAs were finally purified using Quick-RNA 96 Kit following optimized manufacturer’s protocol. For the elution step, 70 μL of DNase/RNase free water was used instead of 15 μL. The detailed procedures were as presented as the Step 2 in Supplementary Fig. 3. All gRNA oligonucleotide ssDNA used in this study were listed in Supplementary Table 2.

### Genomic DNA digestion and purification

The isolation of genomic DNA (gDNA) and purification of FnCas12a enzyme were performed following the protocols described elsewhere^29^. FnCas12a digestion of genomic DNA was performed in a 150 μL reaction system in a 96-well PCR destination plate containing 20 μg of purified genomic DNA, ∼4 μg of each gRNA, 15 μg of FnCas12a enzyme and 15 μL of 10 × NEBuffer 3.1. After mixing 8 times gently by MCA tips, the PCR plate was incubated at 37 °С for 2 h followed by 65 °С for 30 min. Then, 5 μL of 10 mg/mL RNase A was added, mixed gently 8 times and the plate was incubated at 37 °С for 30 min. Following RNase A treatment, 5 μL of 20 mg/mL proteinase K was added and the plate was incubated at 50 °С for 30 min. The reaction mixture was then transferred into a 96 deep-well plate and 0.5-fold volume of magnetic beads (Mag-Bind^®^ RxnPure Plus, M1386-01, Omega BIO-TEK) was added. After being absorbed on magnetic stand for 10 min, the supernatant was removed and the beads were washed using 70% ethanol (v/v) two times (200 μL of each time). The DNA was eluted from beads with 20 μL of 10 mM Tris-HCl pH 8.0 and incubated at 50 °C until fully dissolved. The detailed procedures were presented as Step 3 in Supplementary Fig. 3.

### BGC plasmid construction and preparation

Linear assembly of DNA receivers and digested gDNA was performed in a 25 μL of reaction in a 96-well PCR plate containing 10 μL of digested gDNA (3-4 μg), 5 μL of DNA receiver #1 (∼15 ng of amplified from pBE44, or ∼45 ng of amplified pBE48), 5 μL of DNA receiver #2 (∼35 ng of amplified from pBE45) and 5 μL of T4 DNA polymerase master mix (0.75 U of T4 DNA polymerase, 2.5 μL of NEBuffer 2.1, 2.25 μL of H_2_O). After gently mixing 8 times, the mixture was incubated at 25 °C for 1 h, 75 °C for 20 min, and 50 °C for 30 min followed by 10 °C hold. Then, 5 μL of master mix containing 1 μL of 1 mM NAD^+^, 0.4 μL of 10 mM dNTPs, 1 μL (3 U) of T4 DNA polymerase, 1 μL of *E. coli* DNA ligase, and 1.6 μL of H_2_O were added and after gently mixing 8 times, the mixture was incubated at 37 °C for 1 h, 75 °C for 20 min, and stored at 10 °C until transformation. 20 μL of mixtures was transformed into 70 μL of electrocompetent *E. coli* NEB10β cells containing helper plasmid pBE14 (for Cre recombinase expression) prepared as described elsewhere^29^. Upon electroporation, all cells were plated on 8-well plates (Nunc™ Rectangular Dishes, 267062, Thermo Fisher Scientific) containing 5 mL/well of LB agar medium supplemented with apramycin (50 μg/mL) and X-GAL (40 μg/mL) and IPTG (0.1 mM) for blue/white screening, and incubated at 37 °C until colonies appeared. Then, the colonies were picked from each of the plates using Pickolo colonypicker (SciRobotics, Israel) and inoculated in a 96 deep-well plate with 1.5 mL of LB medium supplemented with 50 μg/mL apramycin in each well. After overnight culturing at 37 °C and 900 rpm in Cytomat_2C2 incubator, the cultures were spun down and the supernatants were removed by MCA tips. Then high throughput plasmid extraction on FluentControl 1080 using the PureLink™ Pro Quick96 Plasmid Purification Kit was performed following manufacturer’s protocol. All purified plasmids were digested with appropriate restriction enzymes and checked via automated parallel capillary gel electrophoresis using Fragment Analyzer (Supplementary Fig. 4). The detailed procedures were described in steps 4-5 in Supplementary Fig. 3.

### Heterologous expression in a *Streptomyces* host

The high-throughput di-parental (*E. coli* and *Streptomyces*) conjugation workflow was implemented on the iBioFAB (Supplementary Fig. 5). *E. coli* WM6026 chemical competent cells prepared following the protocols described elsewhere^29^ were aliquoted into a 96-well PCR plate with 50 μL/well, mixed with ∼400 ng of BGC plasmids and incubated at 0 °C for 20 min. The 96-well plate was then transferred to preheated 42 °C RIC20 Remote Controlled Chilling/Heating Dry Bath (Torrey Pines Scientific) for 2 min and then transferred to precooled 0 °C adaptor for another 3 min. Subsequently, 80 μL of LB medium was added into each well and incubated at 37 °C for 45 min. Cell cultures were plated onto 8-well plates containing 5 mL/well LB agar medium supplemented with apramycin (50 μg/mL) and 2,6-diaminopimelic acid (45 μg/mL), and incubated at 37 °C for overnight. The colonies were picked with Pickolo colonypicker and inoculated into 96 deep-well plates containing 800 μL of LB medium supplemented with apramycin (50 μg/mL) and 2,6-diaminopimelic acid (45 μg/mL), and incubated at 37 °C overnight. In the following day, the seed cultures (1%, v/v) were inoculated into 24 deep-well plates containing 4 mL/well of LB medium supplemented with apramycin (50 μg/mL) and 2,6-diaminopimelic acid (45 μg/mL) and incubated at 37 °C until OD_600_ reached up to 1.0. Cell cultures were harvested through centrifugation at 3,500 rpm for 5 min and the supernatants were discarded. 1 mL of LB medium was added to each well and the cell pellets were resuspended. After centrifugation at 3,500 rpm for 5 min, the supernatants were discarded. After repeating wash step three times, the cell pellets were ready for conjugation usage.

*S. lividans* TK24 heterologous host strains were recovered on MS (20 g/L of D-mannitol, 20 g/L of soybean powder, 20 g/L of agar) solid plate. After culturing for 3∼4 d, the spores were collected using sterile cotton swabs. The spores were then washed using 2 × YT broth (16 g/L of Bacto Tryptone, 10 g/L of Bacto yeast extract, and 5 g/L of NaCl, pH 7.0) for two times. After centrifugation and discarding the supernatants, the spores were resuspended into appropriate 2 × YT broth (200 μL of each transformation) and heated shock at 50 °C for 10 min. Then, the spores were aliquoted into the 24 deep-well plates containing the *E. coli* WM6026 cell pellets prepared above. After mixing using MCA tips for 10 times, the mixture was plated on 8-well plates containing 5 mL/well MS agar medium supplemented with 10 mM MgCl_2_ solution and incubated at 30 °C for 16∼20 h. In the next day, 300 μL/well of antibiotics solution containing 100 μL of 50 mg/mL apramycin, 10 μL of 25 mg/mL nalidixic acid (N8878-25G, Sigma-Aldrich) and 190 μL of sterile H_2_O overlaid on the conjugation plates and incubated at 30 °C until the exconjugants appeared (5∼6 d). The exconjugants were picked with Pickolo colonypicker and inoculated into 96 deep-well plates containing 1.5 mL of TSB medium (Tryptic soy broth, BD 211825, Thermo Fisher Scientific) supplemented with apramycin (50 μg/mL) and nalidixic acid (25 μg/mL), and incubated at 30 °C, 900 rpm until the strains grew well.

### Fermentation, isolation, and detection of natural products

*S. lividans* TK24 strains containing corresponding BGC plasmids were inoculated into 24 deep-well plates containing 5 mL/well MYM medium (10 g/L of malt extract, 4 g/L of yeast extract, 4 g/L D-maltose) (two replicate wells of each strain) and incubated at 30 °C for 6 d. Fermented cultures in 24 deep-well plates were centrifuged at 3,900 rpm for 5 min to separate supernatants and strains. The supernatants were extracted two times, respectively, by ethyl acetate (5 mL/well) in ultrasonic for 20 min of each time. After centrifugation at 3,900 rpm for 5 min, the organic phase was combined and dried by air vacuum in a fume hood to get crude extracts. The strains were soaked in 300 μL of methanol and extracted by ultrasonic for 1 h. The detailed procedures were described in **Supplementary Fig. 5**.

Samples were analyzed using Q Exactive Orbitrap (Thermo Fisher Scientific, USA) equipped with a Vanquish HPLC system (Thermo Fisher Scientific, USA) loaded a Thermo hypersil GOLD aQ column (3 μm, 2.1 × 150 mm; PN 25303-152130) using the following buffers: buffer A (100% water, 0.1 % formic acid) and buffer B (100% acetonitrile, 0.1 % formic acid). A linear gradient from 5% to 95% buffer B over 18 min followed by an isocratic elution with 95% buffer B held 5 min was then applied at a flow rate of 0.2 mL/min. MS spectra were acquired on positive/negative ionization modes using Q Exactive Orbitrap with the scan range *m/z* 200-3000, resolution 70,000 FWHM, AGC target 3 × 10^6^ ions capacity, Maximum IT 100 ms. In MS/MS (MS2) mode, the parameters were set as: resolution 17, 500 FWHM; AGC target 1 × 10^5^ ions capacity; Maximum IT 50 ms.

### Isolation and structural characterization of natural products

The *Streptomyces* strains producing detectable natural products were selected for scale-up fermentation in 5-10 L MYM medium. After culturing around 6 d, the cultures were extracted 3 times with ethyl acetate. The combined extracts were dried and evaporated to dryness. The crude extracts were generally separated and purified by reverse-phase semipreparative HPLC. Semipreparative HPLC was carried out on an Agilent 1260 Infinity series instrument and the following program was run: 5% B to 50% B (linear gradient, 0–15 min), 50% B to 100% B (15–20 min), 100% B (isocratic elution, 20–24 min), 100% B to 5% B (24–25 min), 5% B (25– 30 min); the solvent system comprises solvent A (water supplemented with 0.1% trifluoroacetic acid) and solvent B (acetonitrile supplemented with 0.1% trifluoroacetic acid); flow rate at 1 mL/min. NMR spectroscopic data (^1^H, ^13^C, HSQC, COSY, and HMBC) were collected by Agilent 600 MHz (14.1 Tesla) and analyzed by MestReNova version 11.0.3. HR-ESI-MS data were analyzed by MassLynx software. Structures were drawn using ChemBioDraw Ultra 14.0.

### Antimicrobial assays

The antimicrobial activities of characterized compounds were measured against 10 types of gram-positive bacteria using 2-fold broth microdilution method^4^. These indicator strains included *Bacillus halodurans* C-125, *Bacillus anthracis* str. Sterne, *Bacillus cereus* TZ417, *Bacillus subtilis* ATCC6633, *Enterococcus faecium* U503, *Lactococcus lactis* CNRZ 481, *Micrococcus leteus* ATCC 4698, *Staphylococcus aureus* USA300, *Staphylococcus epidermidis* 15 × 154, *Streptococcus mutans* ATCC 25175. *B. halodurans* C-125, *B. cereus* TZ417, *B. subtilis* ATCC6633, *L. lactis* CNRZ 481 and *M. leteus* ATCC 4698 were grown in 5 mL of TSB at 30 °C. The other five strains were grown in Brain Heart Infusion (BHI, 53286-500G, Sigma-Aldrich) at 37 °C. Overnight cultures were diluted into fresh sterilized medium to achieve an optical absorbance of 0.004∼0.006 at 600 nm and distributed into 96-well microtiter plates, which were supplemented with compounds ranging from 100 to 0.39 μg/mL. The Minimum Inhibitory Concentrations (MIC) value was the concentration that resulted in no visible growth after 16∼18 h cultivation. All the experiments were tested in triplicates.

### Cytotoxicity assays

The cytotoxicity of compounds was tested on 4 types of human cancer cell lines, HCT-116 (human colon carcinoma, ATCC CCL-247™), Hela (human cervical carcinoma, ATCC CCL-2™), HepG2 (human liver carcinoma, ATCC HB-8065™), A549 (human non-small cell lung cancer cells, ATCC CCL-185™), using the Cell Counting Kit-8 (CCK-8) (CK04-13, Dojindo, Japan) method described below. The exponentially growing cells were seeded into 96-well plates with a density of 2.0 × 10^4^ cells/well and incubated for 24 h at 37°C with 5% CO_2_. The tested compounds at indicated concentrations (3.125 μg/mL, 6.25 μg/mL, 12.5 μg/mL, 25 μg/mL, 50 μg/mL, 100 μg/mL, 200 μg/mL) were added to each well accordingly and then incubated for another 24 h. Later, the CCK-8 solution (10 μL) was added to each well and incubated for another 1∼1.5 h. The absorbance was measured at 450 nm on the SpectraMax MX5 (Molecular Devices, CA) Plate Reader. Wells without drug was used as blanks. All the experiments were performed in three independent biological replicates. The IC_50_ values were measured using GraphPad Prism 8 software.

## Data availability

Data supporting the findings of this study are available within the article and its Supplementary Information files.

## Acknowledgements

We thank Behnam Enghiad for assistance with the direct cloning of BGCs. This work was financially supported by the U.S. National Institutes of Health (AI144967 to H.Z.).

## Author contributions

Y.Y. and H.Z. conceived and designed the study. Y.Y., C.H. and H.Z. wrote the manuscript. Y.Y. and N.S. performed the automation workflow development experiments. Y.Y. and C.H. performed the products fermentation and detection experiments. C.H. performed structure elucidation of compounds. Y.Y. performed the antimicrobial activity assays. Y.Y. and G.X. performed the anti-tumor activity assays.

