## Supplementary Materials for "Automated, self-resistance gene-guided, and high-throughput genome mining of bioactive natural products from *Streptomyces*"

**Table of Contents:**

Supplementary Text P3-P4
Supplementary Figure 1 P5
Supplementary Figure 2 P6
Supplementary Figure 3 P7
Supplementary Figure 4 P8
Supplementary Figure 5 P9
Supplementary Figure 6 P10
Supplementary Figures 7-8: NMR data of compound **1**. P11
Supplementary Figure 9: HR-ESI-MS data of compound **2**. P12
Supplementary Figures 10-14: NMR data of compound **2**. P13-P15
Supplementary Figure 15: HR-ESI-MS data of compound **3**. P16
Supplementary Figure 16-20: NMR data of compound **3**. P17-P19
Supplementary Figure 21: HR-ESI-MS data of compound **4**. P20
Supplementary Figure 22-26: NMR data of compound **4**. P21-P23
Supplementary Figure 27: HR-ESI-MS data of compound **5**. P24
Supplementary Figure 28-32: NMR data of compound **5**. P25-P27
Supplementary Figure 33: HR-ESI-MS data of compound **6**. P28
Supplementary Figure 34-38: NMR data of compound **6**. P29-P31
Supplementary Figure 39: HR-ESI-MS data of compound **7**. P32
Supplementary Figure 40-44: NMR data of compound **7**. P33-P35
Supplementary Figure 45-46: NMR data of compound **8**. P36
Supplementary Figure 47-48: NMR data of compound **9**. P37
Supplementary Figure 49-50: NMR data of compound **10**. P38
Supplementary Figure 51: NMR data of compound **11**. P39
Supplementary Table 5 P40
Supplementary Table 6: NMR data of compound **1-2**. P41
Supplementary Table 7: NMR data of compound **3-4**. P42
Supplementary Table 8: NMR data of compound **5-6**. P43
Supplementary Table 9: NMR data of compound **7**. P44
**References** P45

**Supplementary Text**

**Structural characterization of natural products.** Compound **2** was isolated as a yellow crystal. The molecular formula of **2** was established as $C_{26}H_{18}O_{10}$ based on its negative HR-ESI-MS (*m*/*z* 489.0824 $[M-H]^-$, calcd 489.0827, **Supplementary Fig. 9**), suggesting 18 degrees of unsaturation. The $^1H$, $^{13}C$, and HSQC NMR spectroscopic data (**Supplementary Figs. 10–12**, **Supplementary Table 6**) of **2** revealed the presence of one methyl, two methylenes, five methines including four $sp^2$ hybrid olefinic methines and one oxygenated $sp^3$ hybrid methine, and eighteen quaternary carbons including fourteen olefinic ones and four carbonyl ones. Detailed analyses of one-dimensional (1D) and two-dimensional (2D) NMR of **2** clearly suggested the presence of a 1,6,8-trihydroxyanthraquinone moiety. In fact, they were highly similar to those of KS-619-1 (**1**)[1] except for an additional hydroxyl group attached to C-6 in **2**. This assignment was further supported by the COSY correlations of $H_2$-5/H-6 and HMBC correlations from H-6 to C-4a/C-6a/C-7/C-14a (**Supplementary Figs. 13-14**). Finally, the structure of **2** was determined and designated as grianquinone A (**2**).

Grianquinone B (**3**) has a molecular formula $C_{25}H_{18}O_8$, as determined by negative HR-ESI-MS (*m*/*z* 445.0913 $[M-H]^-$, calcd 445.0929, **Supplementary Fig. 15**). The NMR spectroscopic data (**Supplementary Figs. 16–20**, **Supplementary Table 7**) of **3** were strikingly similar to those of **2**. The only major difference of the NMR data between **3** and **2** was the presence of an olefinic proton ($\delta_H$ 6.73, H-2) in **3** in place of the carboxyl group ($\delta_C$ 175.7, C-18) in **2**. The HMBC correlations from H-2 to C-4/C-14b/C-15 and from $H_2$-15 to C-2/C-3/C-4 further confirmed this assignment (**Supplementary Fig. 20**). Therefore, the structure of **3** was identified as loss of $CO_2$ compared to **2** and was assigned as grianquinone B (**3**).

The molecular formula of compound **4** was obtained to be $C_{25}H_{18}O_8$ by negative HRESIMS (*m*/*z* 445.0908 $[M-H]^-$, calcd 445.0929, **Supplementary Fig. 21**), corresponding to 17 degrees of unsaturation. Careful analyses of 1D and 2D NMR (**Supplementary Figs. 22–26**, **Supplementary Table 7**) of **4** revealed that they were similar to the NMR signal patterns of the polyaromatic system of **3**, and allowed the construction of the same 1,6,8-trihydroxyanthraquinone moiety and the 3-hydroxybenzocyclohexanone moiety. The presence of 3-hydroxybenzocyclohexanone moiety was further supported by COSY correlation of H-5/H-6 ($\delta_H$ 7.72/8.56) and HMBC correlations from $H_2$-2 to C-1/C-3/C-4/C-14b, from $H_2$-4 to C-2/C-3/C-4a/C-5/C-14b, from 3-OH to C-2/C-3/C-4, from H-5 to C-4/C-6a/C-14b, and from H-6 to C-4a/C-7/C-14a (**Supplementary Figs. 25–26**). Accordingly, the structure of **4** was characterized as grianquinone C (**4**).

Compound **5** was obtained as a red needle crystal. Its molecular formula was established as $C_{26}H_{23}NO_9S$ by negative HR-ESI-MS (*m*/*z* 524.1016 $[M-H]^-$, calcd 524.1021, **Supplementary Fig. 27**). The 1D and 2D NMR spectroscopic data (**Supplementary Figs. 28–32**, **Supplementary Table 8**) of **5** were highly similar to those of γ-indomycinone containing an anthraquinone-γ-pyrone nucleus[2]. The difference was that the triplet methyl presented in γ-indomycinone was replaced by an *N*-acetylcysteine moiety. This assignment was confirmed by HMBC correlations from $H_2$-15 ($\delta_H$ 3.16/3.26) to C-1′ ($\delta_C$ 34.1), from $H_2$-1′ ($\delta_H$ 2.68/2.96) to C-15/C-2′/C-3′, from H-2′ ($\delta_H$ 4.30) to C-1′/C-3′, from $H_3$-5′ ($\delta_H$ 1.66) to C-4′, and from 2′-NH ($\delta_H$ 8.06) to C-2′/C-4′ (**Supplementary Fig. 32**). Finally, the structure of **5** was named as anthrapyrone A (**5**).

Compound **6** was assigned a molecular formula as $C_{22}H_{16}O_7S$ by negative HRESIMS (*m*/*z* 423.0530 $[M - H]^-$, calcd 423.0544, **Supplementary Fig. 33**), indicating 15 degrees of unsaturation. The comparison of $^1H$ and $^{13}C$ NMR spectroscopic data of **6** (**Supplementary Figs. 34–38**, **Supplementary Table 8**) and **5** suggested that an anthraquinone-γ-pyrone nucleus was also presented in **6**. In addition, the presence and attachment of a 3, 4-dihydroxytetrahydrothiophene moiety was supported by the COSY correlations of $H_2$-16/H-15/15-OH and the HMBC correlations from H-12 to C-14, from 14-OH to C-14/C-15/C-17, from 15-OH to C-14/C-16, and from $H_2$-17 to C-14/C-15/C-16 (**Supplementary Figs. 37-38**). Ultimately, the structure of **6** was designated as anthrapyrone B (**6**).

Anthrapyrone C (**7**) was assigned a molecular formula of $C_{21}H_{16}O_7$ based on HRESIMS (*m*/*z* 379.0812 $[M - H]^-$, calcd 379.0823, **Supplementary Fig. 39**). Likewise, the 1D and 2D NMR data (**Supplementary Figs. 40–44**, **Supplementary Table 9**) of **7** showed the presence of a chrysophanol moiety and a 2,3-dihydro-3-hydroxy-3-methyl-pyran-4-one moiety. The construction of the pyrone-like moiety was verified by HMBC correlations from H-12 ($\delta_H$ 5.50) to C-11/C-14, from $H_2$-15 ($\delta_H$ 4.34) to C-11/C-13/C-14/C-16, and from $H_3$-16 ($\delta_H$ 1.31) to C-13/C-14/C-15 (**Supplementary Fig. 44**). The HMBC correlation from H-12 to C-7 indicated the pyrone-like moiety attached at C-7 in **7**. Finally, the structure of **7** was determined as anthrapyrone C.

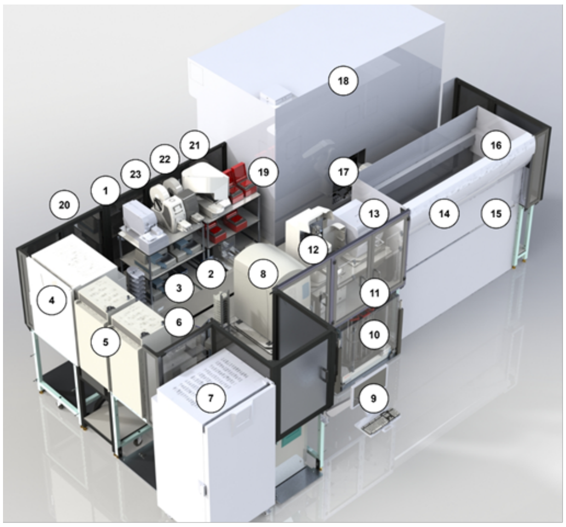


| No. | Equipment | Vendors/Models |
| --- | --- | --- |
| 1 | Xpeel_Peeler | Brooks |
| 2 | Centrifuge | Agilent |
| 3 | Multi-dispenser | Combi |
| 4 | Shakers and incubators | Cytomat_6K |
| 5 | Shakers and incubators | Cytomat_2C1, 2C2 |
| 6 | Robotic arm | Spinnaker_4K |
| 7 | Temperature-controlled incubators | Cytomat_C24 |
| 8 | Liquid handler | Labcyte Echo 550 |
| 9 | PC controls Echo and Fragment analyzer | HP |
| 10 | Plates/tips storage | Skyline_Hotel |
| 11 | Infinite_Plate reader | Tecan M1000 |
| 12 | Fragment analyzer | Agilent |
| 13 | Cytation5 cell imaging reader | Bio Tek |
| 14 | Liquid handler | Tecan FluentControl 1080 |
| 15 | Spiral plater and colony picker | SciRobotics |
| 16 | PC controls Momentum and all instruments | DELL |
| 17 | Central robotic arm | ThermoFisher F5 |
| 18 | Liquid handler | Tecan Freedom EVOware |
| 19 | Thermo cyclers | Biometra_Trobot |
| 20 | Plates/tips storage | Cytomat_Hotel |
| 21 | Barcode labeler | Agilent |
| 22 | ALPS_sealer | ThermoFisher |
| 23 | Automatic roll heat sealer | a4S 4titude |

**Supplementary Fig. 1.** Overview of the iBioFAB biofoundry. The system is integrated with different functional devices (# 1-23). This figure was adapted from ref. 3[3].

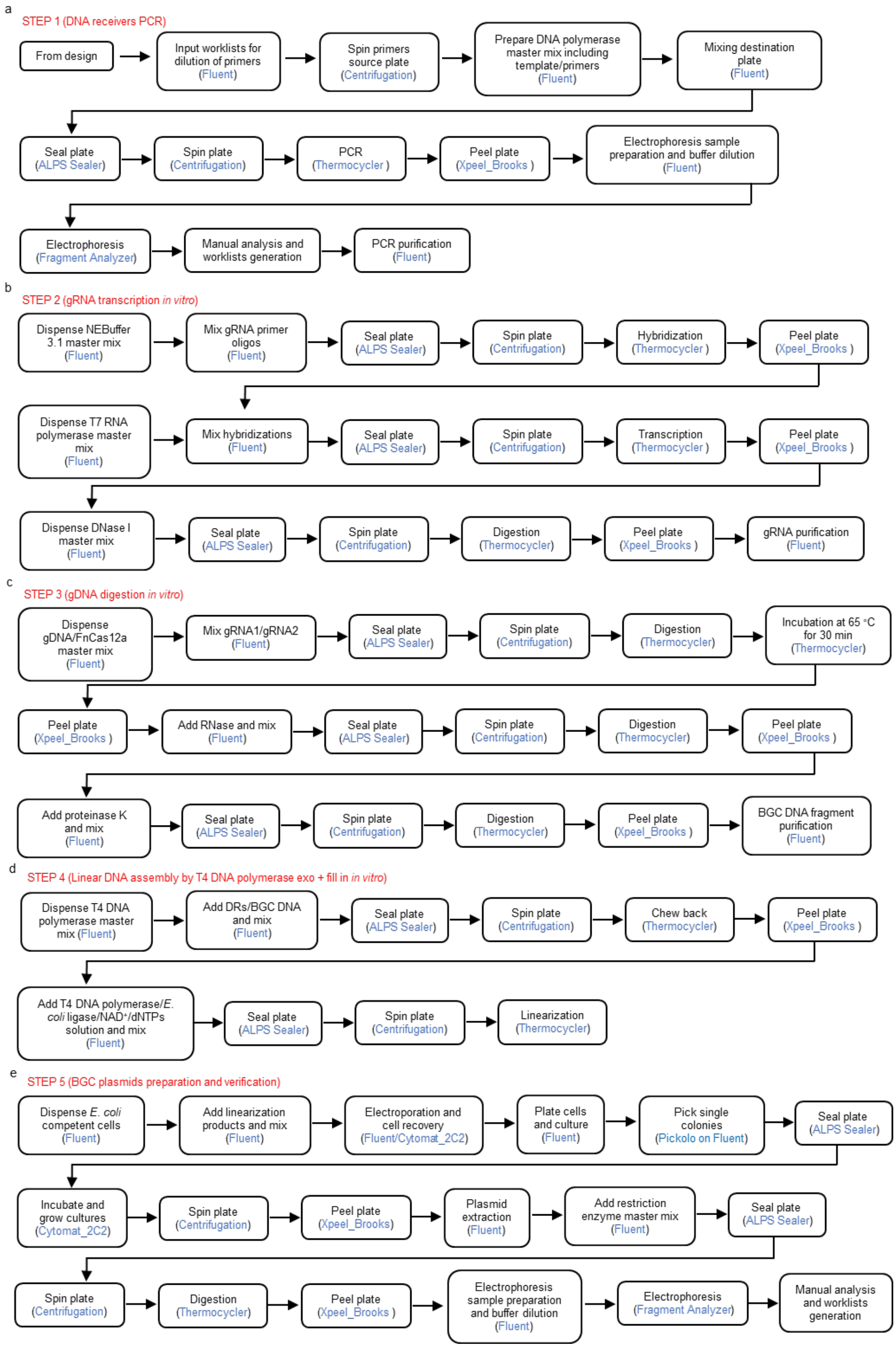


**Supplementary Fig. 2. Detailed workflow of the automated BGC direct cloning pipeline including specific inputs, outputs, and protocols being used**. **a**, DNA receivers (DRs) fragments preparation through PCR and purification. **b**, gRNA preparation through transcription *in vitro* and purification. **c**, gDNA digestion by FnCas12a enzyme *in vitro* and purification to generate target BGC DNA fragments. **d**, Linear assembly of DRs and BGC DNA fragments *in vitro* by T4 DNA polymerase exo + fill in. **e**, BGC plasmids construction by site-specific recombination mediated by Cre-*lox*P system, extraction and verification by restriction enzymes.

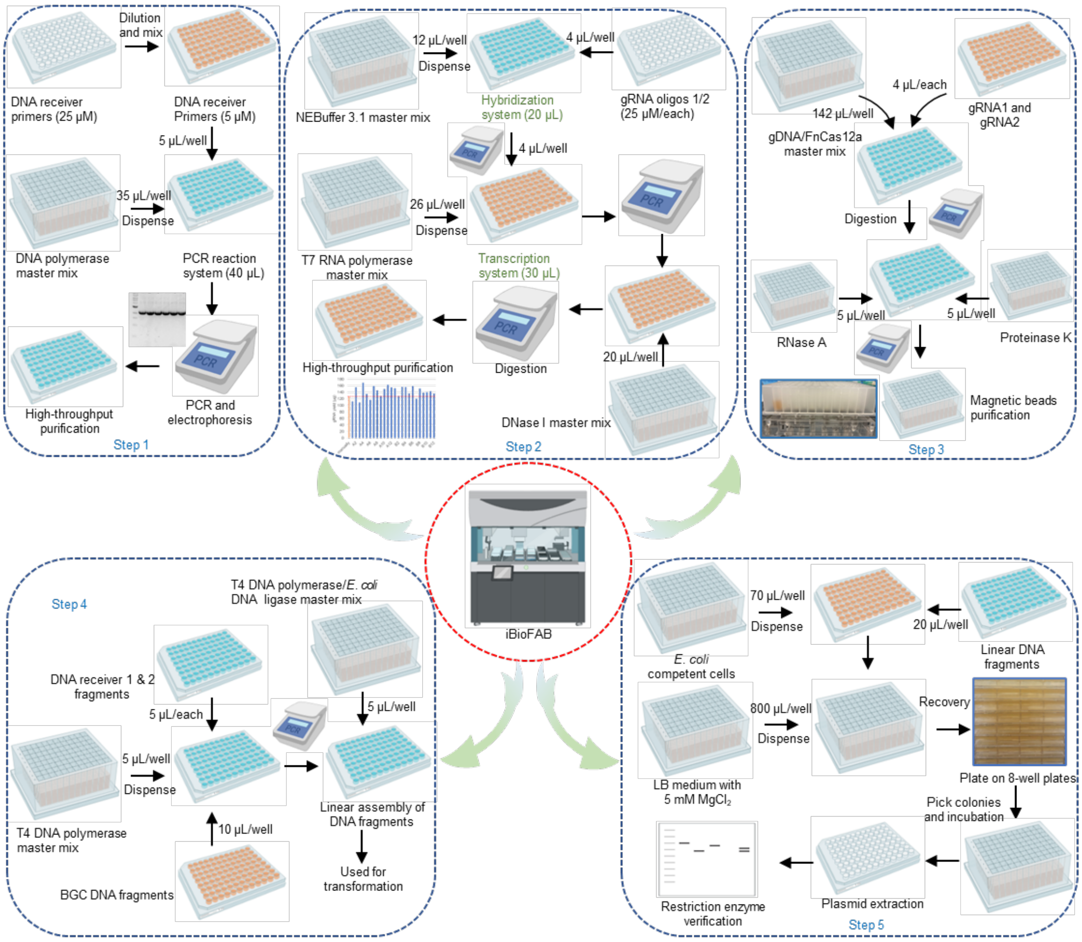


**Supplementary Fig. 3.** Schematic overview for the automated BGC direct cloning workflow implemented on iBioFAB biofoundry with the detailed procedures and equipment listed.

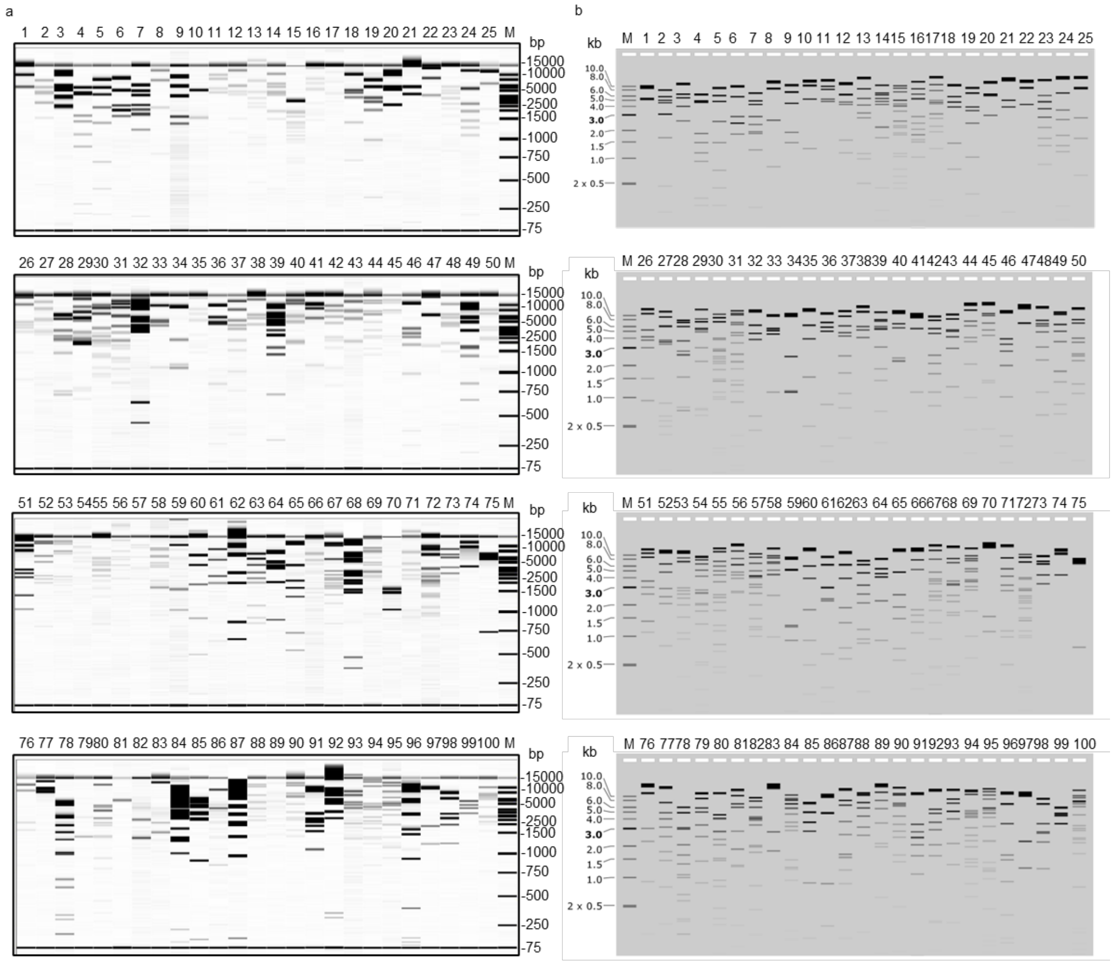


**Supplementary Fig. 4. Verification of automated BGC plasmid construction via corresponding restriction enzyme digestion**. **a**, Capillary gel electrophoresis analysis. The constructed 100 correct plasmids were checked by gel electrophoresis via Fragment Analyzer. Sample numbering and specific restriction enzymes being used for quality control corresponding to the 100 BGCs listed in **Supplementary Table 3**. 75-15,000 bp ladder is shown. **b**, Simulated gel electrophoresis result using SnapGene. 500-10,000 bp ladder is shown. M: marker.

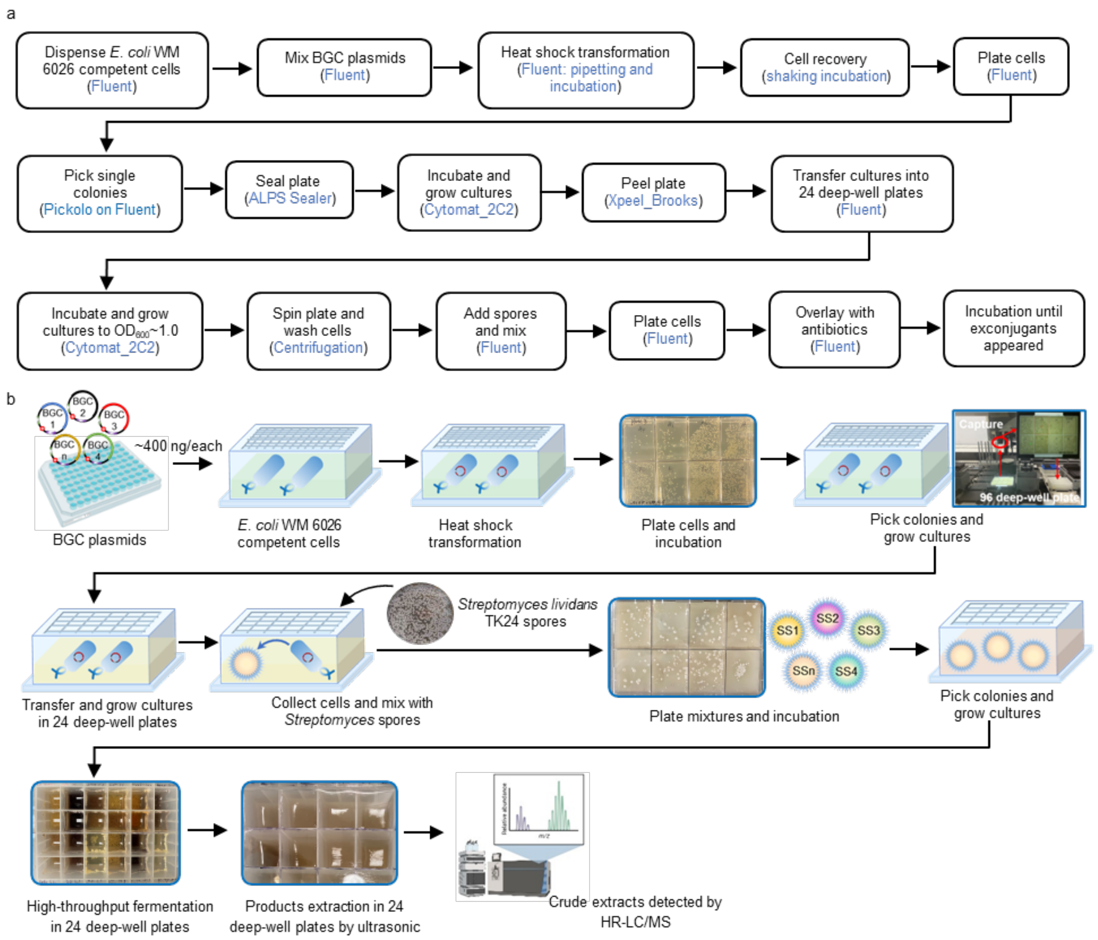


**Supplementary Fig. 5. High-throughput heterologous expression in *Streptomyces* strains (SSs) using iBioFAB biofoundry**. **a**, Detailed workflow of the automated heterologous expression pipeline of *Streptomyces* strains including specific inputs, outputs, and protocols being used. **b**, Schematic overview for the automated heterologous expression workflow of *Streptomyces* strains using iBioFAB biofoundry with detailed procedures and equipment listed.

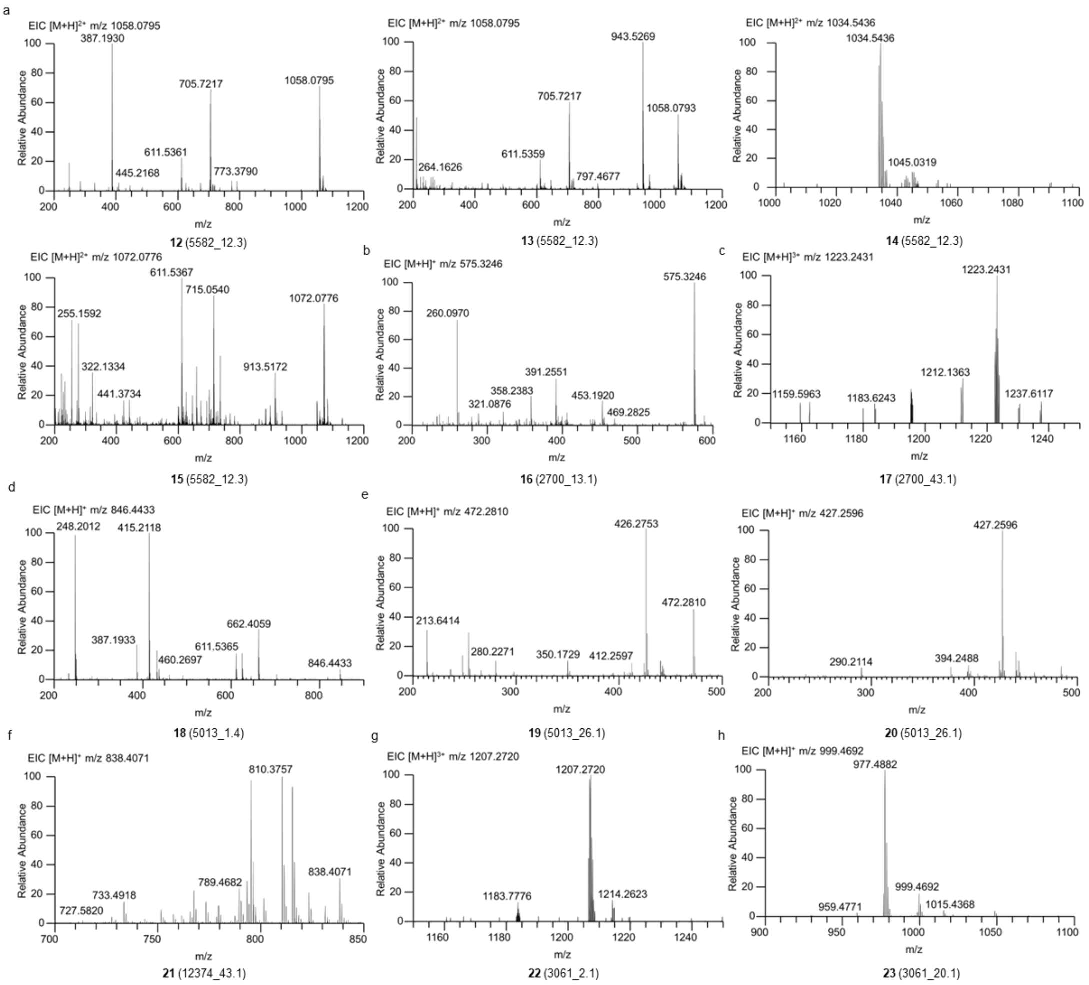


**Supplementary Fig. 6. HR-LC/MS determination of natural products (12-23)**. **a**, The MS spectrum of compounds **12-15** derived from BGC_5582_12.3. **b**, The MS spectrum of compound **16** derived from BGC_2700_13.1. **c**, The MS spectrum of compound **17** derived from BGC_2700_43.1. **d**, The MS spectrum of compound **18** derived from BGC_5013_1.4. **e**, The MS spectrum of compounds **19-20** derived from BGC_5013_26.1. **f**, The MS spectrum of compound **21** derived from BGC_12374_43.1. **g**, The MS spectrum of compound **22** derived from BGC_3061_2.1. **h**, The MS spectrum of compound **23** derived from BGC_3061_20.1.

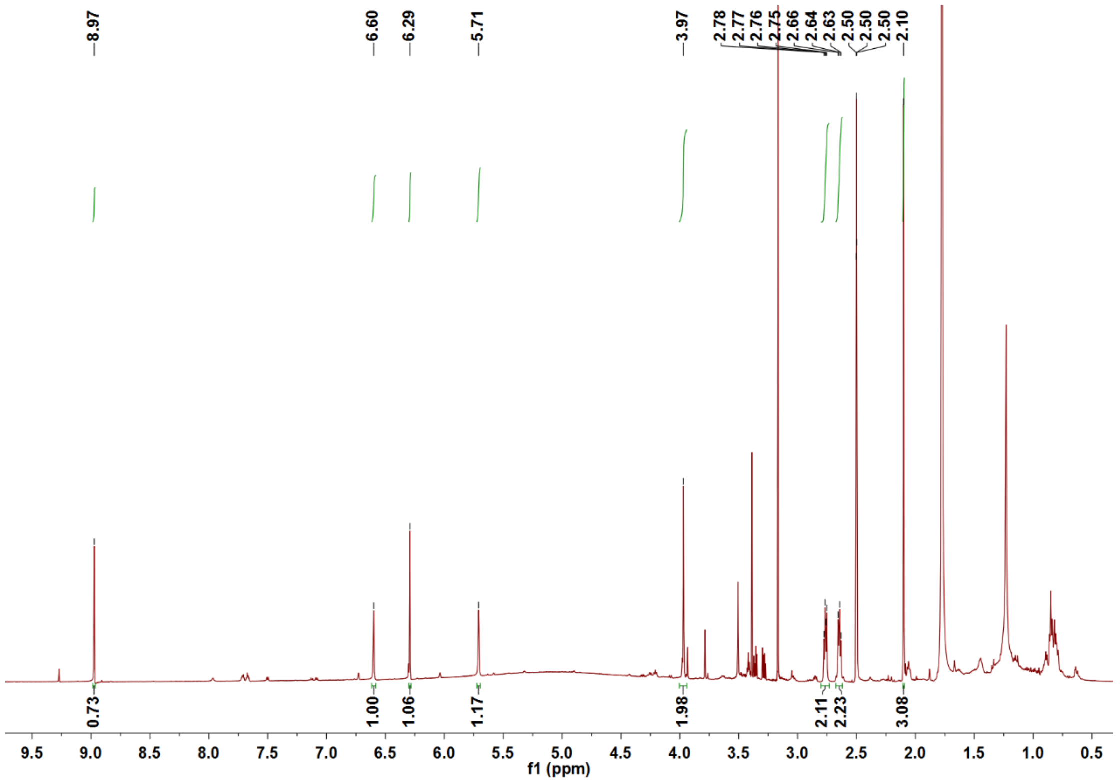


**Supplementary Fig. 7.** The $^{1}H$ NMR spectrum of KS-619-1 (**1**) in DMSO-$d_6$.

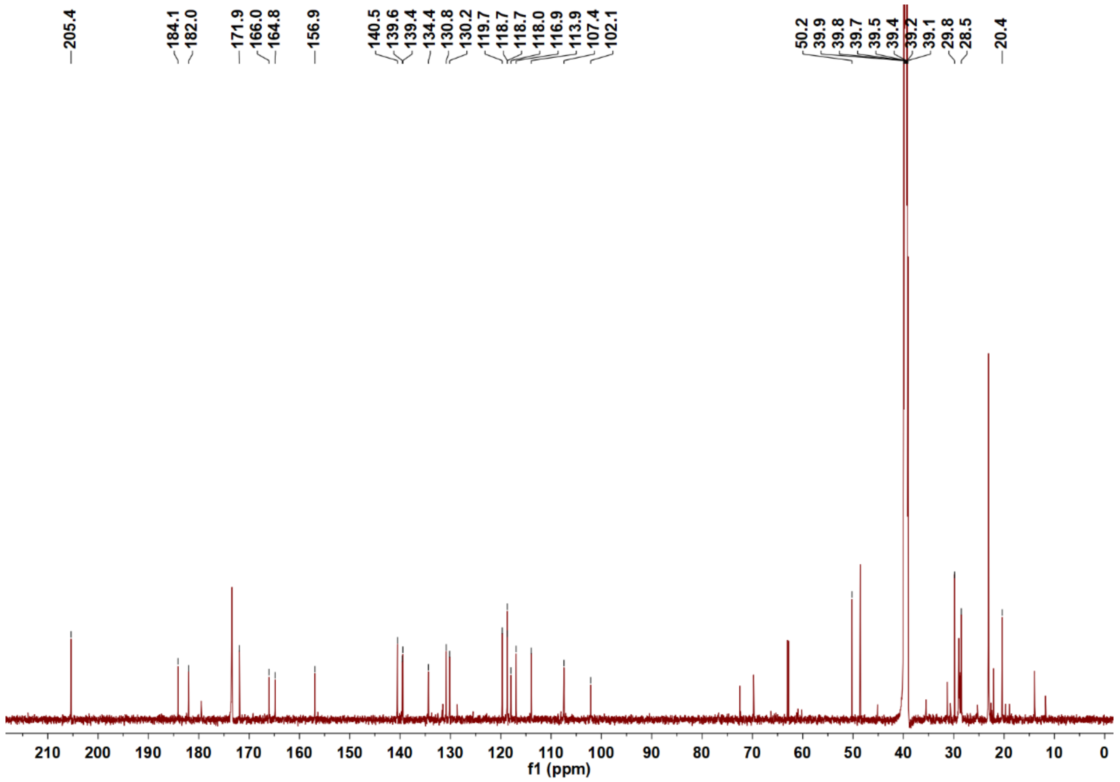


**Supplementary Fig. 8.** The $^{13}C$ NMR spectrum of KS-619-1 (**1**) in DMSO-$d_6$.

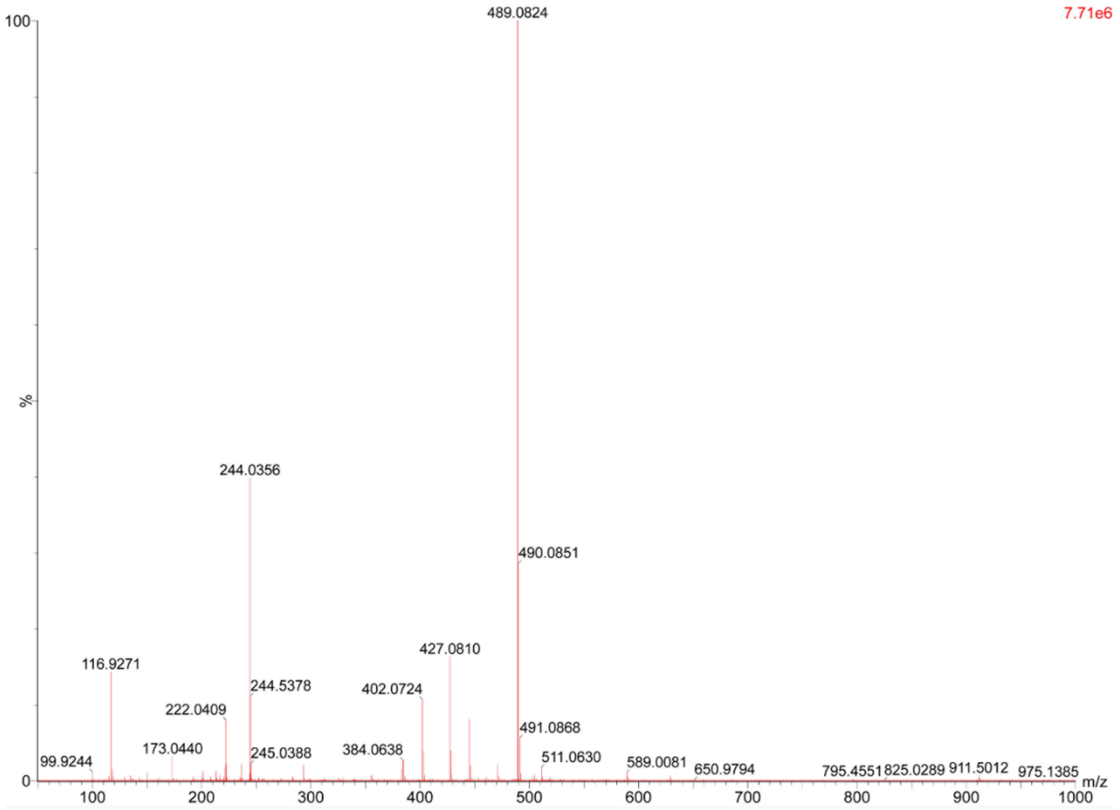


**Supplementary Fig. 9.** HR-ESI-MS spectrum of grianquinone A (**2**).

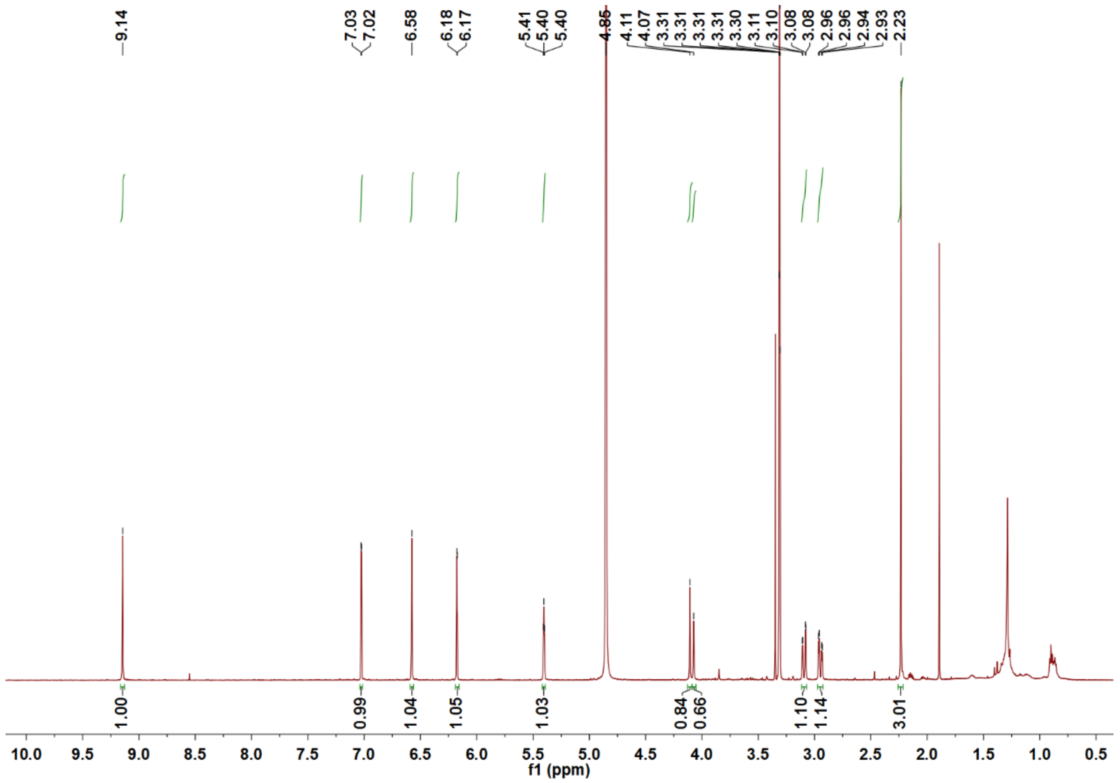


**Supplementary Fig. 10.** The [1]H NMR spectrum of **2** in $CD_3OD$.

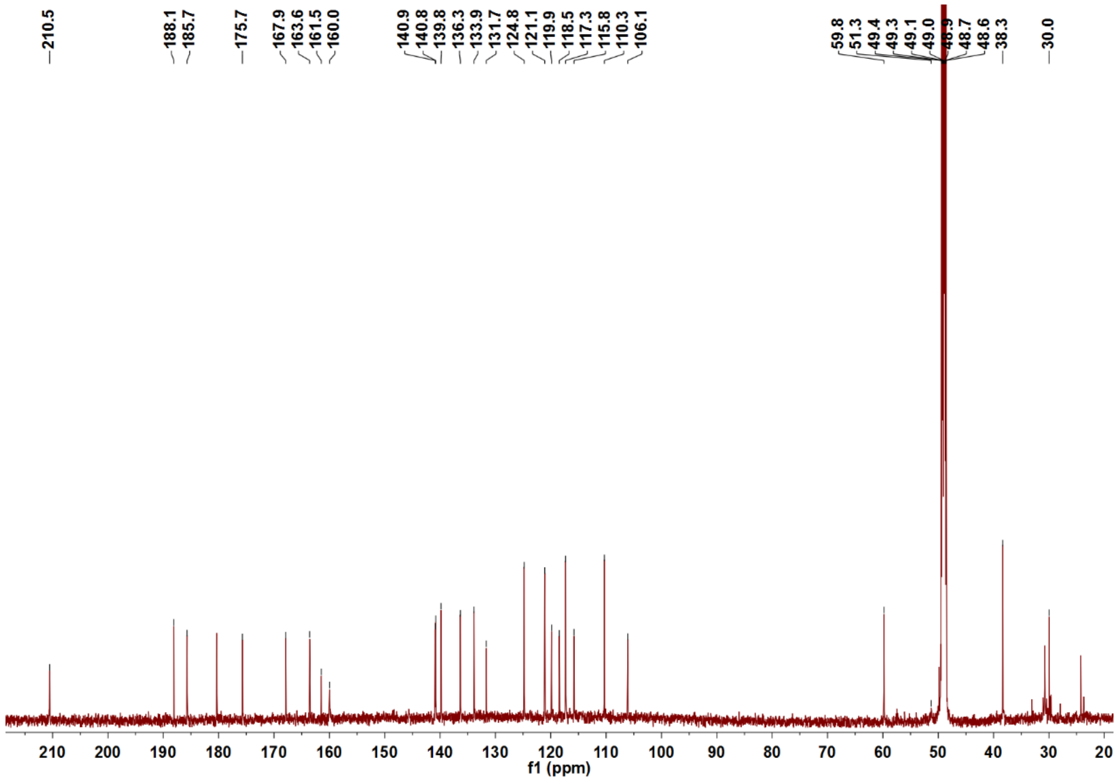


**Supplementary Fig. 11.** The [13]C NMR spectrum of **2** in $CD_3OD$.

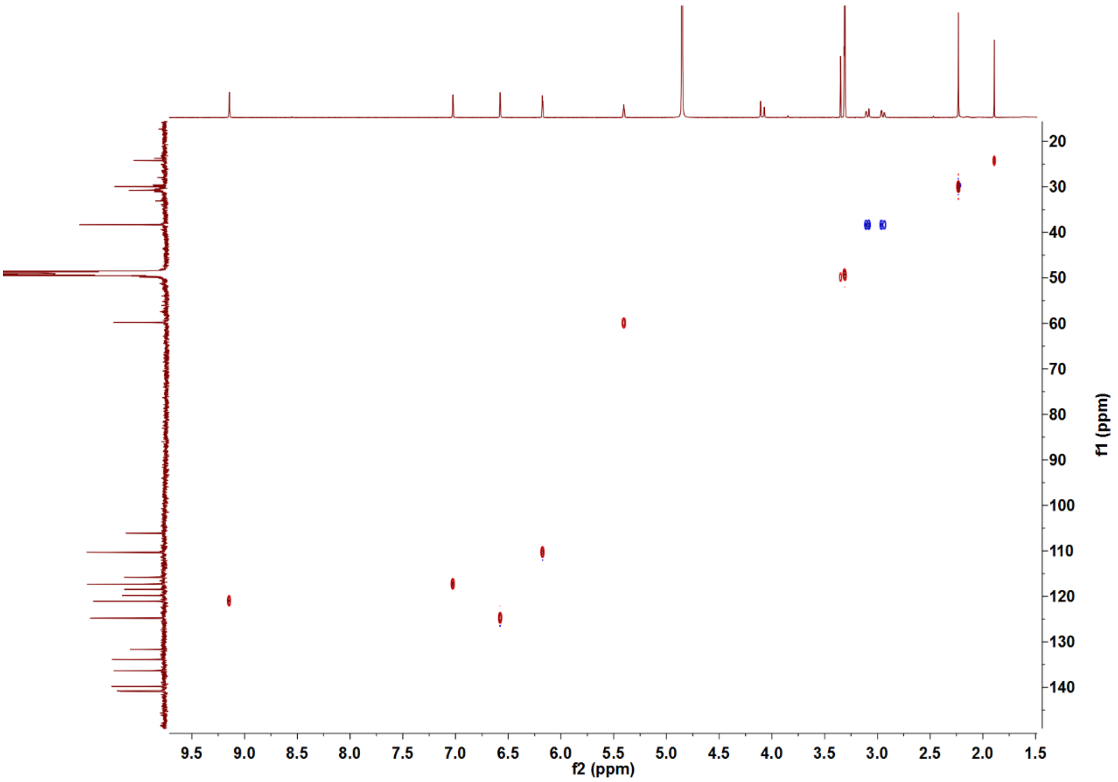


**Supplementary Fig. 12.** The HSQC spectrum of **2** in $CD_3OD$.

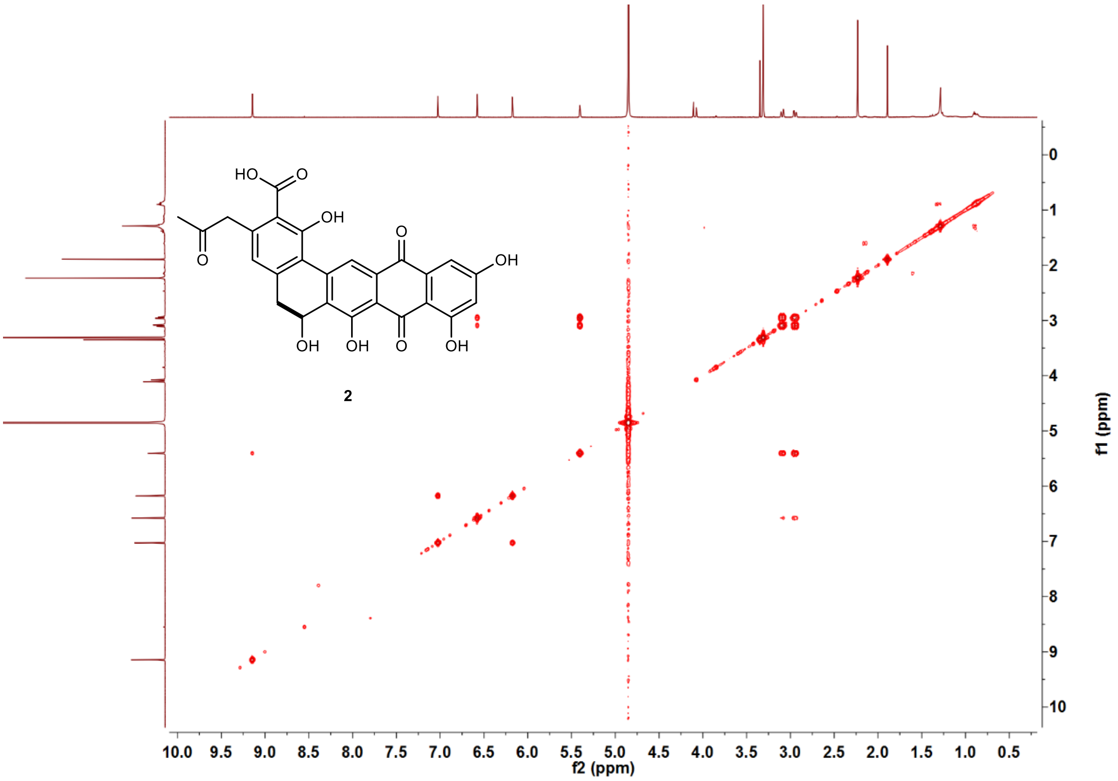


**Supplementary Fig. 13.** The COSY spectrum of **2** in $CD_3OD$.

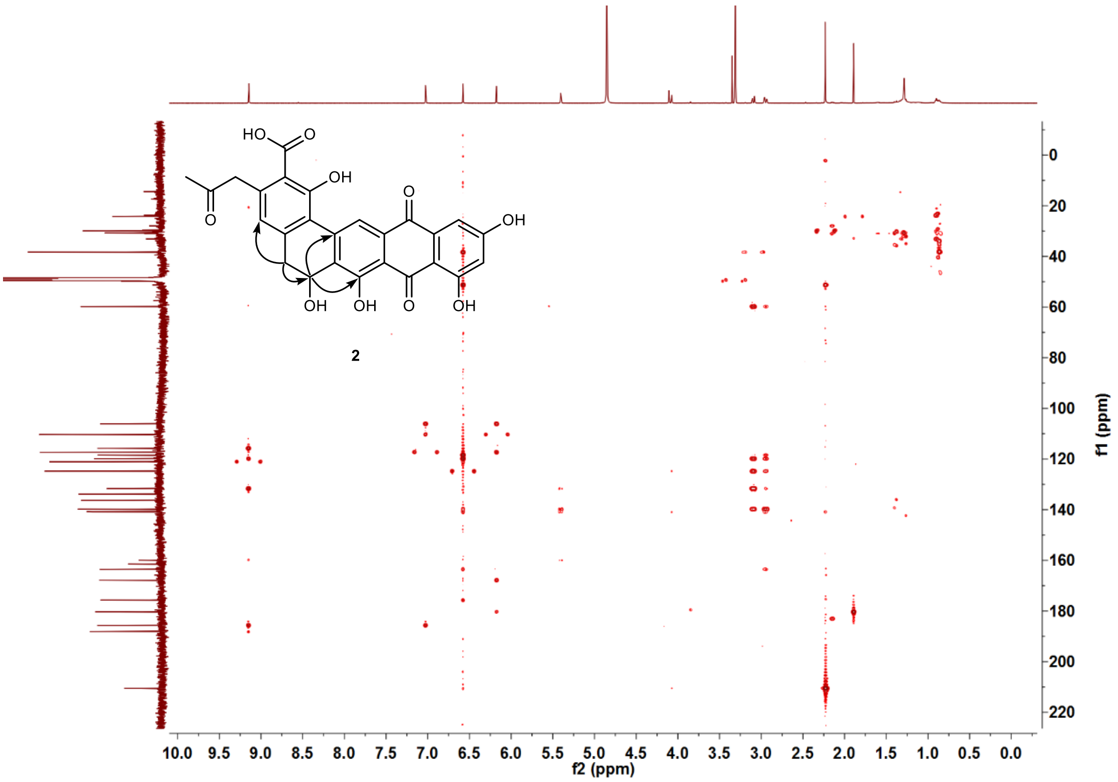


**Supplementary Fig. 14.** The HMBC spectrum of **2** in $CD_3OD$.

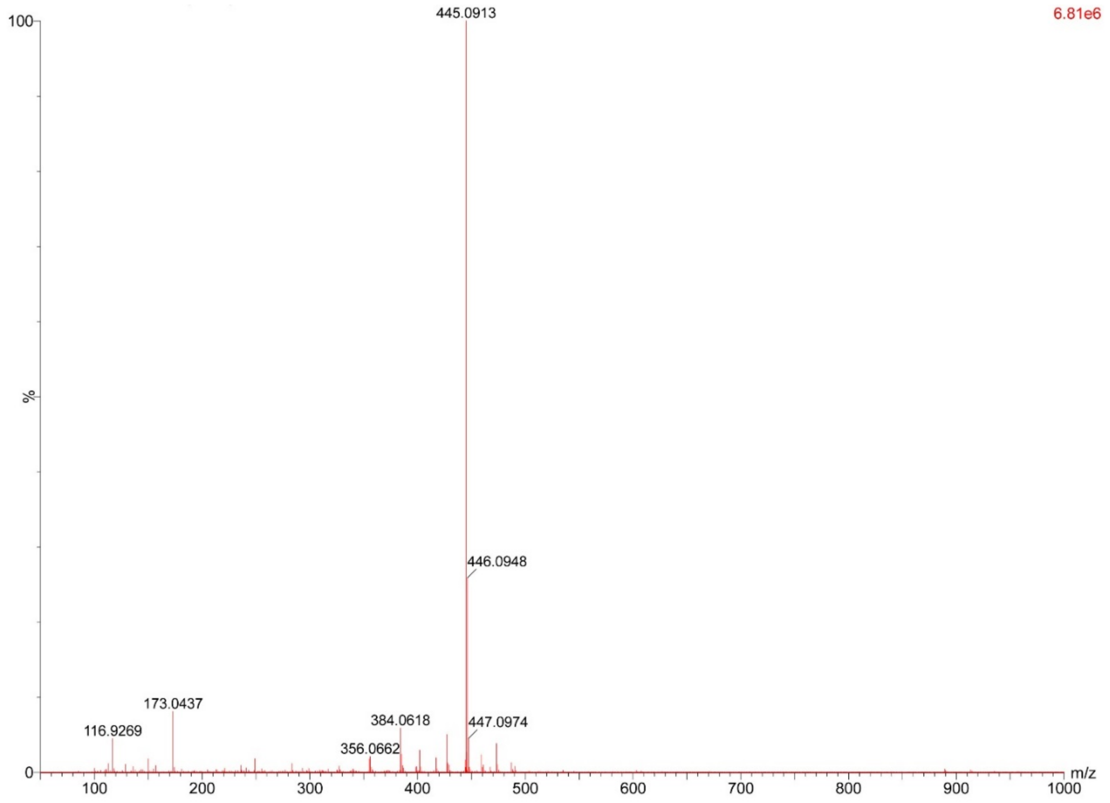


**Supplementary Fig. 15.** HR-ESI-MS spectrum of grianquinone B (**3**).

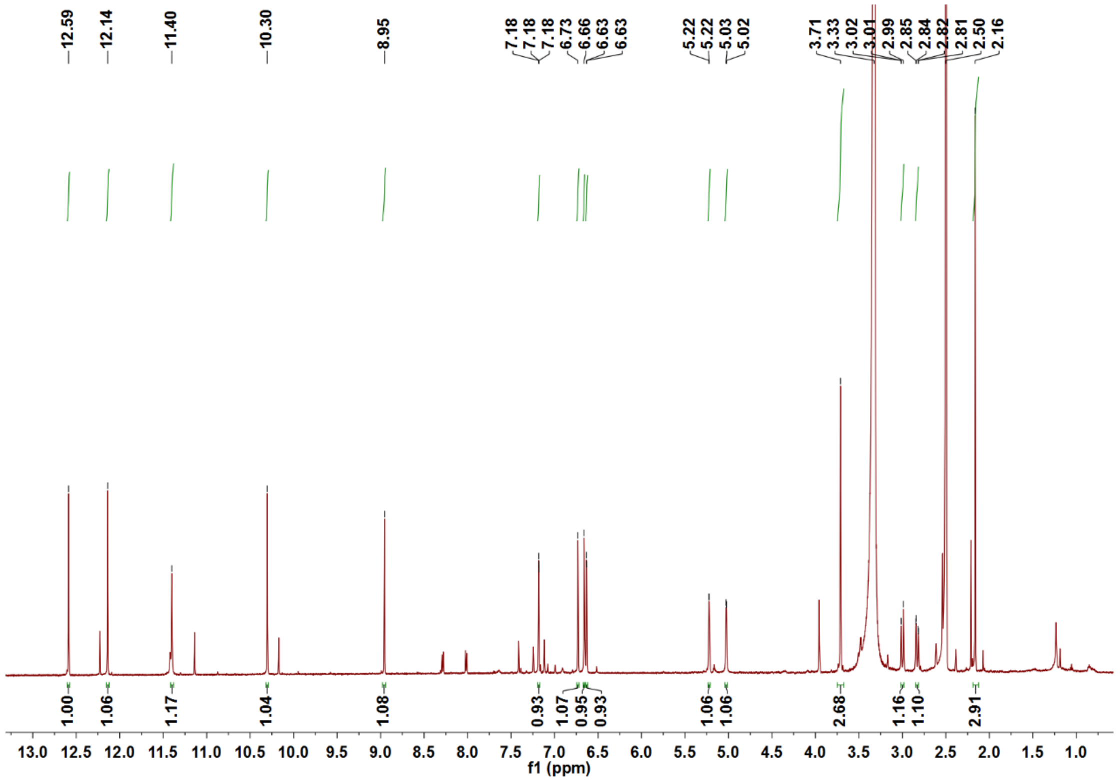

**Supplementary Fig. 16.** The $^{1}H$ NMR spectrum of **3** in DMSO-$d_6$.

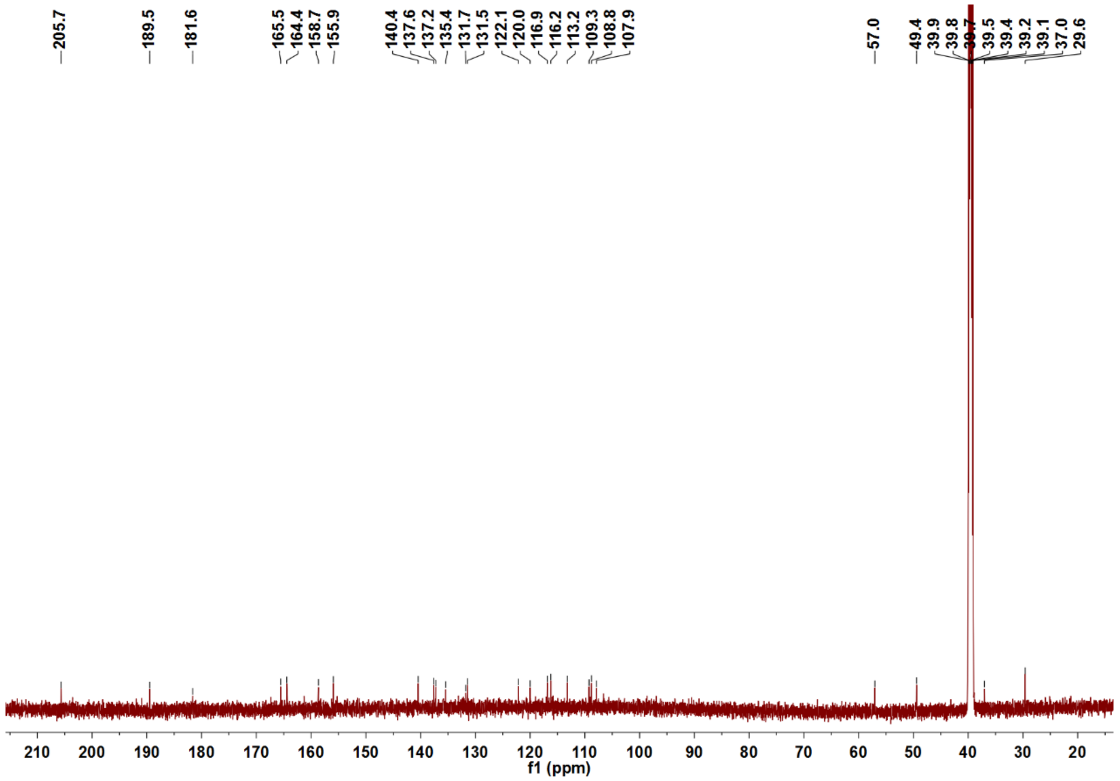

**Supplementary Fig. 17.** The $^{13}C$ NMR spectrum of **3** in DMSO-$d_6$.

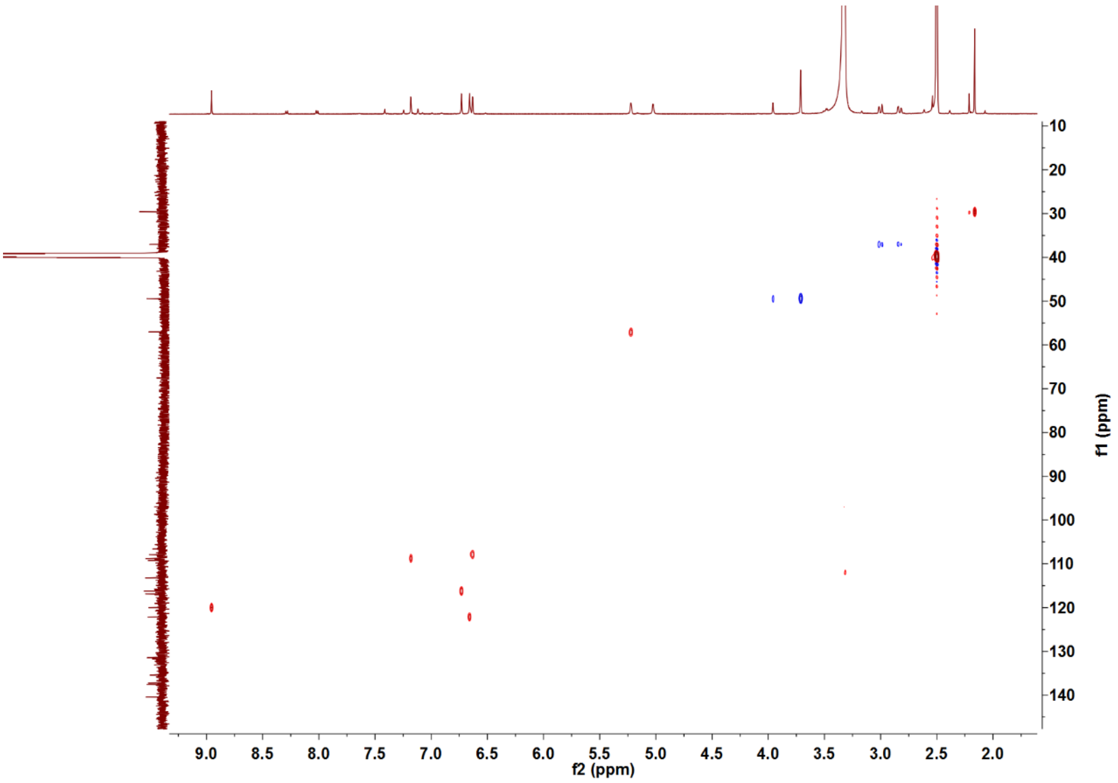


**Supplementary Fig. 18.** The HSQC spectrum of **3** in DMSO-$d_6$.

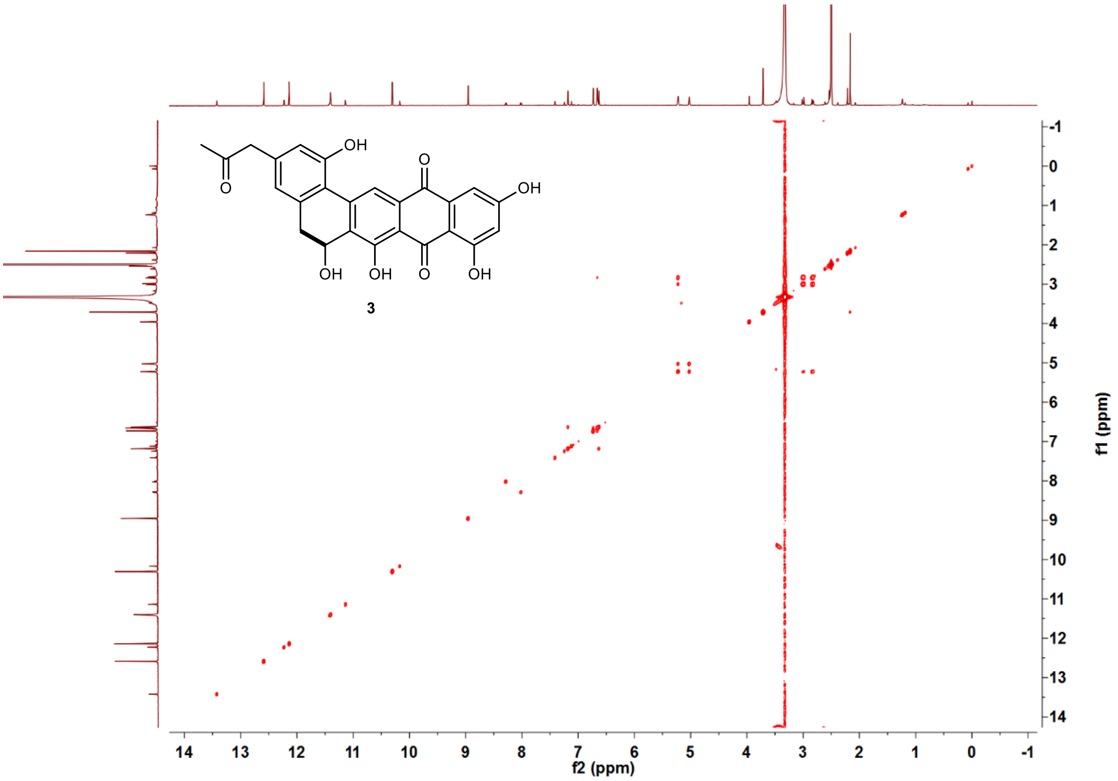


**Supplementary Fig. 19.** The COSY spectrum of **3** in DMSO-$d_6$.

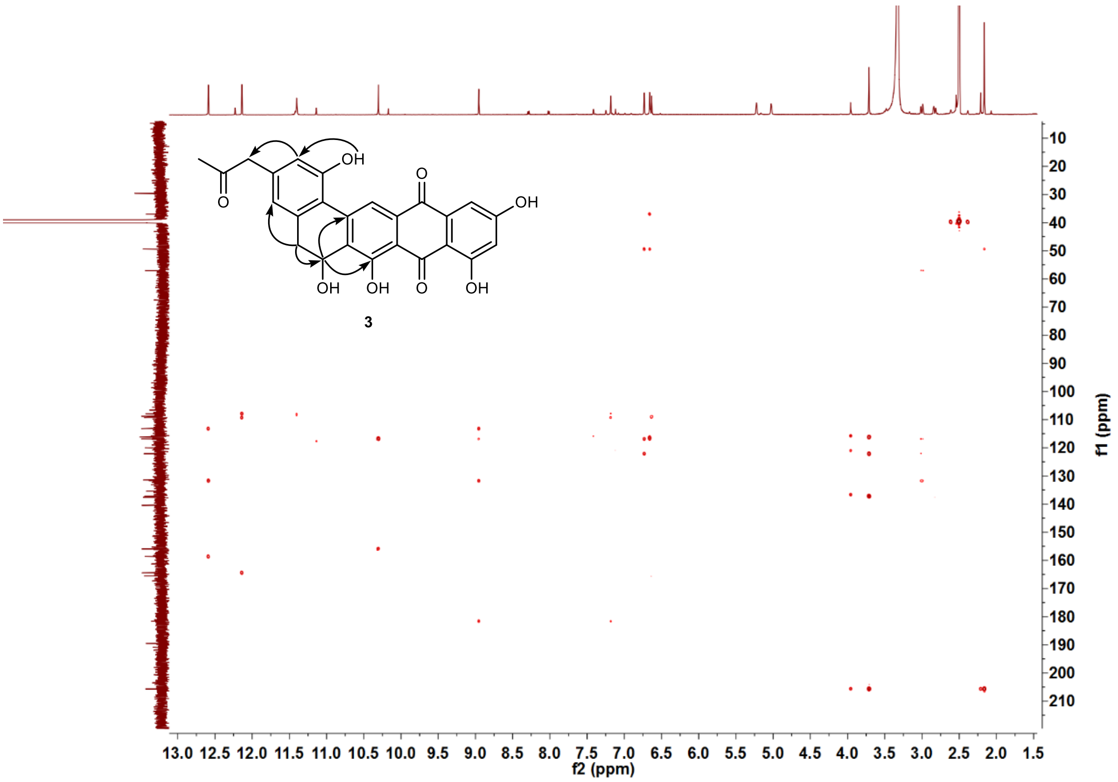


**Supplementary Fig. 20.** The HMBC spectrum of **3** in DMSO-$d_6$.

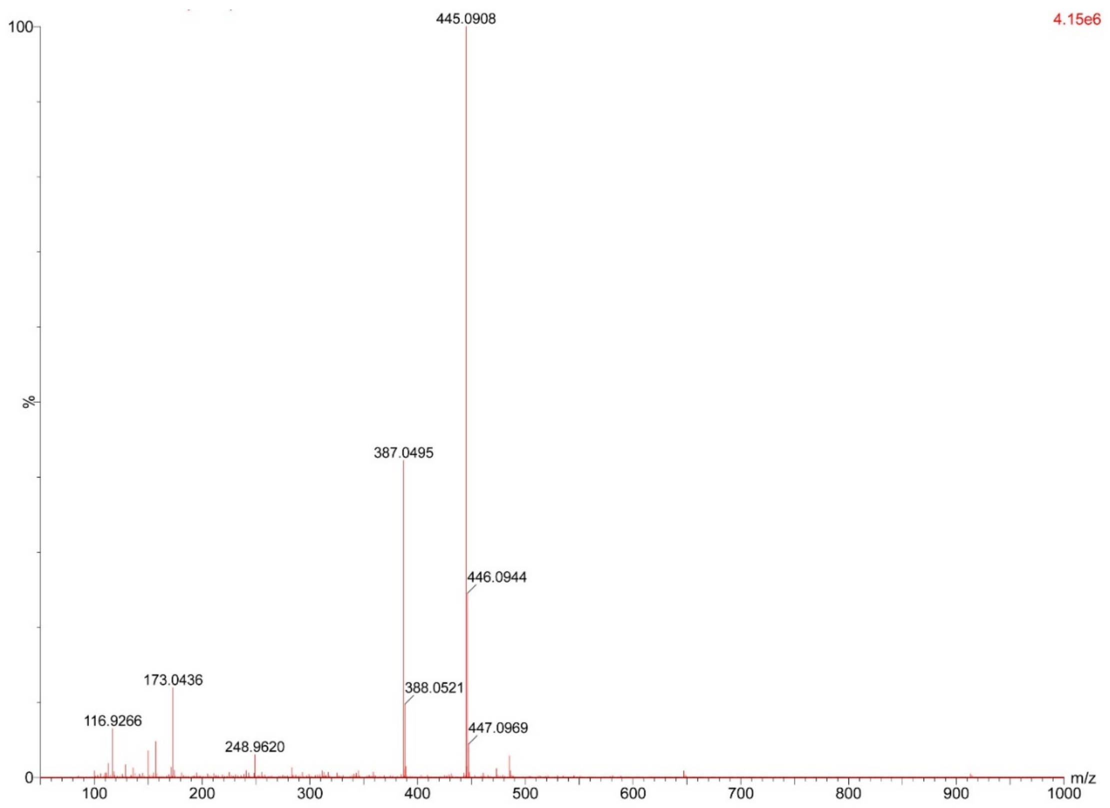


**Supplementary Fig. 21.** HR-ESI-MS spectrum of grianquinone C (**4**).

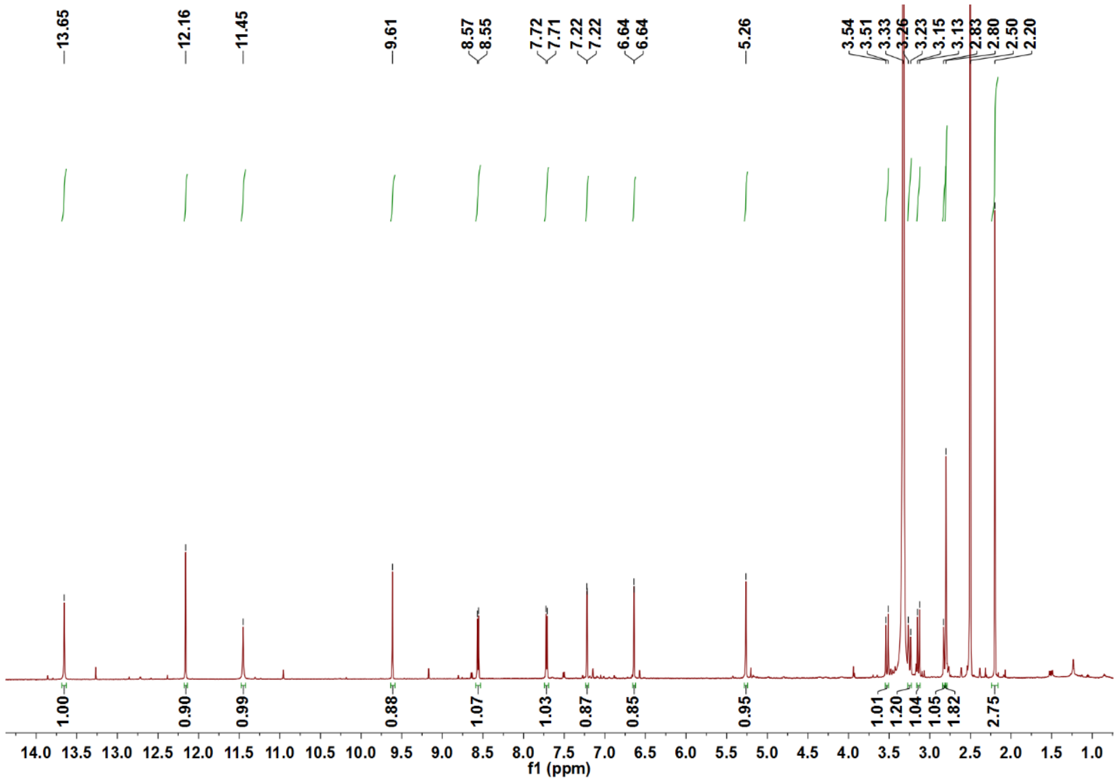


**Supplementary Fig. 22.** $^{1}$H NMR spectrum of **4** in DMSO-$d_6$.

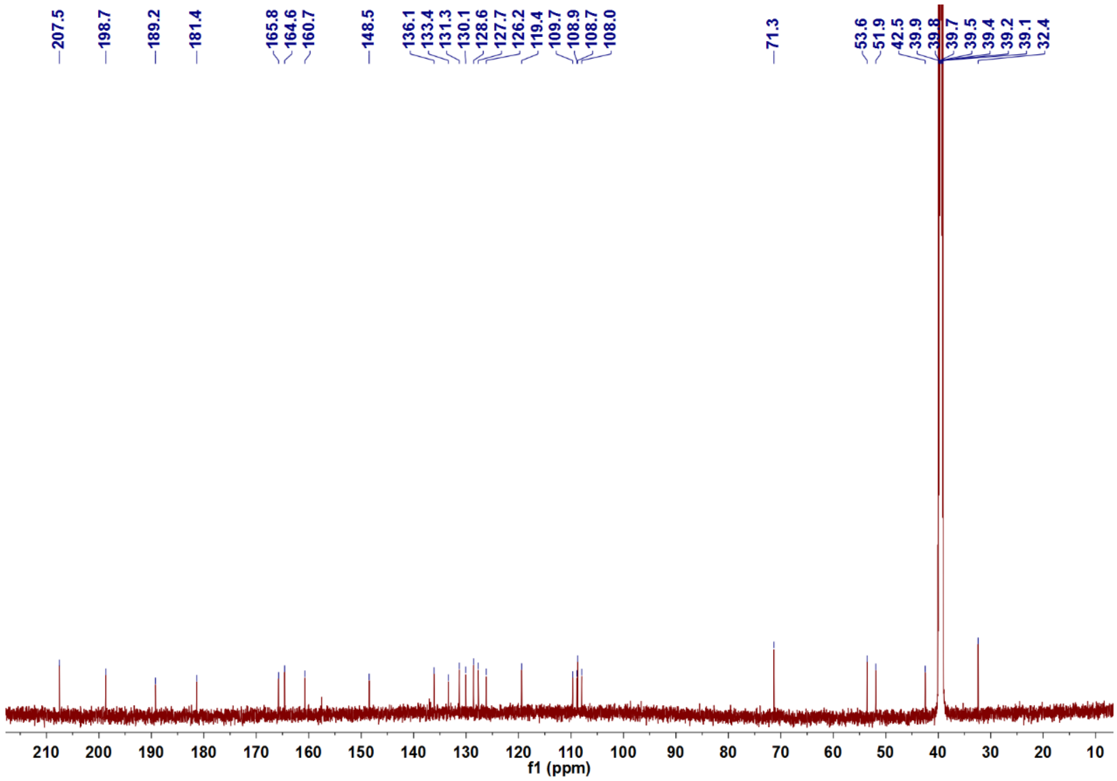


**Supplementary Fig. 23.** $^{13}$C NMR spectrum of **4** in DMSO-$d_6$.

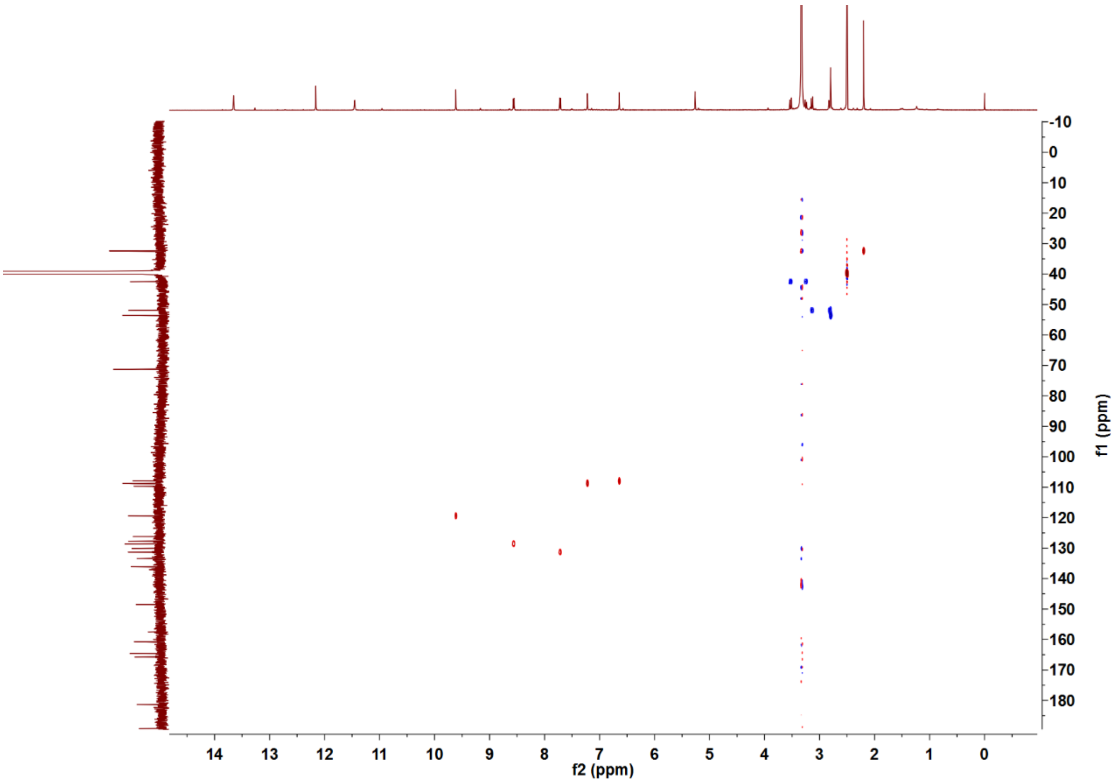


**Supplementary Fig. 24.** HSQC spectrum of **4** in DMSO-$d_6$.

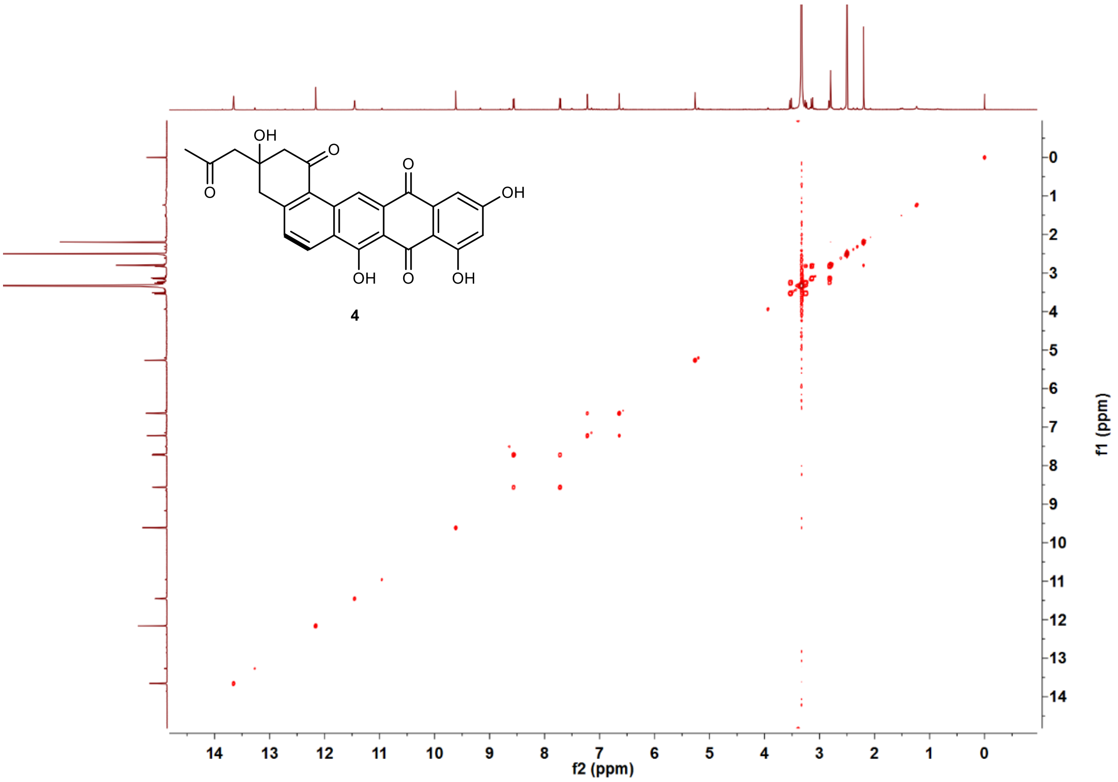


**Supplementary Fig. 25.** COSY spectrum of **4** in DMSO-$d_6$.

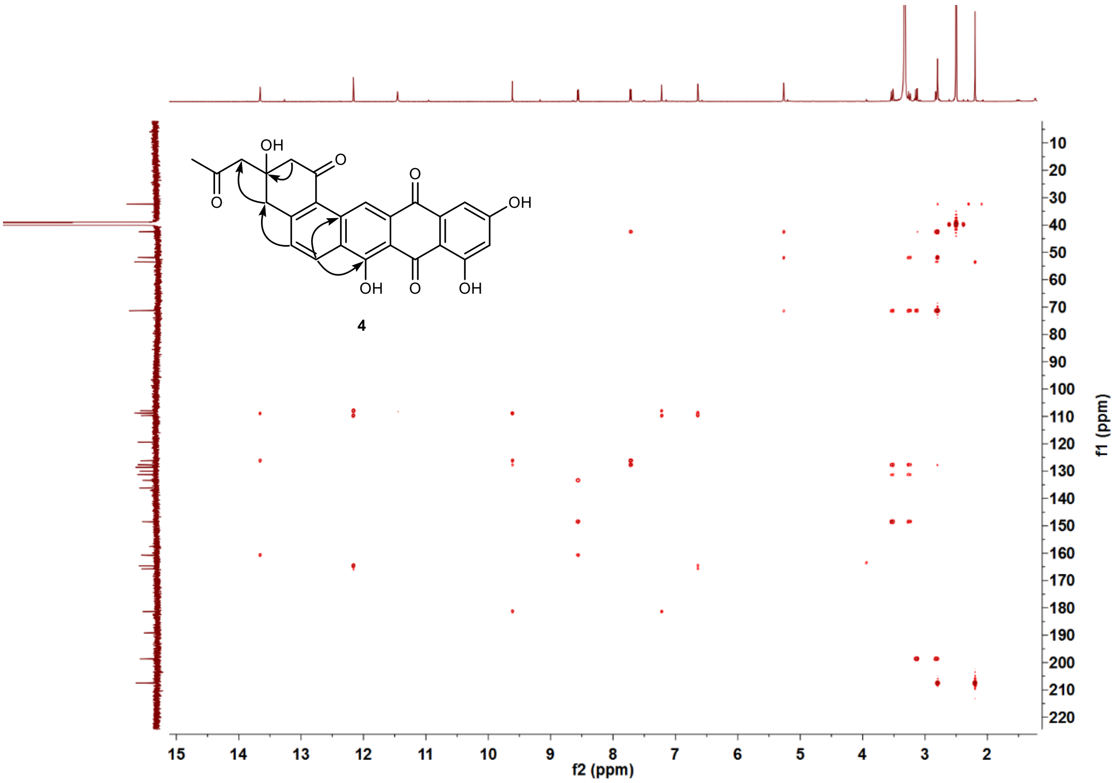


**Supplementary Fig. 26.** HMBC spectrum of **4** in DMSO-$d_6$.

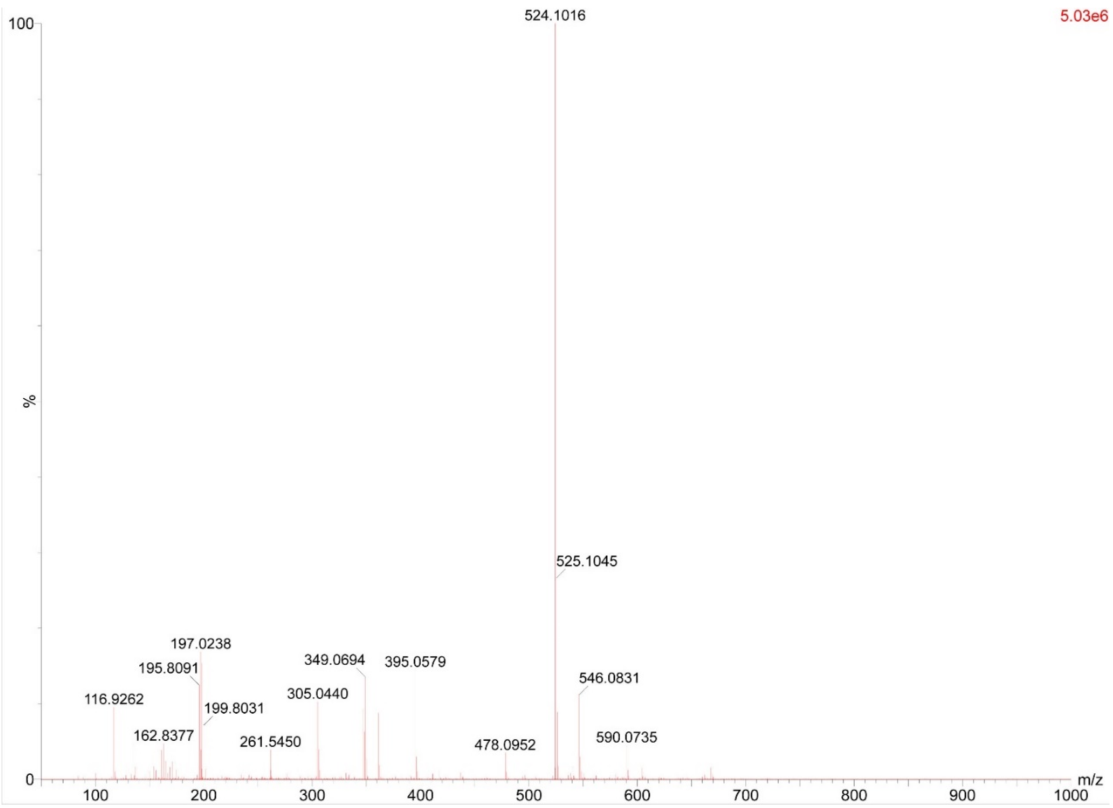


**Supplementary Fig. 27.** HR-ESI-MS spectrum of anthrapyrone A (**5**).

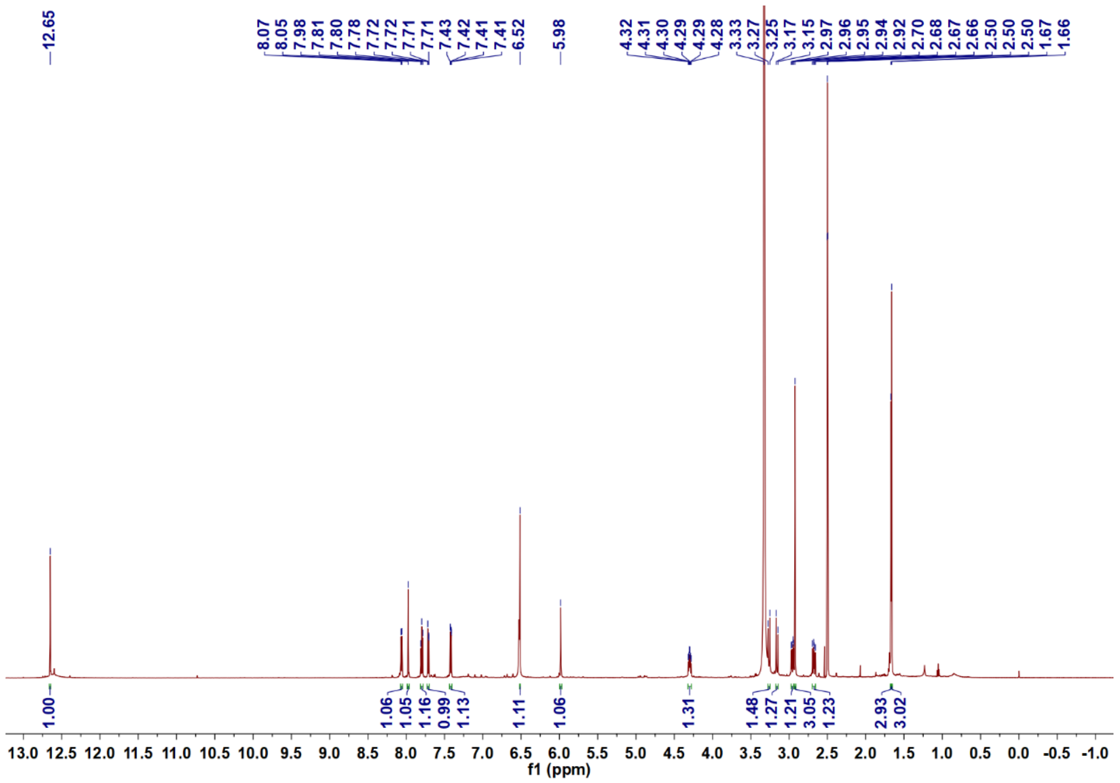

**Supplementary Fig. 28.** $^{1}$H NMR spectrum of anthrapyrone A (**5**) in DMSO-$d_6$.

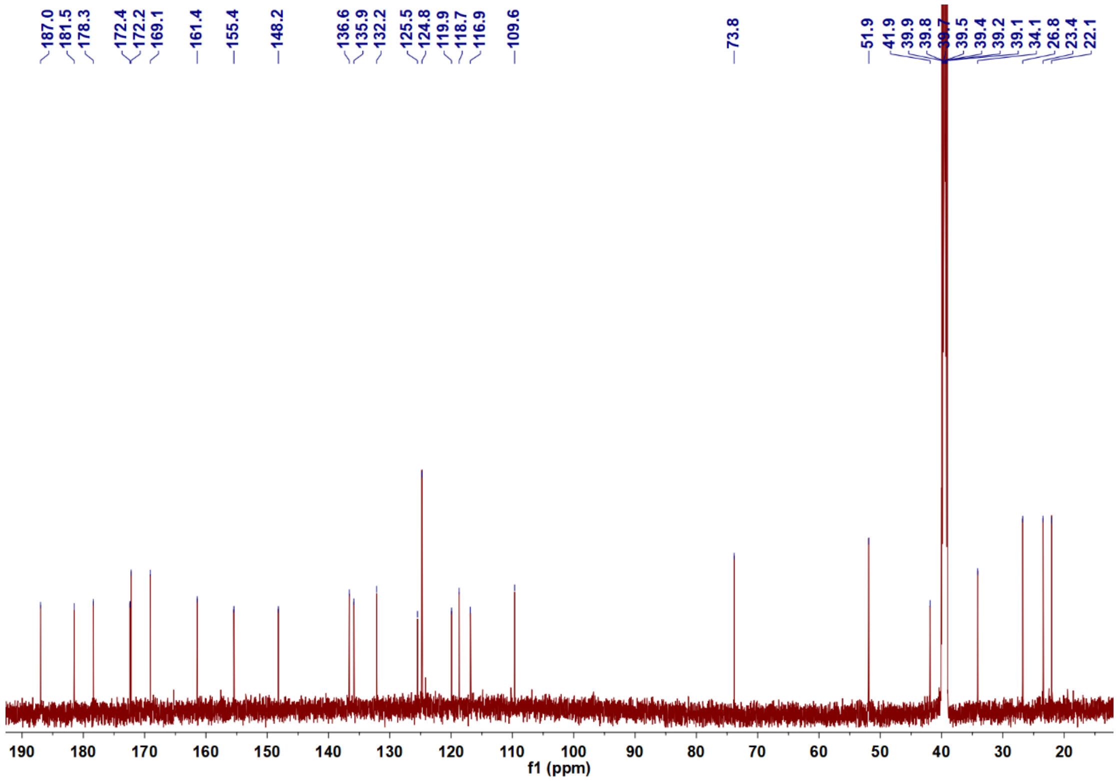

**Supplementary Fig. 29.** $^{13}$C NMR spectrum of anthrapyrone A (**5**) in DMSO-$d_6$.

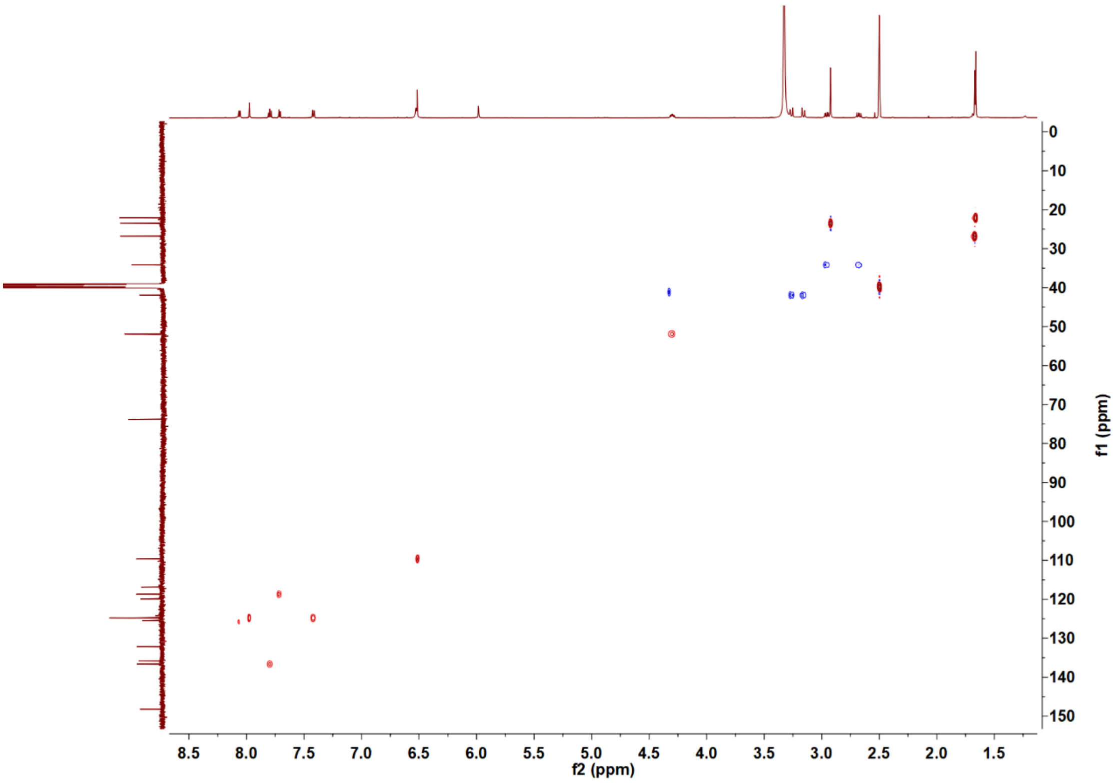


**Supplementary Fig. 30.** HSQC spectrum of anthrapyrone A (**5**) in DMSO-$d_6$.

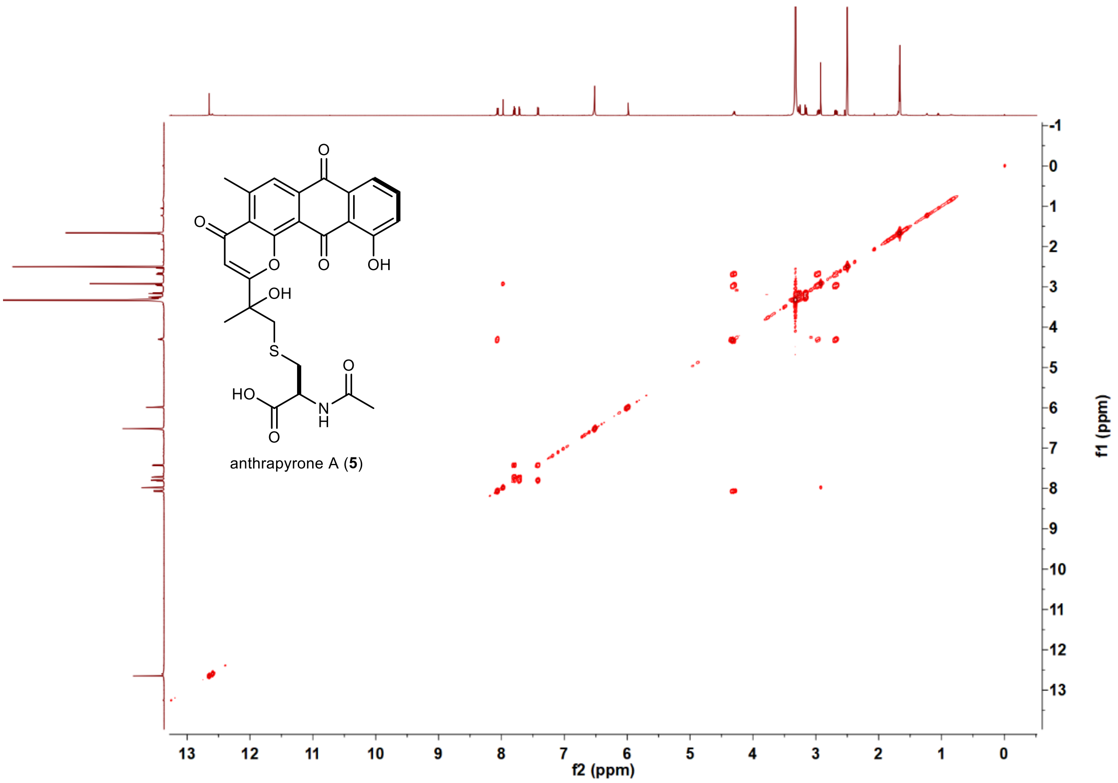


**Supplementary Fig. 31.** COSY spectrum of anthrapyrone A (**5**) in DMSO-$d_6$.

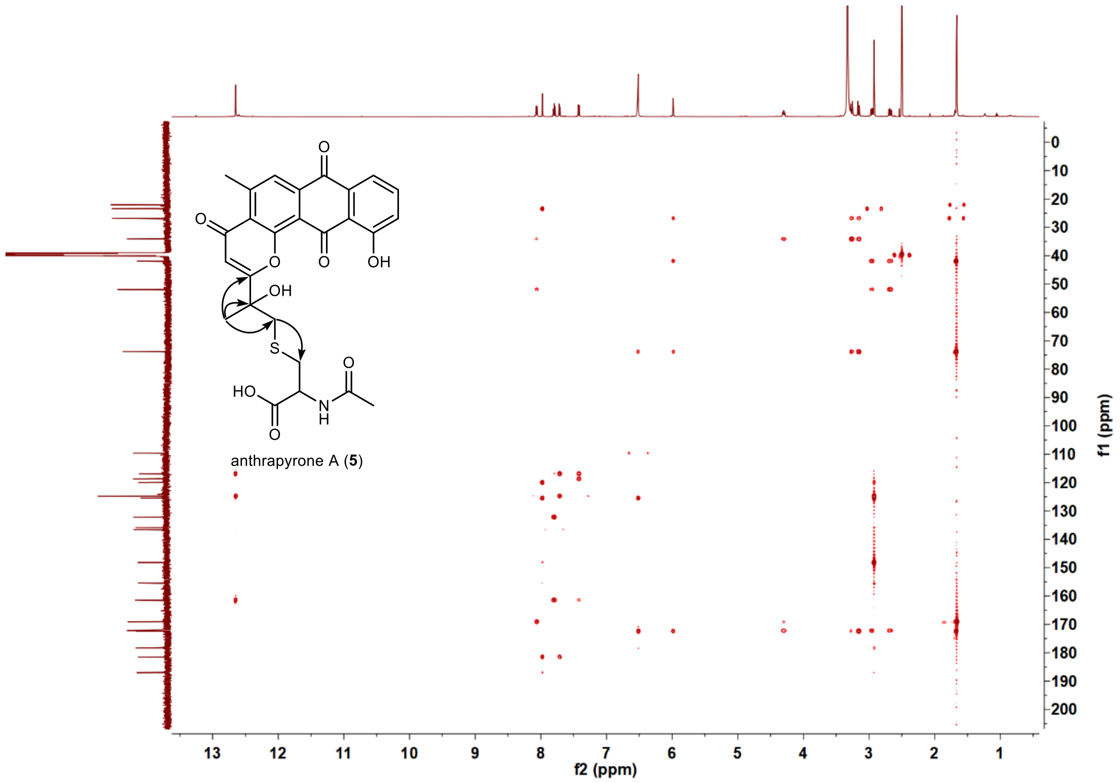


**Supplementary Fig. 32.** HMBC spectrum of anthrapyrone A (**5**) in DMSO-$d_6$.

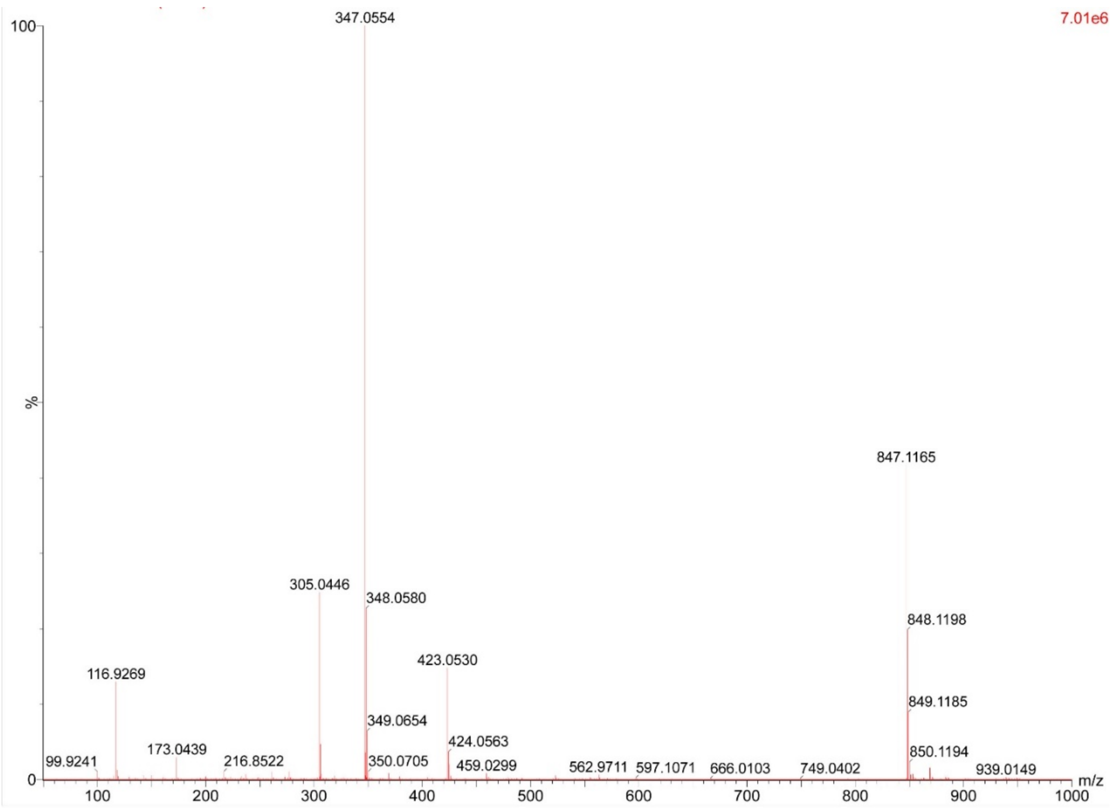


**Supplementary Fig. 33.** HR-ESI-MS spectrum of anthrapyrone B (**6**).

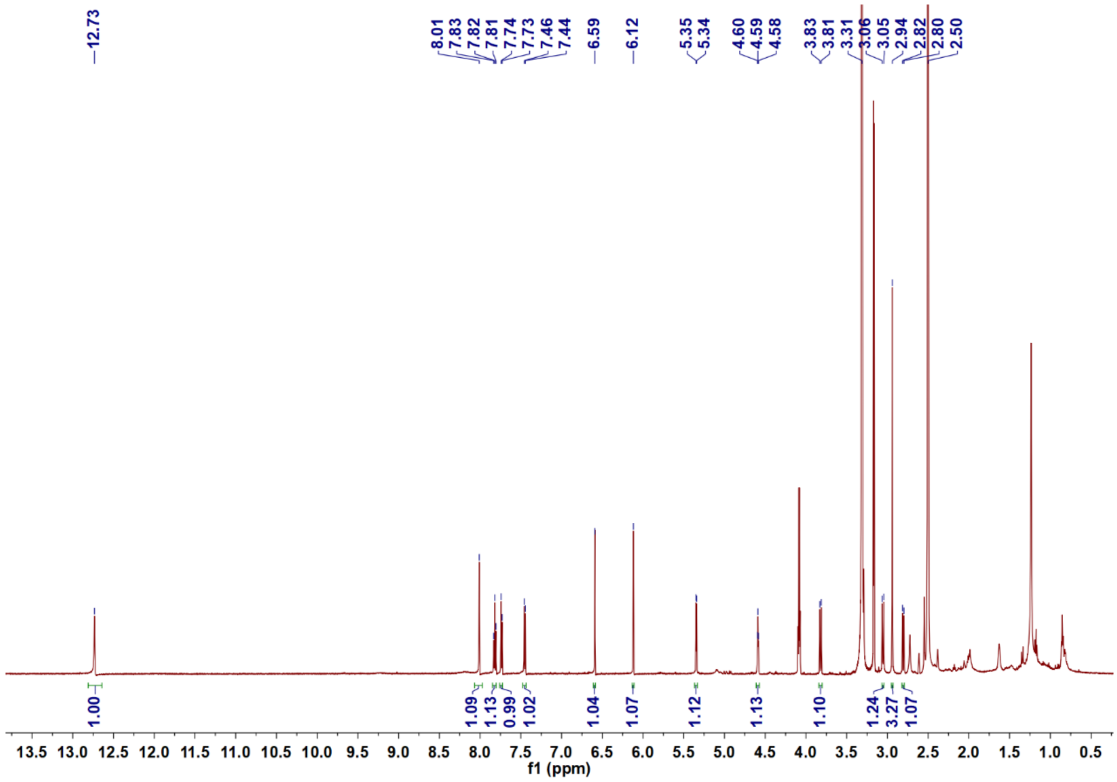


**Supplementary Fig. 34.** $^{1}$H NMR spectrum of anthrapyrone B (**6**) in DMSO-$d_6$.

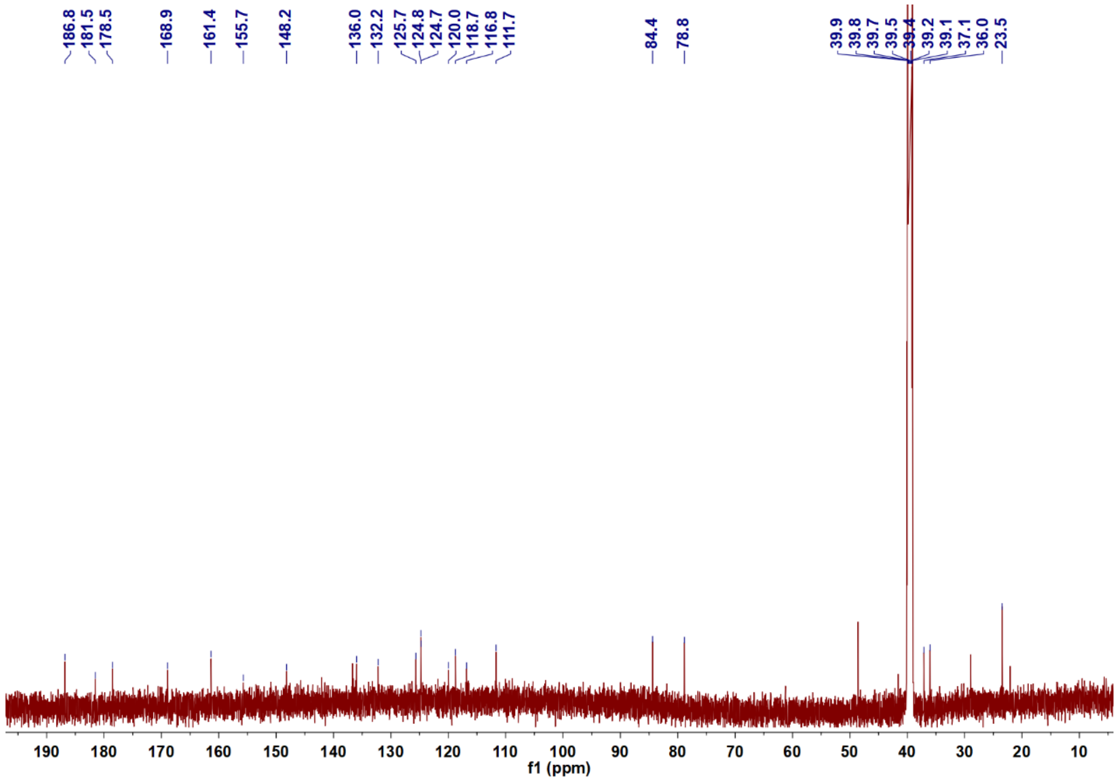


**Supplementary Fig. 35.** $^{13}$C NMR spectrum of anthrapyrone B (**6**) in DMSO-$d_6$.

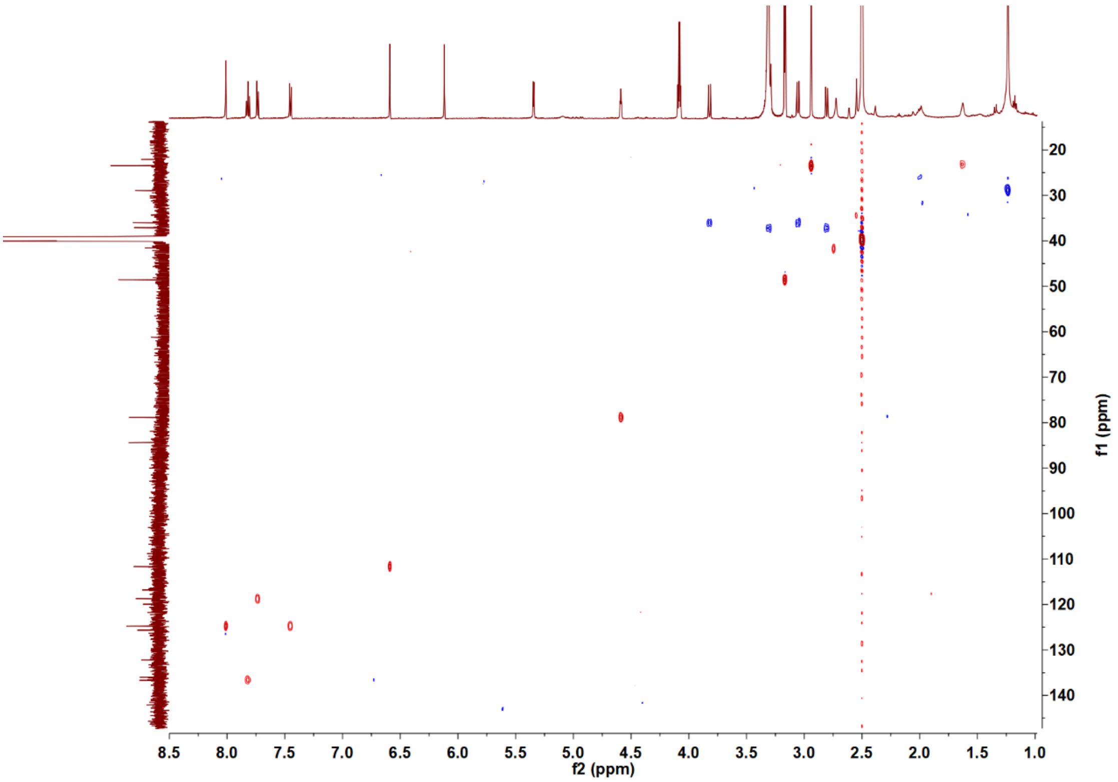


**Supplementary Fig. 36.** HSQC spectrum of anthrapyrone B (**6**) in DMSO-$d_6$.

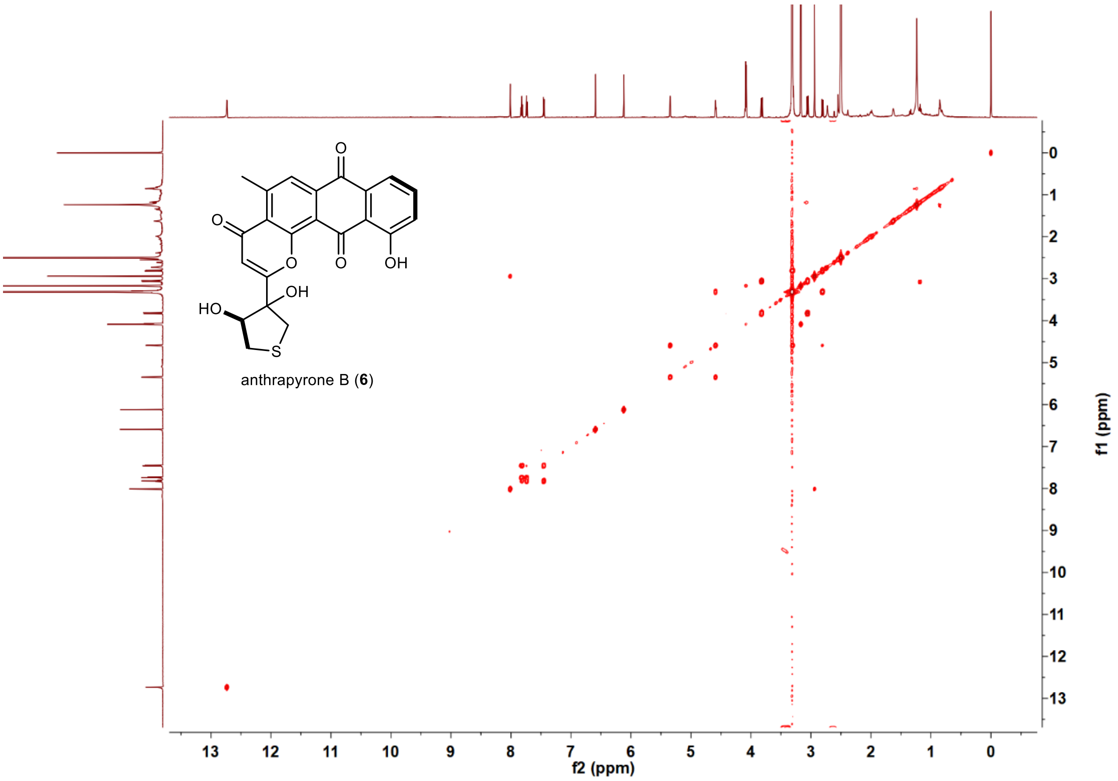


**Supplementary Fig. 37.** COSY spectrum of anthrapyrone B (**6**) in DMSO-$d_6$.

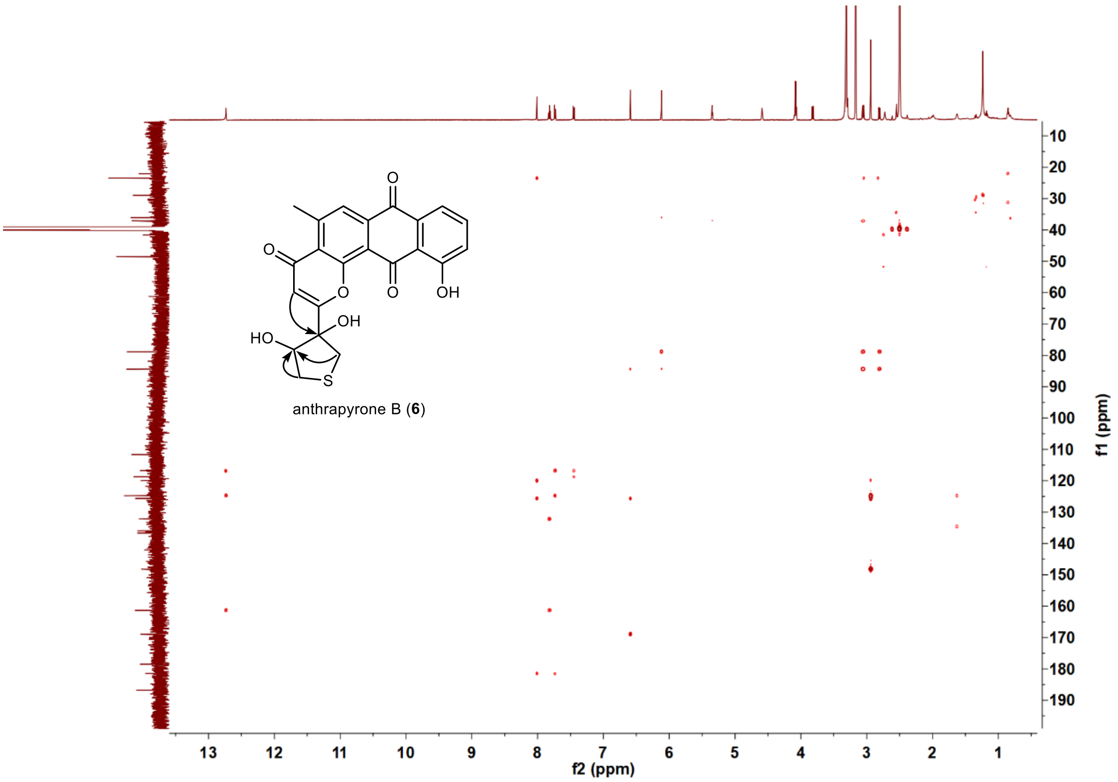


**Supplementary Fig. 38.** HMBC spectrum of anthrapyrone B (**6**) in DMSO-$d_6$.

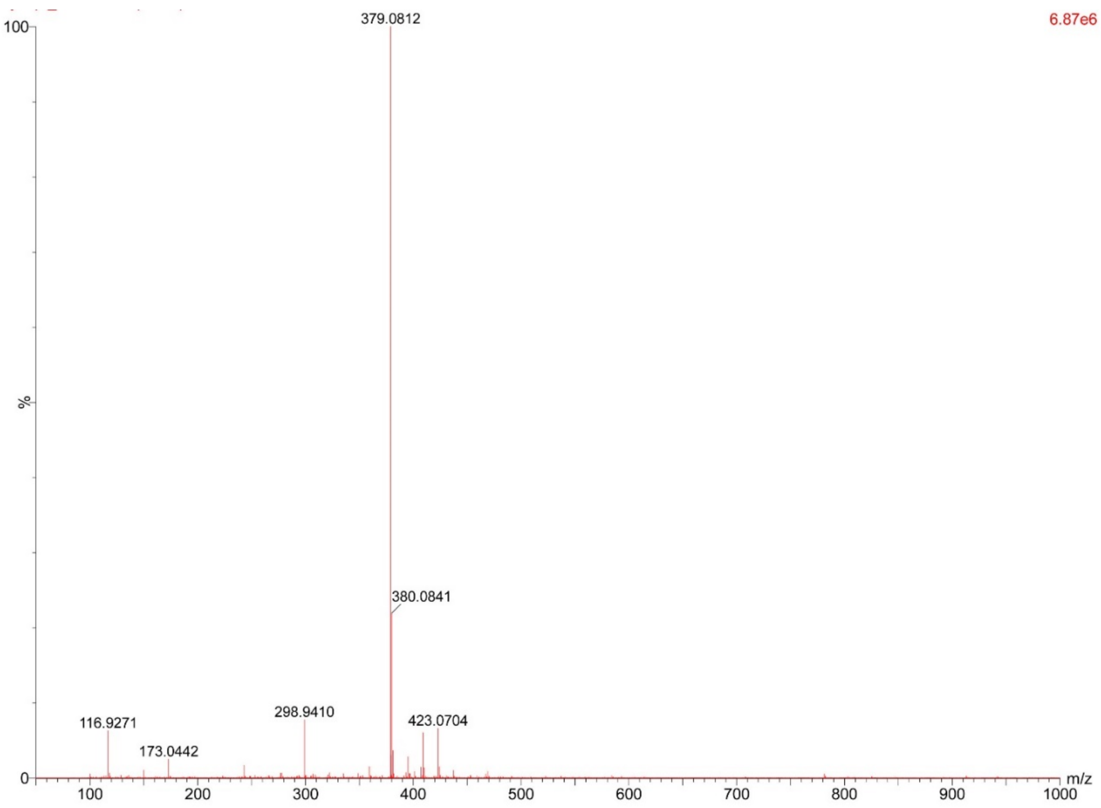


**Supplementary Fig. 39.** HR-ESI-MS spectrum of anthrapyrone C (**7**).

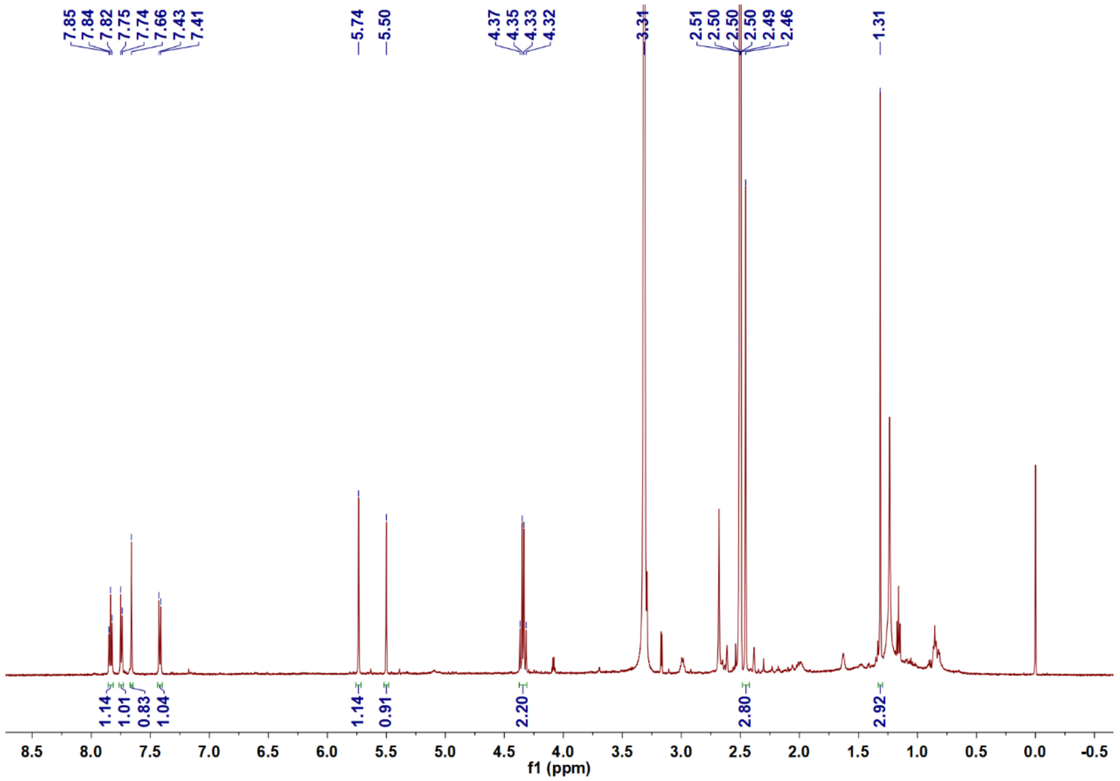


**Supplementary Fig. 40.** $^{1}$H NMR spectrum of anthrapyrone C (**7**) in DMSO-$d_6$.

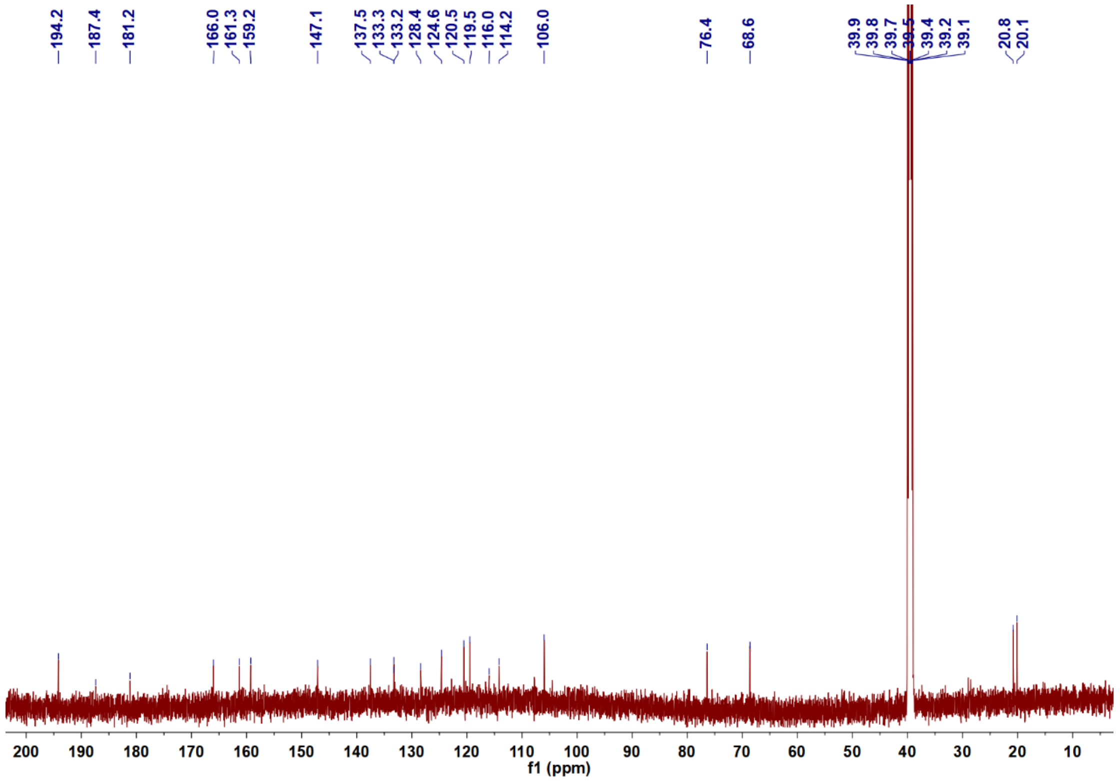


**Supplementary Fig. 41.** $^{13}$C NMR spectrum of anthrapyrone C (**7**) in DMSO-$d_6$.

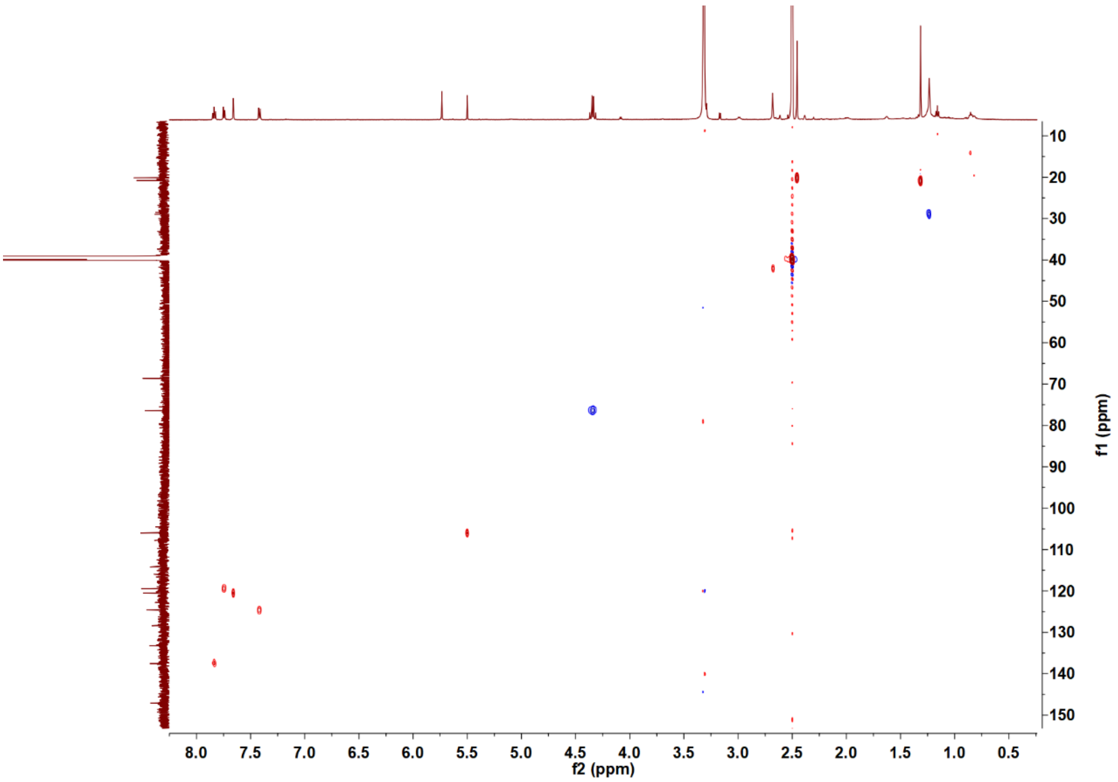


**Supplementary Fig. 42.** HSQC spectrum of anthrapyrone C (**7**) in DMSO-$d_6$.

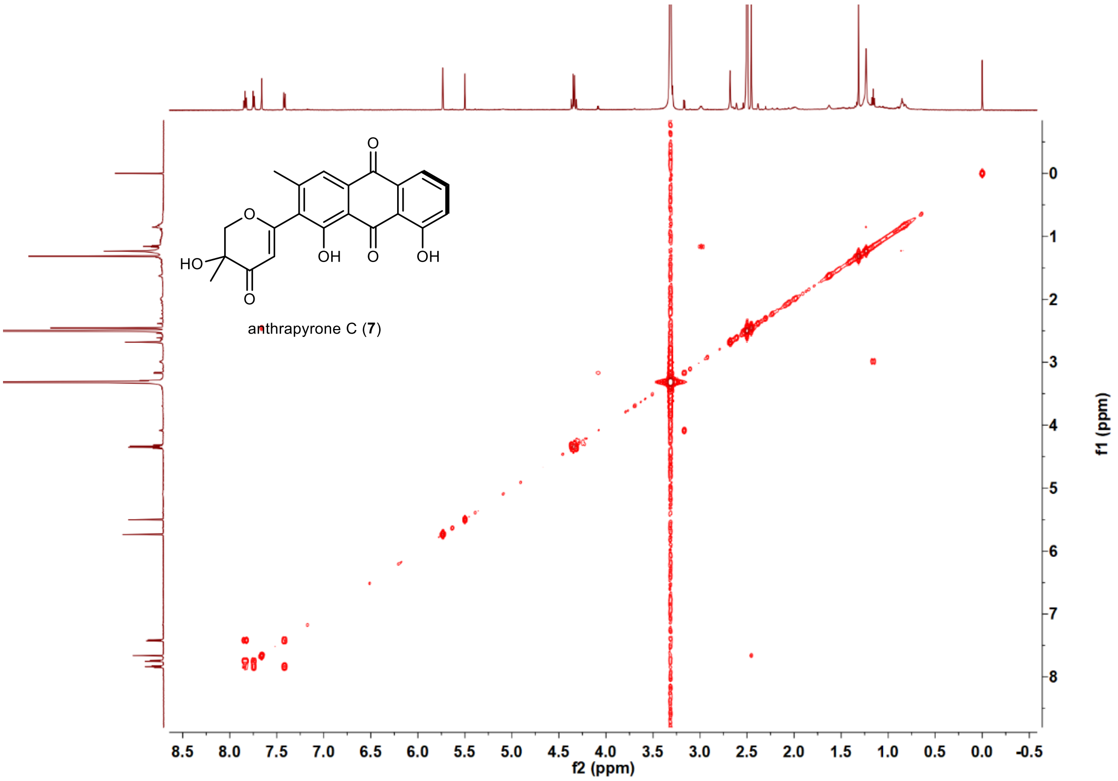


**Supplementary Fig. 43.** COSY spectrum of anthrapyrone C (**7**) in DMSO-$d_6$.

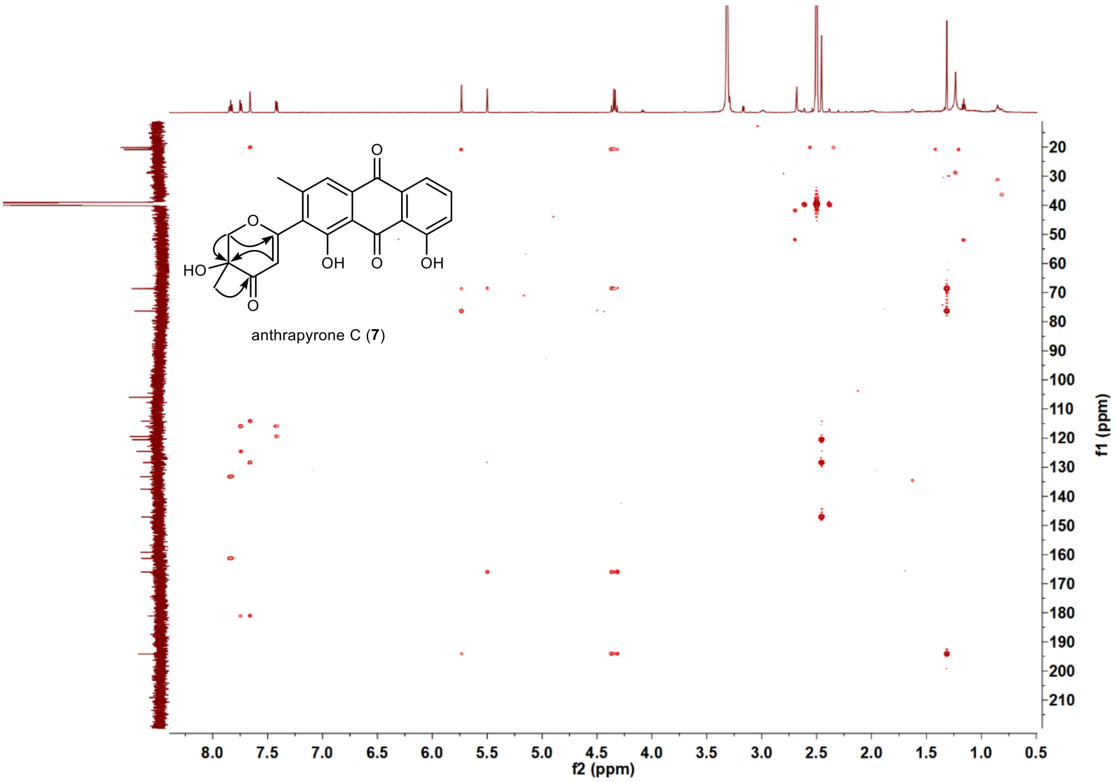


**Supplementary Fig. 44.** HMBC spectrum of anthrapyrone C (**7**) in DMSO-$d_6$.

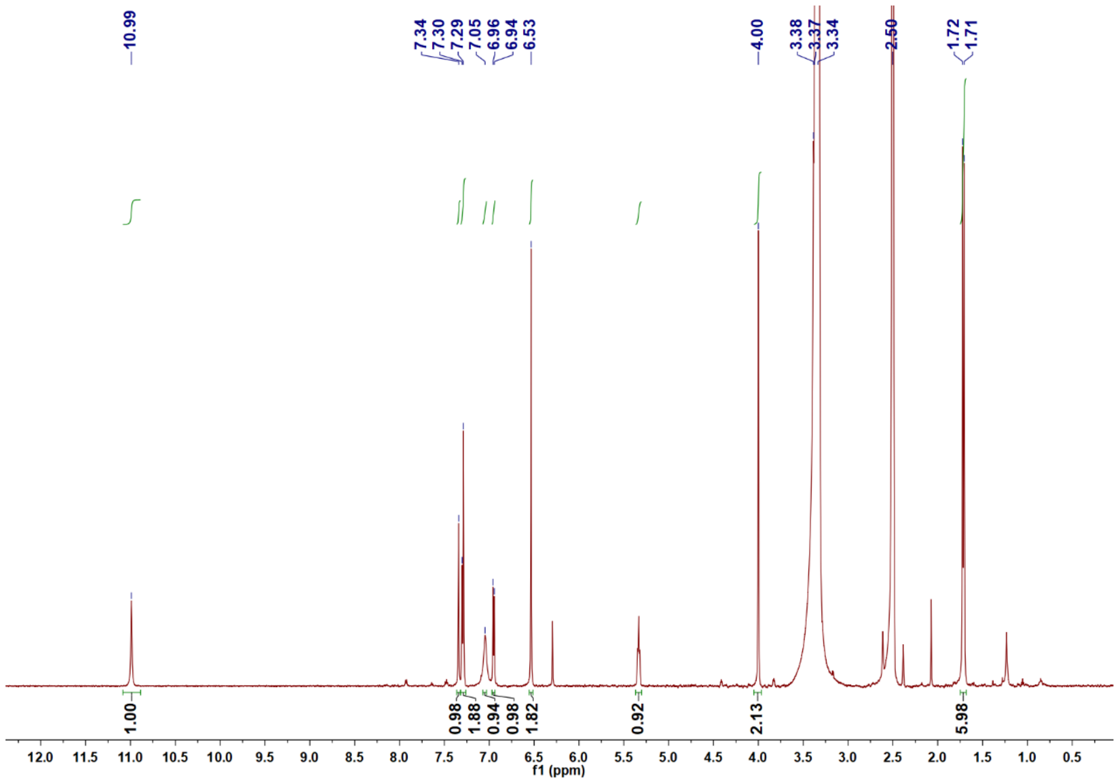


**Supplementary Fig. 45.** $^1$H NMR spectrum of 5-DMAIAN (**8**) in DMSO-$d_6$.

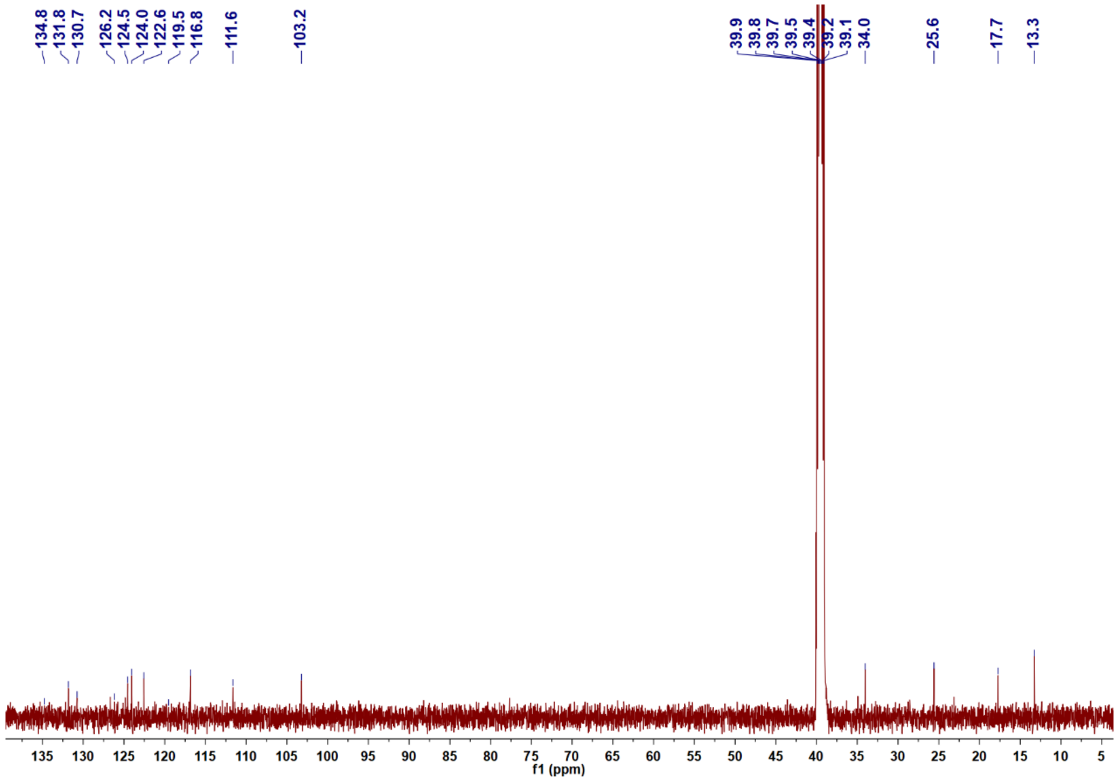


**Supplementary Fig. 46.** The $^{13}$C NMR spectrum of 5-DMAIAN (**8**) in DMSO-$d_6$.

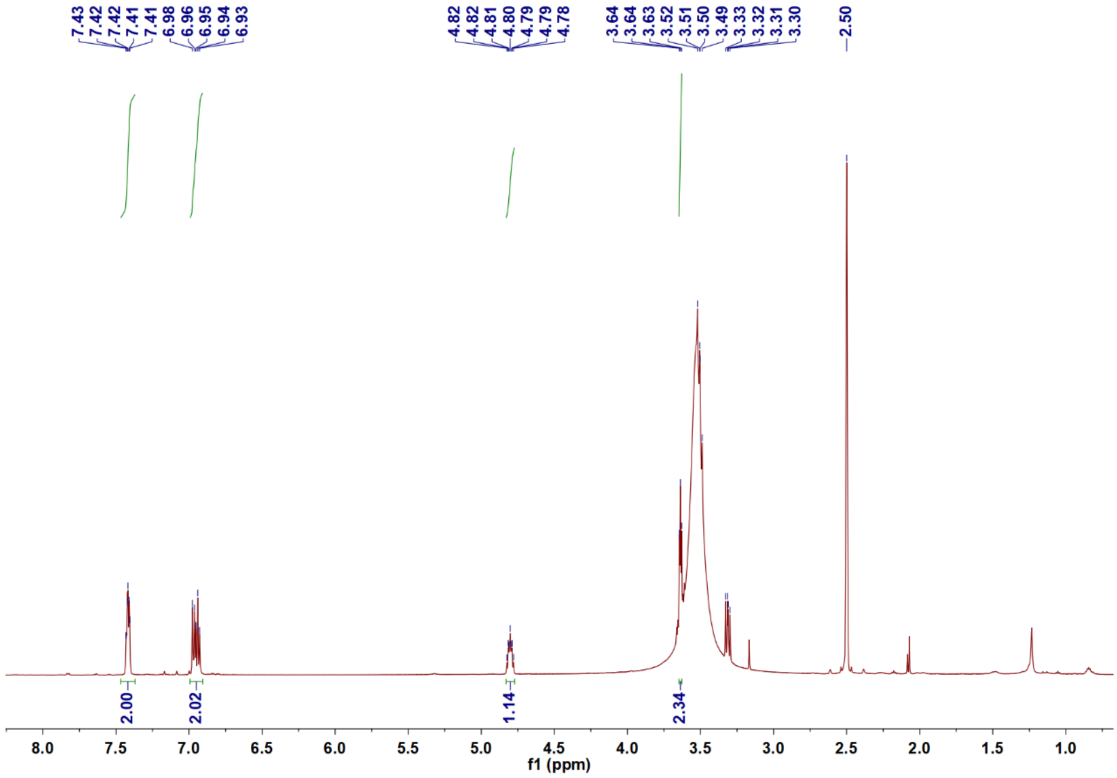


**Supplementary Fig. 47.** $^{1}$H NMR spectrum of aerugine (**9**) in DMSO-$d_6$.

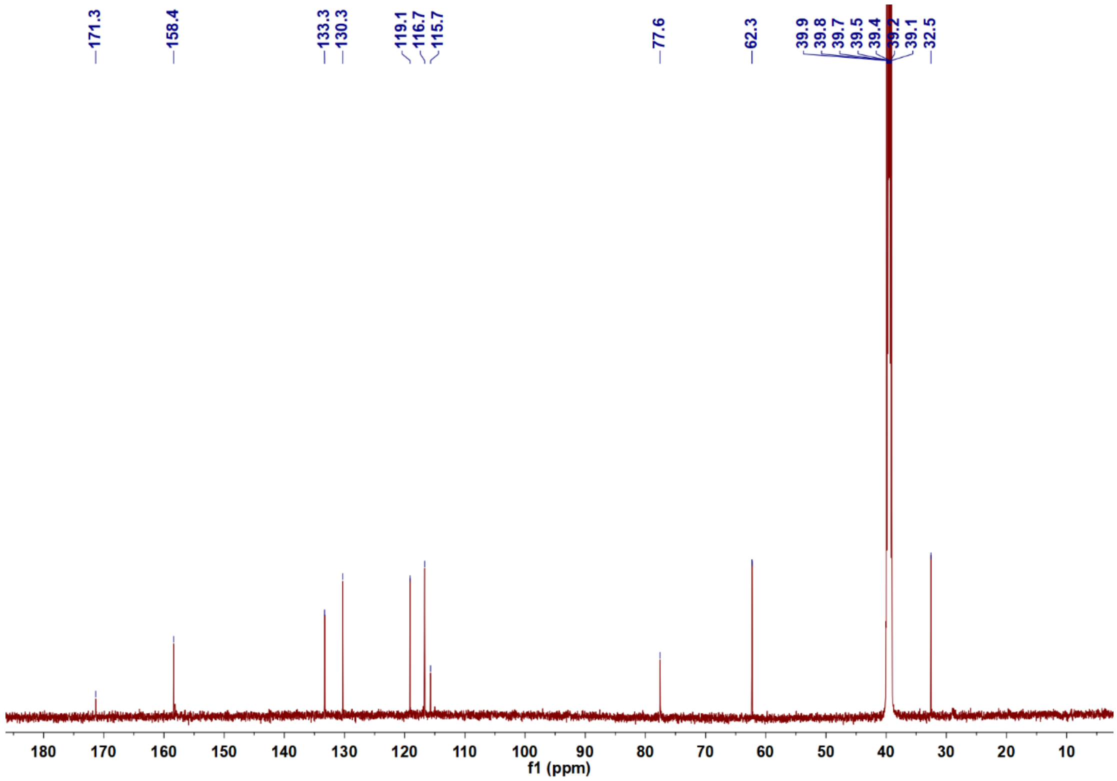


**Supplementary Fig. 48.** $^{13}$C NMR spectrum of aerugine (**9**) in DMSO-$d_6$.

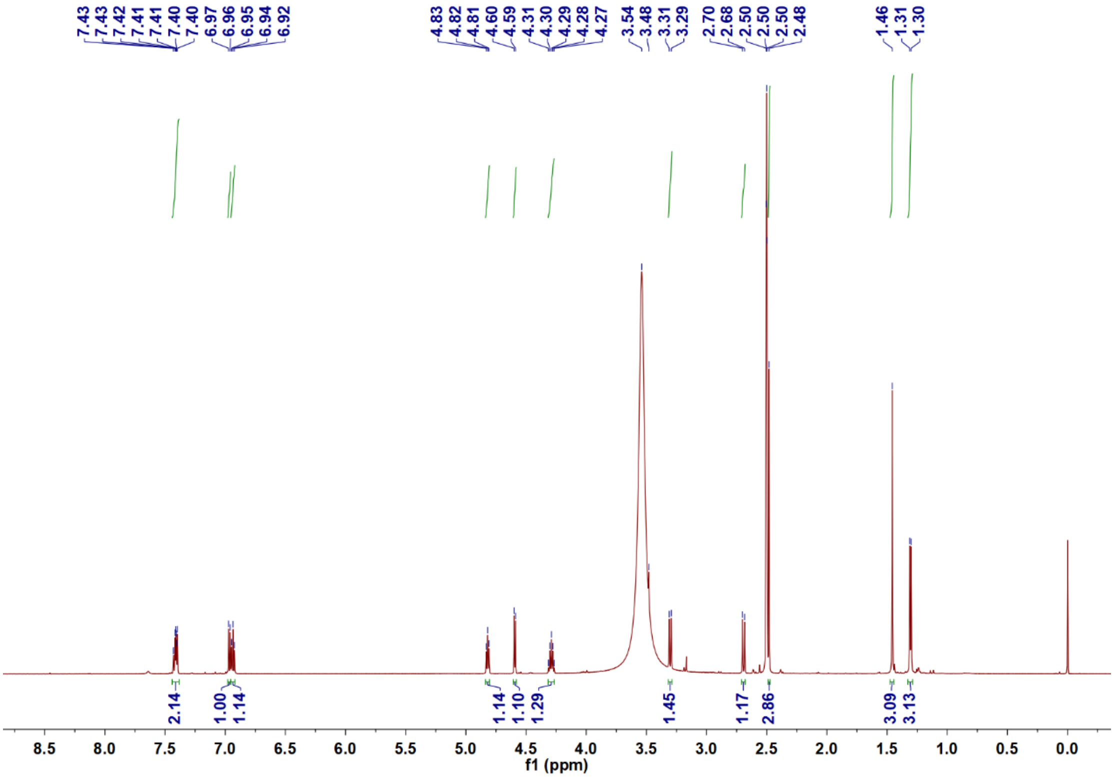


**Supplementary Fig. 49.** $^{1}$H NMR spectrum of watasemycin (**10**) in DMSO-$d_6$.

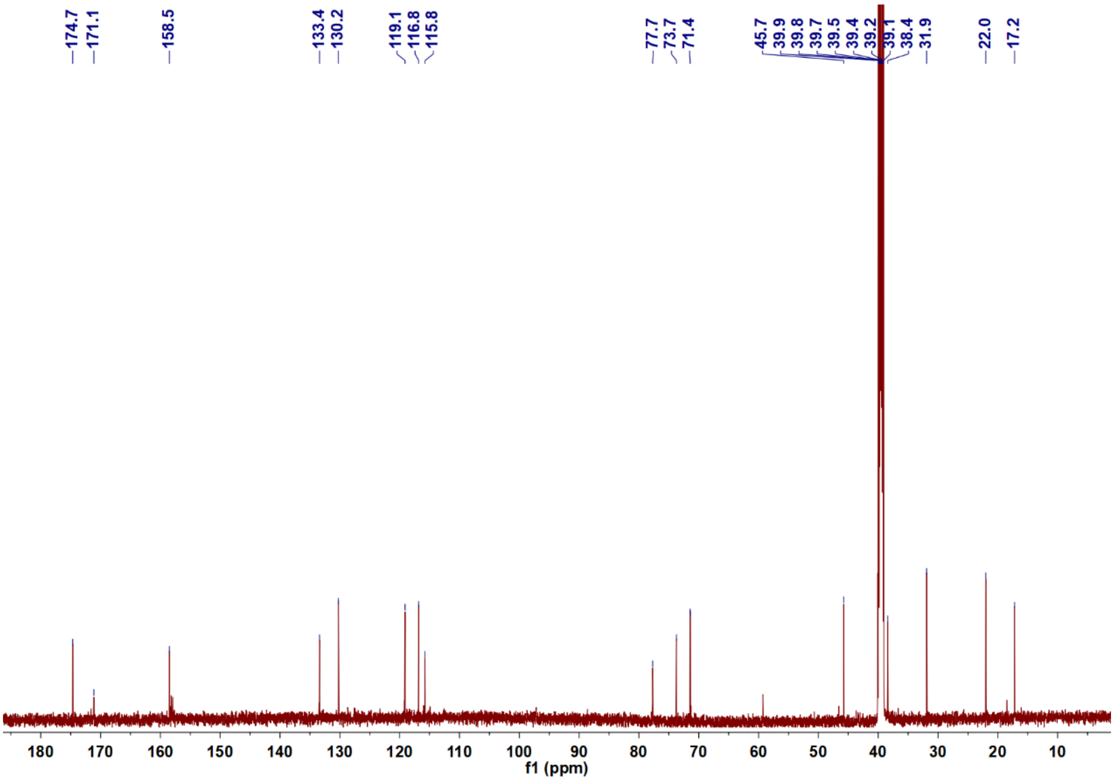


**Supplementary Fig. 50.** $^{13}$C NMR spectrum of watasemycin (**10**) in DMSO-$d_6$.

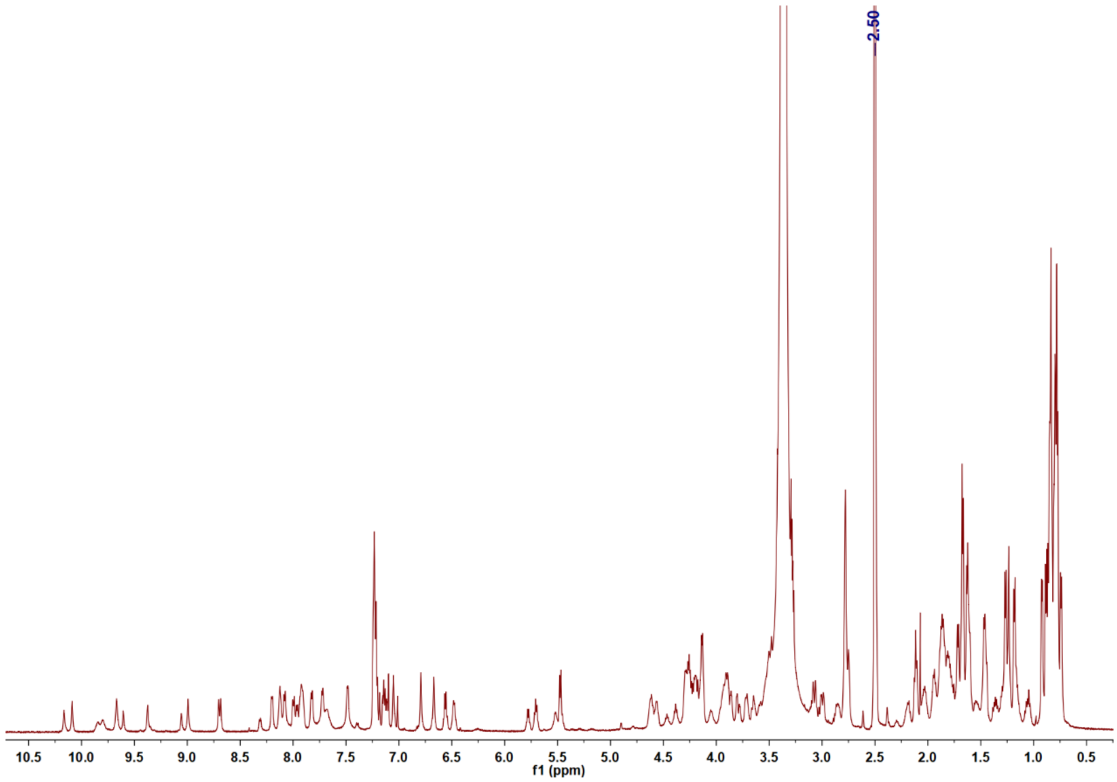


**Supplementary Fig. 51.** $^{1}$H NMR spectrum of cypemycin (**11**) in DMSO-$d_6$.

**Supplementary Table 5.** Summary of the results from the 12 BGCs produced detectable natural products.

| No. | Target BGC | Type | Size/bp | ResModel | HR-LC/MS |
|---|---|---|---|---|---|
| **1** | 16930_BGC #5 | Terpene | 15,198 | Exo beta-lactamase (class a) | 1 peak |
| **2** | 2700_BGC #9 | NRPS-like | 45,934 | DNA gyrase/topoisomerase IV | 1 peak |
| **3** | 2700_BGC #13 | NRPS-like | 31,634 | Other hits[a] | 1 peak |
| **4** | 5013_BGC #9 | T2PKS | 72,542 | Biotin_lipoyl: Biotin-requiring enzyme | 1 peak |
| **5** | 5013_BGC #14 | PKS/NRPS hybrid | 56,989 | Other hits[a] | 2 peaks |
| **6** | 5582_BGC #1 | RiPPS | 22,608 | ABC_efflux: ATP-binding cassette (ABC) antibiotic efflux pump | 5 peaks |
| **7** | 12374_BGC #1 | RiPPS | 16,491 | Other hits[a] | 1 peak |
| **8** | 24309_BGC #1 | T2PKS | 87,460 | acyl-CoA carboxylase subunit beta | 4 peaks |
| **9** | 12769_BGC #9 | T2PKS | 75,378 | MBL fold metallo-hydrolase | 3 peaks |
| **10** | 5635_BGC #6 | NRPS/T1PKS hybrid | 95,646 | ABC_efflux: ATP-binding cassette (ABC) antibiotic efflux pump | 2 peaks |
| **11** | 3061_BGC #8 | RiPPS | 36,274 | Other hits[a] | 1 peak |
| **12** | 3061_BGC #9 | NRPS | 43,318 | OTCace: Aspartate/ornithine carbamoyltransferase | 1 peak |

[a]Other hits: unknown resistance model but the duplication genes in BGCs.

**Supplementary Table 6.** $^1$H NMR (600 MHz) and $^{13}$C NMR (150 MHz) spectroscopic data for KS-619-1 (**1**) and **2** (*δ* in ppm).

| | KS-619-1 (**1**)[a] | | **2**[b] | |
|---|---|---|---|---|
| position | $\delta_C$, type | $\delta_H$, multi (*J* in Hz) | $\delta_C$, type | $\delta_H$, multi (*J* in Hz) |
| 1 | 164.8, C | | 163.6, C | |
| 2 | 116.9, C | | 118.5, C | |
| 3 | 139.4, C | | 140.9, C | |
| 4 | 119.7, CH | 6.29, s | 124.8, CH | 6.58, s |
| 4a | 140.5, C | | 139.8, C | |
| 5 | 28.5, $CH_2$ | 2.64, m | 38.3, $CH_2$ | 2.95, dd (3.6, 16.4)<br>3.09, dd (2.3, 16.4) |
| 6 | 20.4, $CH_2$ | 2.76, m | 59.8, CH | 5.40, m |
| 6a | 130.8, C | | 131.7, C | |
| 7 | 156.9, C | | 160.0, C | |
| 7a | 113.9, C | | 115.8, C | |
| 8 | 182.0, C | | 188.1, C | |
| 8a | 102.1, C | | 106.1, C | |
| 9 | 166.0, C | | 167.9, C | |
| 10 | 107.4, CH | 5.71, s | 110.3, CH | 6.18, d (2.3) |
| 11 | N, C | | 161.5, C | |
| 12 | 118.0, CH | 6.60, s | 117.3, CH | 7.03, d (2.3) |
| 12a | 134.4, C | | 136.3, C | |
| 13 | 184.1, C | | 185.7, C | |
| 13a | 130.2, C | | 133.9, C | |
| 14 | 118.7, CH | 8.97, s | 121.1, CH | 9.14, s |
| 14a | 139.6, C | | 140.8, C | |
| 14b | 118.7, C | | 119.9, C | |
| 15 | 50.2, $CH_2$ | 3.97, s | 51.3, $CH_2$ | 4.07, brs<br>4.11, brs |
| 16 | 205.4, C | | 210.5, C | |
| 17 | 29.8, $CH_3$ | 2.10, s | 30.0, $CH_3$ | 2.23, s |
| 18 | 171.9, C | | 175.7, C | |

[a]Measured in DMSO-$d_6$; [b]Measured in $CD_3OD$. N = no signal.

**Supplementary Table 7.** $^{1}H$ NMR (600 MHz) and $^{13}C$ NMR (150 MHz) spectroscopic data for compounds **3** and **4** in DMSO-$d_6$ ($\delta$ in ppm).

| | **3** | | **4** | |
|---|---|---|---|---|
| position | $\delta_C$, type | $\delta_H$, multi ($J$ in Hz) | $\delta_C$, type | $\delta_H$, multi ($J$ in Hz) |
| 1 | 155.9, C | | 198.7, C | |
| 2 | 116.2, C | 6.73, s | 51.9, $CH_2$ | 2.82, d (15.4) |
| | | | | 3.14, d (15.4) |
| 3 | 137.2, C | | 71.3, C | |
| 4 | 122.1, CH | 6.66, s | 42.5, $CH_2$ | 3.25, d (17.3) |
| | | | | 3.53, d (17.3) |
| 4a | 140.4, C | | 148.5, C | |
| 5 | 37.0, $CH_2$ | 2.83, d (2.3, 16.1) | 131.3, CH | 7.72, d (8.5) |
| | | 3.00, d (2.0, 16.1) | | |
| 6 | 57.0, CH | 5.22, m | 128.6, CH | 8.56, d (8.5) |
| 6a | 131.7, C | | 126.2, C | |
| 7 | 158.7, C | | 160.7, C | |
| 7a | 113.2, C | | 108.9, C | |
| 8 | 189.5, C | | 189.2, C | |
| 8a | 109.3, C | | 109.7, C | |
| 9 | 164.4, C | | 164.6, C | |
| 10 | 107.9, CH | 6.63, d (2.3) | 108.0, CH | 6.64, d (2.1) |
| 11 | 165.6, C | | 165.8, C | |
| 12 | 108.8, CH | 7.18, d (2.3) | 108.7, CH | 7.22, d (2.1) |
| 12a | 135.4, C | | 136.1, C | |
| 13 | 181.6, C | | 181.4, C | |
| 13a | 131.5, C | | 130.1, C | |
| 14 | 120.0, CH | 8.95, s | 119.4, CH | 9.61, s |
| 14a | 137.6, C | | 133.4, C | |
| 14b | 116.9, C | | 127.7, C | |
| 15 | 49.4, $CH_2$ | 3.71, s | 53.6, $CH_2$ | 2.80, s |
| 16 | 205.7, C | | 207.5, C | |
| 17 | 29.6, $CH_3$ | 2.16, s | 32.4, $CH_3$ | 2.20, s |
| 1-OH | | 10.30, s | | |
| 3-OH | | | | 5.26, s |
| 6-OH | | 5.03, d (3.1) | | |
| 7-OH | | 12.59, s | | 13.65, s |
| 9-OH | | 12.14, s | | 12.16, s |
| 11-OH | | 11.40, s | | 11.45, s |

**Supplementary Table 8.** $^{1}$H NMR (600 MHz) and $^{13}$C NMR (150 MHz) spectroscopic data for anthrapyrones A (**5**) and B (**6**) in DMSO-$d_6$ ($\delta$ in ppm).

| | **5** | | **6** | |
|---|---|---|---|---|
| position | $\delta_C$, type | $\delta_H$, multi ($J$ in Hz) | $\delta_C$, type | $\delta_H$, multi ($J$ in Hz) |
| 1 | 161.4, C | | 161.4, C | |
| 2 | 124.8, CH | 7.42, dd (1.0, 8.3) | 124.8, CH | 7.45, d (8.3) |
| 3 | 136.6, CH | 7.80, dd (7.4, 8.3) | 136.7, CH | 7.82, dd (7.4, 8.3) |
| 4 | 118.7, CH | 7.71, dd (1.0, 7.4) | 118.7, CH | 7.74, d (7.4) |
| 4a | 132.2, C | | 132.2, C | |
| 5 | 124.8, CH | 7.98, s | 124.7, CH | 8.01, s |
| 6 | 148.2, C | | 148.2, C | |
| 7 | 125.5, C | | 125.7, C | |
| 8 | 155.4, C | | 155.7, C | |
| 8a | 119.9, C | | 120.0, C | |
| 9 | 187.0, C | | 186.8, C | |
| 9a | 116.9, C | | 116.8, C | |
| 10 | 181.5, C | | 181.5, C | |
| 10a | 135.9, C | | 136.0, C | |
| 11 | 178.3, C | | 178.5, C | |
| 12 | 109.4, CH | 6.52, s | 111.7, CH | 6.59, s |
| 13 | 172.4, C | | 168.9, C | |
| 14 | 73.8, C | | 84.4, C | |
| 15 | 41.9, $CH_2$ | 3.16, d (13.6) | 78.8, CH | 4.59, m |
| | | 3.26, d (13.6) | | |
| 16 | 26.8, $CH_3$ | 1.67, s | 37.1, $CH_2$ | 2.81, d (10.7) |
| | | | | 3.31, overlapped |
| 17 | 23.4, $CH_3$ | 2.92, s | 36.0, $CH_2$ | 3.05, d (11.1) |
| | | | | 3.82, d (11.1) |
| 18 | | | 23.5, $CH_3$ | 2.94, s |
| 1′ | 34.1, $CH_2$ | 2.68, dd (4.9, 13.6) | | |
| | | 2.96, dd (9.0, 13.6) | | |
| 2′ | 51.9, CH | 4.30, m | | |
| 3′ | 172.2, C | | | |
| 4′ | 169.1, C | | | |
| 5′ | 22.1, $CH_3$ | 1.66, s | | |
| 1-OH | | 12.65, s | | 12.73, s |
| 14-OH | | 5.98, s | | 6.12, s |
| 15-OH | | | | 5.35, d (4.3) |
| 2′-NH | | 8.06, d (7.8) | | |

**Supplementary Table 9.** $^{1}H$ NMR (600 MHz) and $^{13}C$ NMR (150 MHz) spectroscopic data for anthrapyrone C (**7**) in DMSO-$d_6$ ($\delta$ in ppm).

| | **7** | |
|---|---|---|
| position | $\delta_C$, type | $\delta_H$, multi ($J$ in Hz) |
| 1 | 161.3, C | |
| 2 | 124.6, CH | 7.42, d (8.3) |
| 3 | 137.5, CH | 7.84, dd (7.6, 8.3) |
| 4 | 119.5, CH | 7.74 d (7.6) |
| 4a | 133.2, C | |
| 5 | 120.5, CH | 7.66, s |
| 6 | 147.1, C | |
| 7 | 128.4, C | |
| 8 | 159.2, C | |
| 8a | 114.2, C | |
| 9 | 187.4, C | |
| 9a | 116.0, C | |
| 10 | 181.2, C | |
| 10a | 133.3, C | |
| 11 | 166.0, C | |
| 12 | 106.0, CH | 5.50, s |
| 13 | 194.2, C | |
| 14 | 68.6, C | |
| 15 | 76.4, $CH_2$ | 4.34, d (11.5) |
| 16 | 20.8, $CH_3$ | 1.31, s |
| 17 | 20.1, $CH_3$ | 2.46, s |
| 14-OH | | 5.74, s |
